# Unexpected Activities of CYP152 Peroxygenases towards Non-carboxylic Substrates Reveal Novel Substrate Recognition Mechanism and Catalytic Versatility

**DOI:** 10.1101/2024.12.20.629671

**Authors:** Yuanyuan Jiang, Piqian Gong, Zhong Li, Yuxuan Li, Binju Wang, Wei Peng, Xiang Gao, Shengying Li

## Abstract

Exploring and exploiting the catalytic promiscuity of enzymes is a central topic and captivating challenge in enzymology. CYP152 peroxygenases are attractive biocatalysts for diverse reactions under mild conditions using H_2_O_2_ as cofactor. However, their substrate scope is limited by a carboxyl group required for acid-base catalysis, following the well-accepted principle that heme-dependent H_2_O_2_-utilizing enzymes employ a carboxyl group within their active sites to facilitate H_2_O_2_ activation. Herein, we reveal for the first time that several CYP152 family members can directly degrade various aromatic pollutants without any carboxyl group, exhibiting novel aromatic hydroxylation and dehalogenation activities. Through crystal structure analysis, isotope tracing experiments, and QM/MM calculations, we elucidate that the phenolic hydroxyl group activated by electron-withdrawing substituent(s) functionally replaces the carboxyl group, forming hydrogen bonds with the conserved arginine leading to Compound I formation. The oxygen atom of the newly formed hydroxyl group originates from water, bypassing the conventional oxygen rebound step. These findings provide first insights into the mechanisms of P450 peroxygenases towards non-carboxylic substrates, expanding our knowledge of biological C-H activation and C-halogen bond cleavage beyond canonical P450 reactions. This discovery holds immense potential for harnessing these enzymes in innovative strategies for industrial biocatalysis and environmental remediation.

## Introduction

The capability of a single enzyme to catalyze diverse and mechanistically distinct reactions is pivotal in enabling its host organism’s past adaptation to external stimuli and continues to be observed in contemporary enzymes._[1–4]_ Given the importance of biocatalytic promiscuity in nature, exploring and engineering enzymes for improved catalytic promiscuity holds the key to broadening the realm of enzymatic synthesis_[5,6]_ and hence the promise for both biomedical and biotechnological applications._[7,8]_ This endeavor can also spur the advances in fine-tuning existing enzymatic activities, and even more profoundly, facilitate the discovery and development of novel catalytic abilities._[9]_

Cytochrome P450 monooxygenases (P450s or CYPs) are versatile biocatalysts that excel in oxyfunctionalization of diverse substrates with different kinds of C−H bonds._[10–15]_ Nonetheless, practical utilization of P450s *in vitro* remains constrained due to their pronounced reliance on NAD(P)H as electron supplier and a sophisticated network of electron transport proteins (*i.e.*, redox partners) for the catalytically required O_2_ activation._[16,17]_ In contrast, P450 peroxygenases stand out as appealing biocatalysts for selective oxidation of organic molecules under mild conditions with hydrogen peroxide as the sole oxygen and electron source._[18–20]_ The so-called peroxide shunt pathway (Figure S1) offers a more straightforward and cost-effective route to bypassing the intricate electron-delivering pathway in normal P450 catalysis. Consequently, the unique catalytic properties of this class of H_2_O_2_-utilizing enzymes have ignited intense interests in recent years, propelling the expansion of their catalytic repertoire and fostering their potential of applications across various disciplines._[21–24]_

In comparison, a vast majority of native P450 monooxygenases lack an acid-base pair of amino acid residues situated at the distal position of the heme reactive center, which is a common principle among heme-dependent H_2_O_2_-utilizing enzymes (*e.g.*, peroxidases, peroxygenases) that activate hydrogen peroxide to form the highly reactive species Compound I (Cpd I)._[19,25–27]_ Of note, a group of P450s (*e.g.*, CYP152 peroxygenases) are capable of using H_2_O_2_ through a substrate-assisted reaction mechanism._[18,28]_ The structures of CYP152 peroxygenase in complex with fatty acid substrate have shed molecular insight into the carboxyl group (of fatty acid) assisted substrate recognition and H_2_O_2_-activation with the conserved arginine residue (*e.g.*, R242 for P450_BSβ_) and unique catalytic mechanism (Figure 1A)._[29–31]_ The representative examples of CYP152 peroxygenases, possessing the ability to catalyze the hydroxylation or decarboxylation of fatty acids with H_2_O_2_, primarily include the fatty acid hydroxylases P450_SPα_ and P450_BSβ_, and fatty acid decarboxylase OleT_JE_ (Figures 1B, S2)._[32–44]_ Despite well-expanded substrate scope and notable enhancement in activity diversity,_[22,45–50]_ it is widely accepted that CYP152 peroxygenases are unable to use H_2_O_2_ to initiate their reaction in absence of the specific enzyme-substrate interaction via the guanidyl-carboxyl pairing. The carboxylic substrates are deemed to serve dual roles as both initiators and activators of the reaction._[19]_ Nonetheless, inspired by the salt bridge formation between the substrate carboxyl group and R242, P450_BSβ_ was enabled to oxidize non-carboxylic substrates (*e.g.*, styrene, ethylbenzene) in the presence of a decoy molecule that mimics the fatty acid structure in the active site (Figure 1B)._[51]_ Similar to decoy molecules, the substrate scope of P450_SPα_ was expanded by introducing a carboxylate group of glutamic acid into the active site nearing heme; the resulted A245E mutant achieved styrene epoxidation._[52]_ These examples showcase successful broadening of the substrate scope of CYP152 peroxygenases through introduction of an acid-base catalyst in the active site by amino acid substitution or the use of decoy molecules. To the best of our knowledge, the catalytic repertoire of CYP152 peroxygenases has been limited to the reactions relying on the key carboxyl group in the active site to facilitate H_2_O_2_ activation, thus posing constrains on exploring the diversity of substrate types and catalytic potentials, particularly in the uncharted area of non-carboxylic substrates.

**Figure 1.**
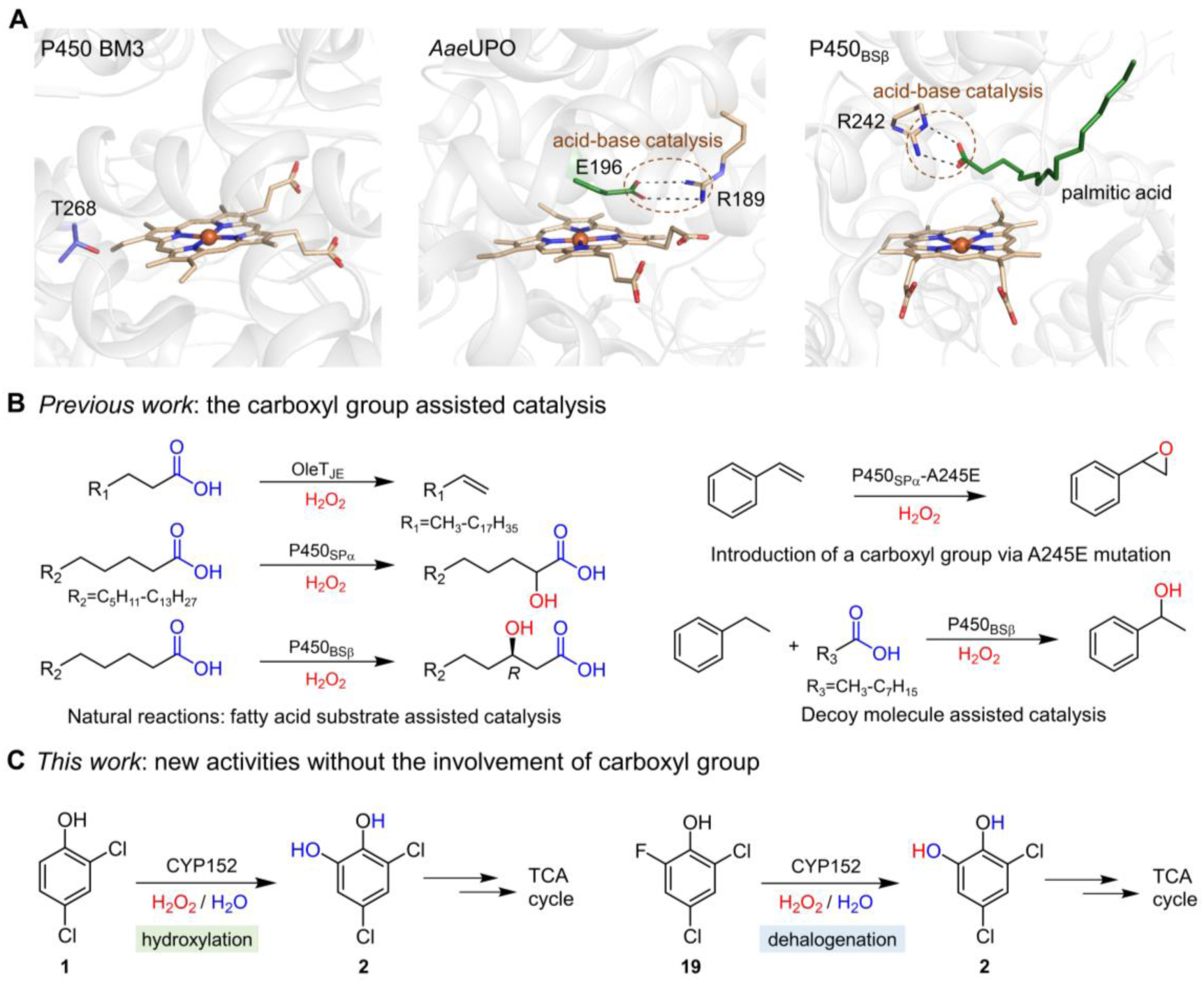
Substrate assisted acid-base catalysis of CYP152 peroxygenases. (A), Acid-base catalytic residue(s) in the active site of heme-dependent H_2_O_2_-utilizing enzymes: A representative P450 monooxygenase (*e.g.*, P450 BM3, PDB accession code: 3WSP) lacks native acid-base catalytic residues but possesses a conserved threonine which is important for proton transfer to form Cpd I. The introduction of an acid-base catalyst via T268E mutation significantly augments its peroxygenase activity. The fungal peroxygenase *Aae*UPO (PDB accession code: 2YOR) featuring a glutamic acid residue that interacts with an arginine residue for H_2_O_2_ activation. The bacterial P450 peroxygenase P450_BSβ_ with the substrate-derived carboxyl group interacting with a conserved arginine residue (PDB accession code: 1IZO), assisting in the formation of Cpd I upon H_2_O_2_ binding. (B), Previously reported hydroxylation, epoxidation and decarboxylation reactions catalyzed by various CYP152 peroxygenases with the assistance of a carboxyl group of different origins. (C), Novel activities of CYP152 peroxygenases towards non-carboxylic substrates. The catechol product serves as a key substrate in aromatic ring cleavage reactions, and commonly seen in biodegradation pathways of aromatic pollutants. TCA cycle: tricarboxylic acid cycle.

Recently, a class of benzene ring-containing small molecules was reported to enhance the catalytic activity of OleT_SA_ towards both hydrocinnamic acid and bromophenyl propionic acid, underscoring a flexible regulatory interaction between the P450 active center and small molecule modulator._[53]_ However, the precise manner how these modulators influence the catalytic activity of the P450 peroxygenase remained unclear. Moreover, a select number of P450 engineering studies revealed that the free space proximal to the heme-iron is crucial for the ability to utilize H_2_O_2_._[54,55]_

In this study, we delved into the catalytic promiscuity of CYP152 peroxygenases and, strikingly, identified 2,4-dichlorophenol (**1**) as the first non-carboxylic substrate that can be directly recognized by native P450 peroxygenases without any helper molecule (Figure 1C). Through engineering the substrate access channel, broadening the substrate scope, and identifying key catalytic groups, we unlocked the novel catalytic activities of CYP152 peroxygenases for aromatic hydroxylation and dehalogenation towards a wide range of non-carboxylic substrates including halophenols, nitrophenols, and cyanphenols. We also elucidated novel mechanisms for substrate recognition and oxidative modification of the non-carboxylic substrates using X-ray crystallography, ^18^O isotopic tracing experiments, and quantum mechanics/molecular mechanics (QM/MM) calculations.

## Results

### Discovery of the activity of CYP152 peroxygenases towards non-carboxylic substrates

Drawing inspiration from the significant enhancement of P450 peroxygenase activity by small molecules containing a benzene ring and by altering heme environment,_[53,54]_ we examined a select number of commercially available aromatic derivatives with varying substituents (−OH, −Cl, −SH, −NH_2_, −COOH, and −OCH_3_) as potential activity modulators by evaluating the catalytic performance of OleT_JE_, P450_BSβ_ and P450_SPα_ against myristic acid as substrate in the presence of each individual aromatic compounds (Figure S3). Despite no expected modulating effects, strikingly, both OleT_JE_ and P450_BSβ_ substantially hydroxylated the non-carboxylic substrate 2,4-dichlorophenol (**1**) at the *ortho*-position, producing 3,5-dichlorocatechol (**2**) regardless of the presence or absence of the fatty acid substrate (Figures 2A, 2B, S4-S5). Both conversion ratios were above 20% (Figure 2B) when using 5 μM AldO/10% glycerol as the *in situ* H_2_O_2_-releasing system._[48]_ These results essentially challenge the “common sense” that the catalytic activity of CYP152 peroxygenases requires a carboxyl group (from substrate, decoy molecule or mutated amino acid residue) for substrate recognition and hydrogen peroxide activation.

**Figure 2.**
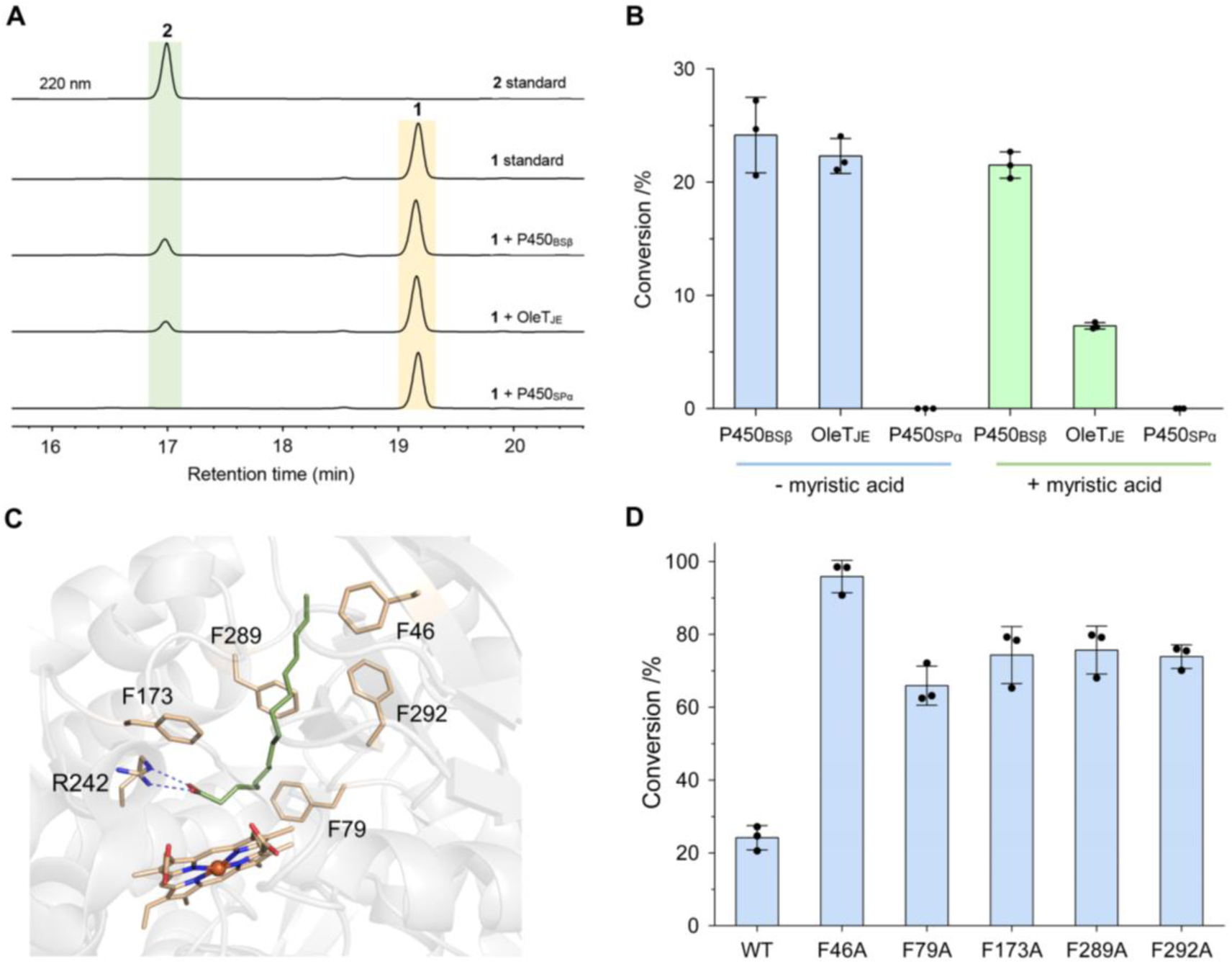
The activities of CYP152 peroxygenases towards non-carboxylic substrate 1. (A), HPLC analysis of the hydroxylation reactions of **1** catalyzed by three representative CYP152 peroxygenases. (B), The activities of three representative CYP152 peroxygenases towards **1** in the presence and absence of 2 mM myristic acid. (C), Five phenylalanine residues in the active site of P450_BSβ_ subject to alanine scanning. (D), The activities of the wild-type (WT) and mutant P450_BSβ_ towards **1**. Reaction conditions: 1 μM P450, 500 μM **1**, and the AldO/glycerol system (5 μM/10 %) to generate H_2_O_2_ *in situ* at 30 °C for 6 h.

### Engineering substrate access channel for activity improvement

We chose P450_BSβ_ for **1** hydroxylation activity improvement owing to its superior activity and stability._[44]_ The wild-type enzyme features a narrow and elongated substrate access channel to accommodate its natural fatty acid substrates._[29]_ Considering that an aromatic substrate likely favors more spacious substrate entrance channel and active site, we conducted alanine scanning for the five space-constraining phenylalanine residues including Phe46, Phe79, Phe173, Phe289 and Phe292 (Figure 2C). As expected, all the mutants (*i.e.*, P450_BSβ_-F46A, P450_BSβ_-F79A, P450_BSβ_-F173A, P450_BSβ_-F289A and P450_BSβ_-F292A) exhibited a significant increase in their activity towards **1**. In particular, P450_BSβ_-F46A efficiently converted **1** into **2** with the highest conversion ratio of 95.9%, alongside a marked enhancement in substrate binding affinity (*K_D_* = 7.6 μM) compared to the wild-type P450_BSβ_ (*K_D_* = 73.2 μM, Figures 2D, S6-S9). Thus, P450_BSβ_-F46A was selected for the following studies.

### Determination of non-carboxylic substrate specificity of P450_BSβ_-F46A

To investigate the non-carboxylic substrate specificity, we initially tested the activity of P450_BSβ_-F46A towards a number of halogen-free phenolic substrates including phenol (**3**), catechol (**4**), 2,4-dimethylphenol (**5**) and 2-methoxy-phenol (**6**), as well as non-phenolic halogenated benzene derivatives 1,3-dichlorobenzene (**7**), 2,4-dichloranisole (**8**), 2,4-dichlorobenzenethiol (**9**) and 2,4-dichlorobenzoic acid (**10**). The *in vitro* activity assays indicated that the non-carboxylic substrates devoid of either the phenolic hydroxyl group or halogen substituent failed to undergo enzymatic transformation, suggesting their indispensability for the catalytic activity of P450_BSβ_-F46A. (Figures 3, S10-S17). Next, a broader range of halogen phenol derivatives were examined (Figures 3, S18-S42, Tables S2-S4). For mono-halogenated phenols, P450_BSβ_-F46A preferentially catalyzed *para*- and *orth*o-hydroxylation of 2-chlorophenol (**11**) and 3-chlorophenol (**12**), respectively. By contrast, hydroxylation of 4-chlorophenol (**13**), 2-chloro-4-methylphenol (**14**), and 4-chloro-2-methylphenol (**15**) exclusively occurred at the *orth*o-position. The highest conversion ratio of 52.4% was achieved by **12**. Notably, the presence of an electron-donating methyl group negatively impacted the catalytic efficiency, resulting in significantly lower conversion ratios for **14** and **15**. As for di-halogenated phenols **1**, 2,4-dibromophenol (**16**), and 2,4-diiodophenol (**17**), P450_BSβ_-F46A demonstrated significantly higher activities and absolute *ortho*-hydroxylation selectivity compared to mono-halogenated phenols. Furthermore, P450_BSβ_-F46A showed the highest conversion ratio of 96.6% towards the tri-halogenated phenol 2,4,5-trichlorophenol (**18**). Taken together, it is evident that the presence of more halogen substituents led to higher substrate conversions (Figure 3).

**Figure 3.**
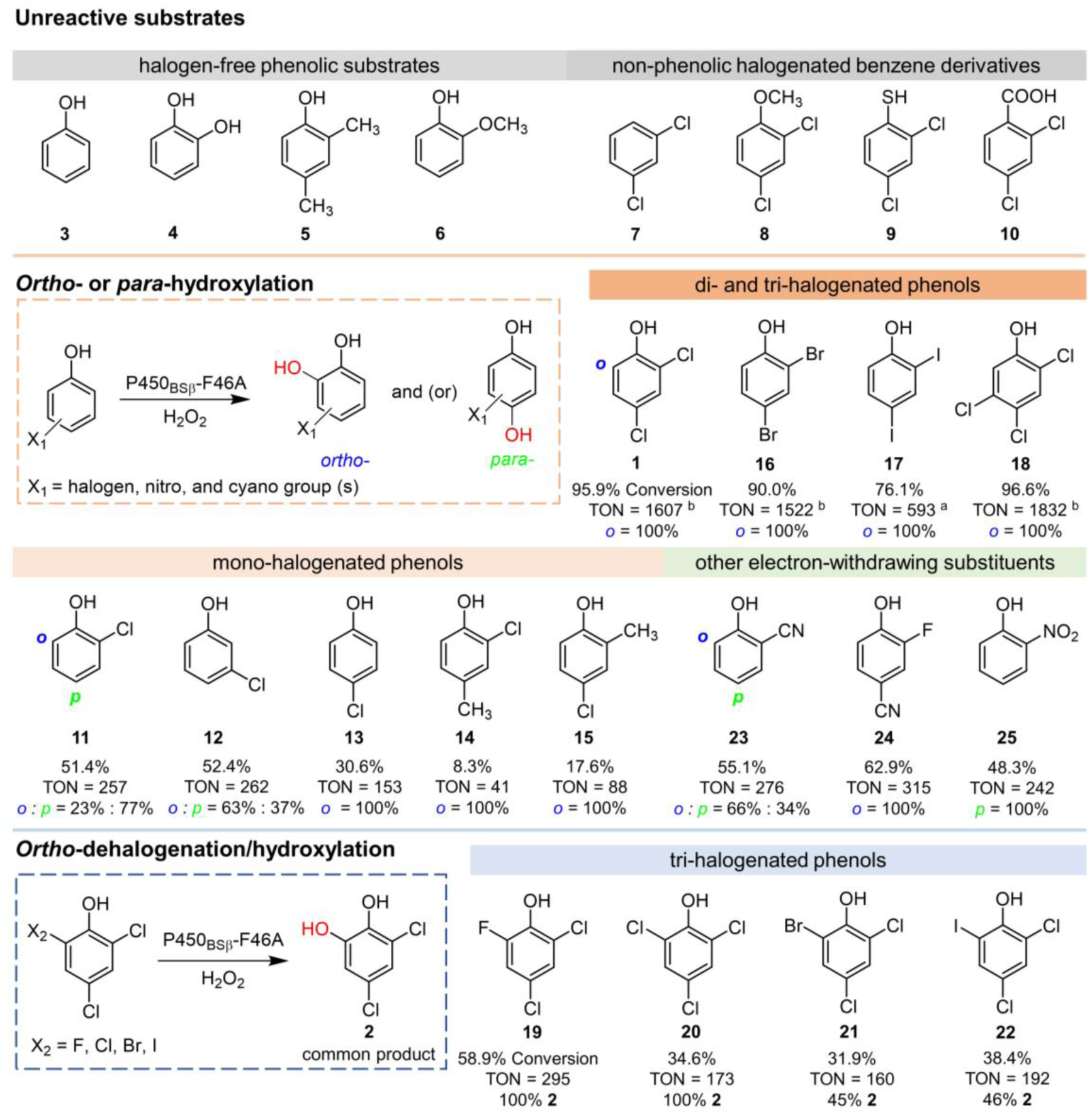
Catalytic activities of P450_BSβ_-F46A towards non-carboxylic substrates. A typical reaction containing 1 μM P450_BSβ_-F46A, 0.5 mM substrate, and the AldO-glycerol based H_2_O_2_ releasing system in 200 μL reaction buffer (pH 7.4, 50 mM NaH_2_PO_4_) was carried out at 30 °C for 6 h. The relative ratios between the *ortho*-(*o*) and *para*-hydroxylation (*p*) are indicated. *Notes*: ^a^1 mM substrate, ^b^2 mM substrate. See Supplementary materials for details.

Next, a broader range of halogen phenol derivatives were examined (Figures 3, S18-S42, Tables S2-S4). For mono-halogenated phenols, P450_BSβ_-F46A preferentially catalyzed *para*- and *orth*o-hydroxylation of 2-chlorophenol (**11**) and 3-chlorophenol (**12**), respectively. By contrast, hydroxylation of 4-chlorophenol (**13**), 2-chloro-4-methylphenol (**14**), and 4-chloro-2-methylphenol (**15**) exclusively occurred at the *orth*o-position. The highest conversion ratio of 52.4% was achieved by **12**. Notably, the presence of an electron-donating methyl group negatively impacted the catalytic efficiency, resulting in significantly lower conversion ratios for **14** and **15**. As for di-halogenated phenols **1**, 2,4-dibromophenol (**16**), and 2,4-diiodophenol (**17**), P450_BSβ_-F46A demonstrated significantly higher activities and absolute *ortho*-hydroxylation selectivity compared to mono-halogenated phenols. Furthermore, P450_BSβ_-F46A showed the highest conversion ratio of 96.6% towards the tri-halogenated phenol 2,4,5-trichlorophenol (**18**). Taken together, it is evident that the presence of more halogen substituents led to higher substrate conversions (Figure 3).

Intriguingly, the coupled dehalogenation/hydroxylation reactions were observed when three halogen atoms occupy the *para*- and both *ortho*-positions of the carboxyl-free phenol substrates, as exemplified by tri-halogenated phenols 2,4-dichloro-6-fluorophenol (**19**, Figure 1C), 2,4,6-trichlorophenol (**20**), 2,4-dichloro-6-bromophenol (**21**) and 2,4-dichloro-6-iodophenol (**22**). The apparent replacement of *ortho*-halogen with a hydroxyl group occurred with a declining efficiency in the order F > Cl > Br (Figures 3, S43-S54, Table S5). Among them, P450_BSβ_-F46A showed the highest activity against **19**, producing **2** at a 58.9% yield.

In consideration of the key role played by phenolic hydroxyl group and varying halogen substituents in substrate recognition, we further examined the catalytic activity of P450_BSβ_-F46A towards the carboxyl-free substrates that harbor alternative electron-withdrawing substituents, namely nitro and cyanide groups (Figures 3, S55-S64, Table S6). The P450 peroxygenase effectively recognized these compounds including 2-hydroxybenzonitrile (**23**), 3-fluoro-4-hydroxybenzonitrile (**24**) and 2-nitrophenol (**25**), catalyzing either *ortho*- or *para*-hydroxylation. These results consolidate the importance of the electron-withdrawing substituents.

### Mechanistic analysis of aromatic hydroxylation and dehalogenation/hydroxylation catalyzed by P450_BSβ_-F46A

To elucidate the mechanisms for P450_BSβ_-F46A mediated *ortho*-hydroxylation of **1** and *ortho*-dehalogenation/hydroxylation of **19** leading to the same product **2**, we first verified that **2** was the final stable product without other detectable intermediates by time-course experiments (Figures S65-S67).

Subsequently, we sought to determine the crystal structures of both substrate-free and substrate-bound forms of P450_BSβ_-F46A. However, despite our utmost efforts, effective removal of endogenous palmitic acid from the active site of the crystallized P450_BSβ_ mutant proved unattainable, similar to previous studies on CYP152 structures._[49,56]_ This could be due to the abundant presence of palmitic acid in *E. coli* and its high binding affinity with P450_BSβ_-F46A. Alternatively, the palmitic acid bound P450_BSβ_-F46A proteins were enriched by the crystallization conditions. Thus, we determined the complex structure of P450_BSβ_-F46A with palmitic acid (PDB accession code: 9IY1) at a resolution of 2.3 Å (Figure 4A, Table S7), which was utilized for further analysis. The structural alignment of P450_BSβ_-F46A and the wild-type P450_BSβ_ (P450_BSβ_-WT) structures (PDB accession code: 1IZO)_[29]_ reveals highly similar conformation, with a root mean square deviation (RMSD) of 0.282 Å over 372 C*α* atoms (Figure 4B). However, compared to P450_BSβ_-WT, the F46A substitution effectively expands the distal substrate access channel (Figures 4C, 4D, S68), thereby facilitating the accessibility of aromatic substrates.

**Figure 4.**
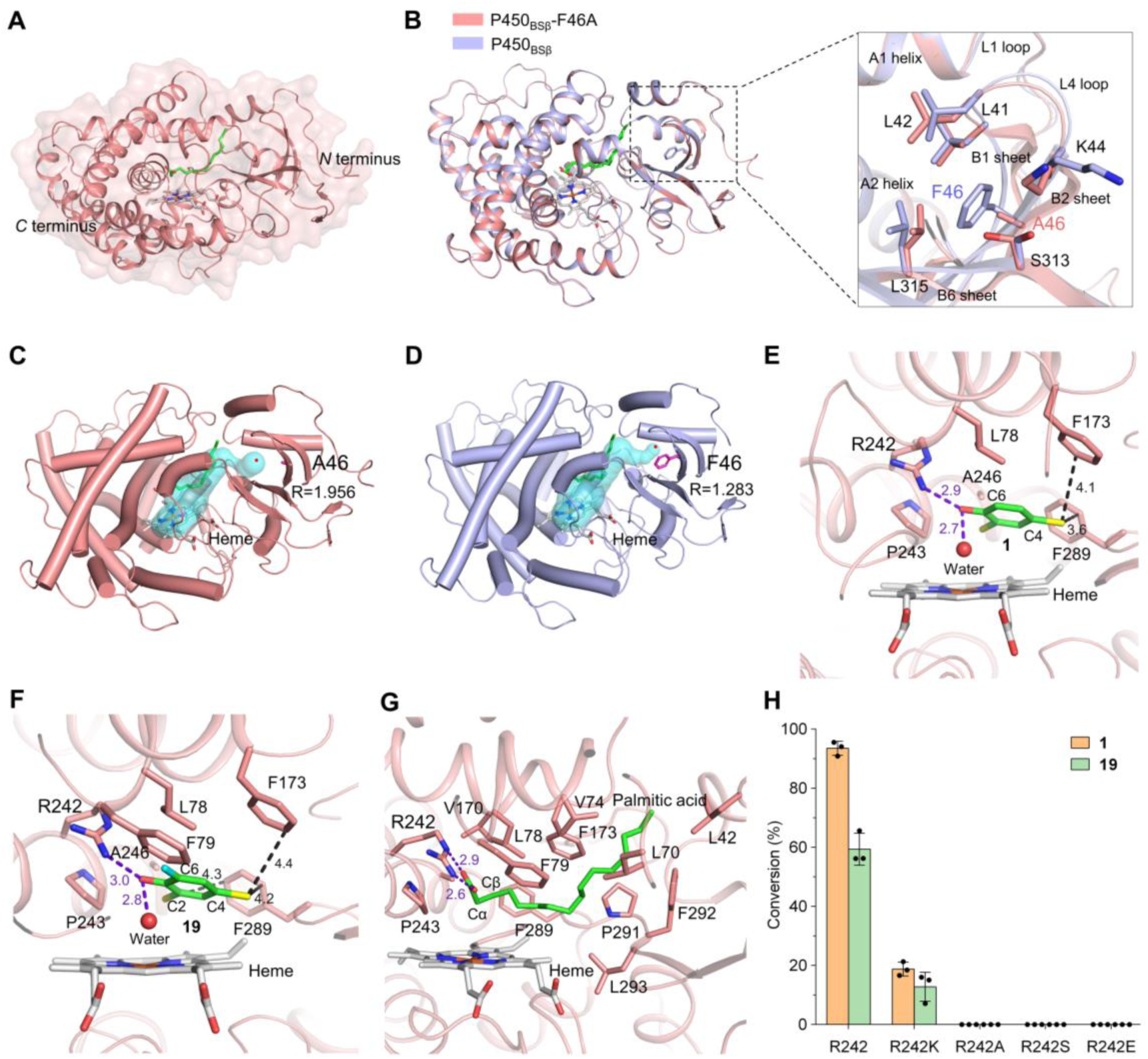
Structural analysis of P450_BSβ_-F46A. (A), Overall structure of P450_BSβ_-F46A (PDB accession code: 9IY1). The heme group (grey) and palmitic acid (green) are shown as sticks. (B), Structural overlay of P450_BSβ_-F46A and wild-type P450_BSβ_ with the modified region amplified. (C and D), Comparative analysis of the crystal structures of P450_BSβ_-F46A (C) and P450_BSβ_ (D) at the same view to highlight their different substrate access channels. The channels are determined using the Caver 3.0.3 plugin in PyMOL 2.3.2 and shown in cyan. The central point of the distal channel is denoted by red dots, with ‘R’ representing its radius in angstrom (Å). (E and F), Structures of P450_BSβ_-F46A with docked substrate **1** (E) and **19** (F) in the active site. Key interacting amino acids are labelled and shown as sticks, including R242 involved in polar interactions, F173 and F289 engaged in halogen-π interactions, and hydrophobic amino acids for hydrophobic interactions (within a 4.5 Å distance range). The carbon atoms of both substrates are depicted in green, chlorine atoms in yellow, fluorine atom in light blue, and hydroxyl groups in red. Purple dashed lines represent hydrogen bonds, while black dashed lines indicate halogen-π bonds in angstrom. (G), The active site of P450_BSβ_-F46A in complex with palmitic acid. Residues mediating the interactions between palmitic acid and P450_BSβ_-F46A are labelled and shown as sticks and polar interactions are indicated by purple dashed lines in angstrom. (H), Catalytic activities of different R242 mutants of P450_BSβ_-F46A. Reaction conditions: 1 μM P450, 500 μM **1** or **19**, and the AldO/glycerol system to generate H_2_O_2_ at 30 °C for 6 h.

To better understand the interaction mechanism of P450_BSβ_-F46A with non-carboxylic substrates, we removed palmitic acid from the active site *in silico*. Molecular docking was further conducted using AutoDockTools 1.5.6_[57]_ to predict the binding modes of **1** and **19** in P450_BSβ_-F46A. The docking results revealed that **1** and **19** exhibited similar binding modes, characterized by the formation of stable hydrogen bonds between the phenolic hydroxyl group and residue R242, with bond lengths of 2.9 Å and 3.0 Å, respectively (Figures 4E, 4F). Additionally, the halogen atoms of the substrates establish halogen-π interactions with the surrounding phenylalanine residues. Specifically, the C4-Cl of substrate **1** interacts with F173 and F289, while the C4-Cl of substrate **19** interacts with F173 and both C2-Cl and C4-Cl show halogen-π interactions with F289. Consequently, the C6-H of **1** and C6-F of **19** are positioned above the heme iron (at 5.6 and 5.4 Å, respectively) along with a water molecule (Figures 4E, 4F). Moreover, hydrophobic interactions are also observed between substrates and surrounding hydrophobic amino acid residues (Figures 4E, 4F). These binding modes are distinct from that of palmitic acid, whose carboxyl group interacts with R242 with the distances between the two oxygen atoms of carboxyl group and the guanidyl group being 2.6 and 2.9 Å, respectively (Figure 4G). Beyond electrostatic interaction, the fatty acid substrate is further stabilized by hydrophobic interactions with the alkyl side chain of surrounding hydrophobic residues including L42, L70, V74, L78, F79, F173, F289, F292 and L293 (Figure 4G). This specific binding mode positions the linear substrate almost perpendicular to the heme plane, with the C_α_ and C_β_ carbons situated closest to the heme iron (at 4.6 and 5.3 Å, respectively; Figure 4G). The interaction between the guanidyl group of R242 and halogenated phenolic hydroxyl group derived from a non-carboxylic substrate demonstrates a novel substrate recognition and binding mechanism, presenting an unprecedented perspective on the acid-base catalysis of heme-dependent H_2_O_2_-utilizing enzymes._[19,58]_ Site-directed mutagenesis highlighted the pivotal role of the conserved arginine residue in facilitating the catalytic process of non-carboxylic substrates. Specifically, substituting this basic R242 in P450_BSβ_-F46A with a nonpolar, neutral or acidic amino acid (yielding R242A, R242S, and R242E variants), led to complete activity abolishment (Figures 4H, S69-S72). By contrast, R242K mutant with an analogous replacement by another basic residue lysine retained approximately 20% activity, suggesting that R242 should still act as an acid-base catalyst during the enzymatic transformation of **1** and **19**.

To unravel the catalytic mechanisms of P450_BSβ_-F46A-mediated **2** formation from *ortho*-hydroxylation of **1**, combined molecular dynamics (MD) simulations and quantum mechanics/molecular mechanics (QM/MM) calculations were conducted. MD simulations indicate that both hydrogen bond networks (Figures 4E, 4F) are fairly stable, facilitating the formation of a hydrogen bond between the substrate −OH group and the distal O2 of H_2_O_2_ (Figure S73). The catalytic mechanism was investigated based on the MD equilibrated conformation. First, the direct homolytic cleavage of Fe(III)---H_2_O_2_ (Figure S74) and proximal proton transfer from H_2_O_2_ to the substrate’s chlorine ion (Figure S75) was excluded. Then, we investigated the possible route that the substrate’s hydroxyl group may drive the O−O cleavage of H_2_O_2_ via a proton-coupled electron transfer (PCET) process. QM/MM calculations support that such reaction pathway is accessible, involving a barrier of 21.8 kcal/mol (RC_[1]_ → TS1_[1]_, Figures 5A, S76), generating Fe(IV)-OH^−^ and H_2_O with an exothermic release of 13.8 kcal/mol (IC1_[1]_). The generated H_2_O subsequently facilitates the hydrogen atom transfer (HAT) from Fe(IV)-OH^−^ to the substrate radical, forming Fe(IV)=O^2–^ (Cpd I). This step involves a small barrier of 3.4 kcal/mol and exothermicity of 27.6 kcal/mol (IC1_[1]_ → IC2_[1]_, Figures 5A, S76). Thus, our computations identify a novel mechanism for P450_BSβ_-F46A catalyzed oxidation of **1** that involves the substrate OH group assisted generation of Cpd I via a PCET process.

**Figure 5.**
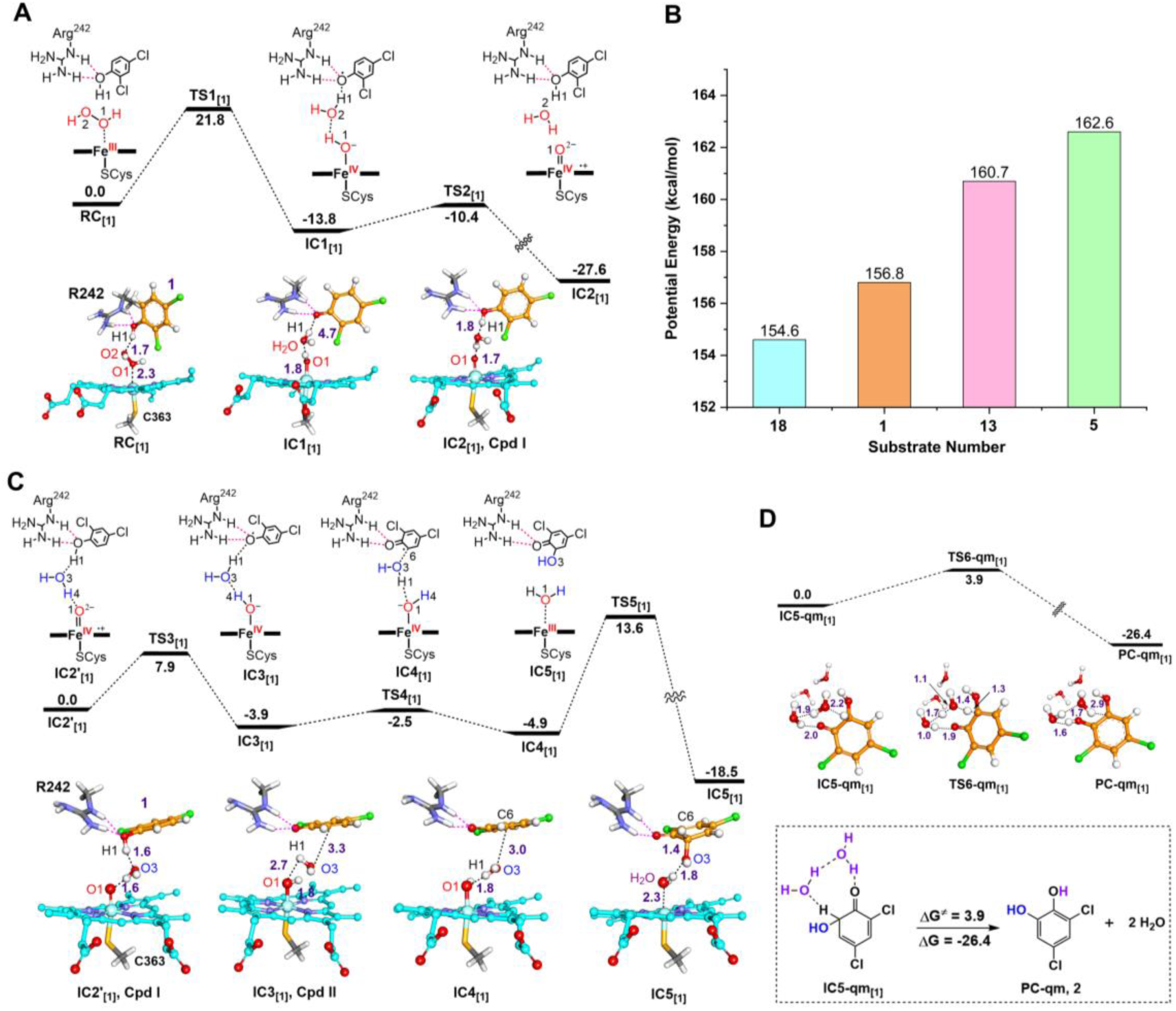
Mechanism of the aromatic hydroxylation of 1 catalyzed by P450_BSβ_-F46A. (A), The calculated mechanism (with energies in kcal/mol) for the formation of Cpd I by P450_BSβ_-F46A in the presence of the non-carboxylic substrate **1**. The phenolic hydroxyl group interacting with the guanidyl group of R242 is located above the heme. The key distances are given in angstrom (Å). (B), QM calculated relative energies (in kcal/mol) for the ability of **1**, **5**, **13**, and **18** to provide protons in the presence of R242. (C), QM/MM calculated mechanism for the formation of ene-ketone intermediate starting from Cpd I of P450_BSβ_-F46A, along with the QM/MM optimized structures of key species involved in the reactions. (D), QM calculated energy barrier (in kcal/mol) for the water-mediated proton transfer for the generation of **2**.

Next, we comprehensively evaluated the effects of R242 and substrate during the formation of Cpd I. First, QM model calculations indicate that the energy barrier for H_2_O_2_ cleavage without R242 (ΔE^≠^ = 21.1 kcal/mol) is ∼4 kcal/mol higher than that with R242 (ΔE^≠^ = 17.3 kcal/mol), confirming that this arginine residue enhances the acid-base catalysis of the substrate OH group. (Figure S77). In addition, the mutant P450_BSβ_-F46A-R242A displayed a reduced affinity for **1**, as evidenced by the higher *K*_D_ value than that of P450_BSβ_-F46A, indicating that the arginine also contributes to substrate binding (Figure S78, Table S9). Of note, we noticed that in the presence of non-carboxylic substrates, the hydroxyl group of the substrate is required for the activation of H_2_O_2_ through PCET, leading to the formation of Cpd I (Figure 5A). Thus, there arises a natural question: does the presence of a hydroxyl group in substrate suffice? Despite the substrate OH group is necessary for Cpd I generation, we found that **5** with the OH group failed to undergo a reaction with P450_BSβ_-F46A (Figure 3, Table S9). To understand this, we compared the acidity of **1**, **5**, **13**, and **18** in the presence of R242. QM calculations show that proton dissociation energies of **18** and **1** are much lower than those of **13** and **5** (Figure 5B). This is likely because electron-withdrawing substituents (*e.g.*, halogens, nitro groups) could enhance the acidity of the phenolic hydroxyl group and stabilize the delocalized electrons of phenoxy group._[59]_ Thus, both R242 and electron-withdrawing substituents are key to the acidity of substrate OH group, which in turn enhances the PCET process for Cpd I generation (Figure S79).

After the formation of Cpd I, a stable water bridge exists between the O1 of Cpd I and the hydroxyl group of the substrate. The bridge water is initially generated from the homolytic cleavage of H_2_O_2_. However, MD simulations show this water can undergo exchange with the solvent water, which then participates in subsequent reactions (Figure S80). As shown in Figure 5C, starting from IC2’_[1]_, Cpd I readily accepts H1 atom of the substrate, overcoming an energy barrier of 7.9 kcal/mol (IC2’_[1]_ → TS3_[1]_ in Figures 5C, S81) to form Cpd II and a semiquinone radical intermediate (IC3_[1]_). Subsequent water conformational change undergoes a very low energy barrier (1.4 kcal/mol), orienting H1 of the water towards O1 of Cpd II, which is more suitable for water cleavage. After overcoming an energy barrier of 18.5 kcal/mol, the reaction forms the ene-ketone intermediate IC5_[1]_ with exothermicity of 18.5 kcal/mol. In contrast, the direct attack the substrate by Cpd II requires overcoming a significantly higher energy barrier of 45.0 kcal/mol (Figure S82). This is due to the long distance between O1 and the substrate’s C6 (5.3 Å) and the presence of a water molecule between them, thus causing considerable steric hindrance. Finally, in aqueous solution, IC5_[1]_ undergoes an intramolecular rearrangement to form the product **2** (Figure 5D). To confirm the proposed reaction pathway in which the OH radical attacks the substrate after water cleavage, we performed isotope tracing experiments using H_2_^18^O_2_/H_2_^18^O. It was clear that the hydroxyl oxygen atom of **2** does not originate from hydrogen peroxide; instead, LC-MS analysis indicated that it should derive from water (Figures S83-S85).

For substrate **19**, the catalytic mechanism is highly similar to that of **1**. The hydroxylation specifically occurs at the F-substituted C6 instead of the Cl-substituted C2 (Figures 6, S86). This preference arises from the need to overcome a markedly higher energy barrier when attacking the Cl-substituted C2 (30.4 kcal/mol). Despite the conformational alteration, the most critical factor lies in the F-substitution at C6, which significantly reduces the electronegativity of other C atoms and inhibits the coupling of OH radical to C2 (Figure 6A). Then solvent water mediates the proton transfer from the intermediate OH to F^-^, forming HF and a quinone intermediate (Figures 6B, S87-S88). This intermediate then abstracts two hydrogen atoms from H_2_O_2_ and H_2_O, generating the product **2** and O_2_ (Figures 6C, S89). In addition, our isotopically labeled H_2_^18^O_2_/H_2_^18^O tracing experiments supported that the oxygen atom in **2** indeed originated from water, consistent with our calculation results (Figures S83, S90-S91). Notably, the observed selectivity towards this specific dehalogenation product does not extend to other substrates such as **21**. This phenomenon is primarily attributed to the energy barriers for attacking the Cl-substituted C2 and Br-substituted C6 are at a comparatively level, leading to the generation of two products in similar proportions (Figures 3, S92).

**Figure 6.**
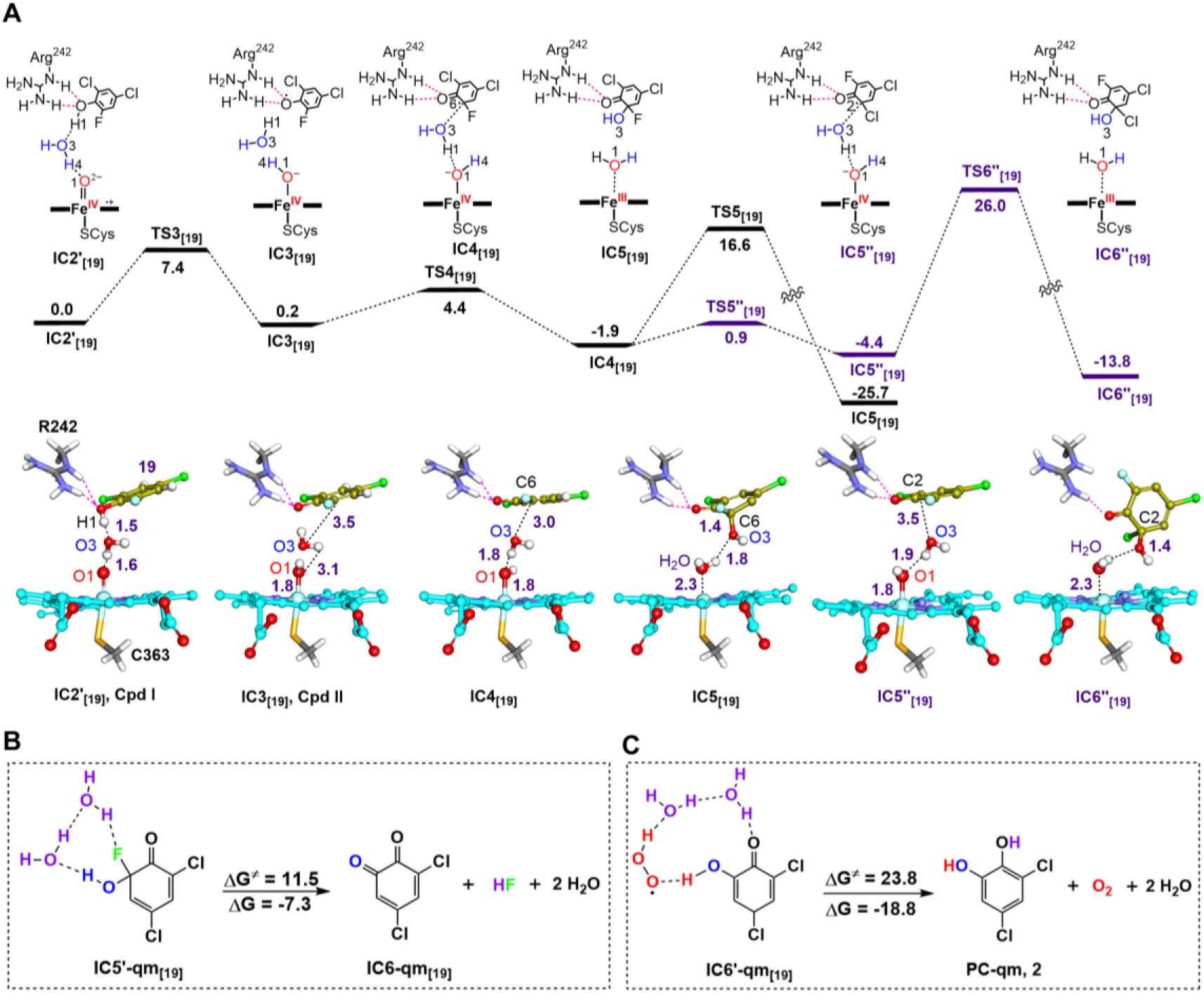
Mechanism of the dehalogenation/hydroxylation of 19 catalyzed by P450_BSβ_-F46A. (A), QM/MM calculated mechanism for the formation of hydroxylation intermediate starting from Cpd I of P450_BSβ_-F46A. The phenolic hydroxyl group interacting with the guanidyl group of R242 is located above the heme, along with the QM/MM optimized structures of key species involved in the reactions. The key distances are given in angstrom (Å). (B), QM calculated energy barrier (in kcal/mol) for the water-mediated proton transfer to generate the quinone intermediate and HF. (C), QM calculated energy barrier (in kcal/mol) for the water-mediated hydrogen transfer from H_2_O_2_ to quinone for the generation of **2** and O_2_.

Overall, the catalytic mechanism involves the substitution of halogen groups on the benzene ring, thus facilitating the formation of an acid-base pair between the substrate and R242. The interaction promotes the acid-base catalytic activation of H_2_O_2_, leading to the formation of Cpd I. The process includes the following steps: (1) Initial proton transferred from the substrate hydroxyl group drives electron transfer, promoting the homolytic cleavage of H_2_O_2_, which is the rate-determining step. The acidity of the substrate substituent correlates positively with the promotion of the PCET process. (2) Cpd I mediates the HAT process from the substrate hydroxyl group to form Cpd II. (3) Cpd II then facilitates the cleavage of a water molecule, generating the resting state Fe(III)–H_2_O and the *ene*-ketone intermediate, which is evidenced by the ^18^O tracing experiments showing that the hydroxyl oxygen atom in the product should originate from solvent water. (4) Finally, the *ortho*-hydroxylation products are formed in the presence of water molecules.

### Catalytic activity and substrate binding affinity of CYP152 peroxygenases

To further evaluate the catalytic activity and application potential of CYP152 peroxygenases to deal with non-carboxylic substrates, we comparatively determined the turnover numbers (TONs) of P450_BSβ_-F46A when supported by the *in situ* H_2_O_2_ releasing system consisting of AldO and glycerol (Table S8). Again, the results (Figure 3) further confirmed a positive correlation between the halogen content of the substrate and the catalytic efficiency, exemplified by the highest TON observed for the tri-halogenated substrate **18** (TON = 1832), significantly outperforming that of the di-halogenated **1** (TON = 1607) and mono-halogenated **13** (TON = 153), with a remarkable 12-fold enhancement over **13**. Although both **18** and **20** are trichlorophenols, **20** displayed a markedly lower dehalogenation/hydroxylation activity than the aromatic hydroxylation activity of **18**. Beyond the influence of substrate acidity as mentioned above, to rationalize these discrepancies on multiple dimensions, we assessed the substrate binding affinities (*K_D_*) of these non-carboxylic substrates (Table S9, Figures S93-S96). As a result, the poorer substrate **20** was found to exhibit a higher binding affinity than **18** (*K_D_* = 4.5 μM of **20** *versus* 12.9 μM of **18**). Interestingly, substrates **1** and **14** only differ in one substituent on the benzene ring, but the aromatic hydroxylation activity of P450_BSβ_-F46A to **14** (TON = 41) with a *para*-methyl group was much lower than that to **1** (TON = 1607) with a corresponding chlorine substituent (Figure 3, Table S8). This activity difference could not be explained by the insignificant variation in their dissociation constant values (*K_D_* = 7.6 μM of **1** *versus* 4.0 μM of **14**). Furthermore, substrates **14** and **15**, which differ from each other solely in the positioning of their substituents on the benzene ring, showed a remarkable approximately twofold difference in catalytic activity, despite their comparable substrate binding affinities (*K_D_* = 4.0 μM of **14** *versus* 4.6 μM of **15**). These results underscore the paramount importance of substituent positioning and electronic properties in dictating the catalytic efficiency of P450_BSβ_-F46A.

## Discussion

In this study, we discovered and engineered the ability of P450 peroxygenases to catalyze the oxidative transformations (aromatic *ortho*- and *para*-hydroxylation and oxidative dehalogenation/hydroxylation) of an array of non-carboxylic substrates for the first time, thus expanding the functions of the CYP152 family peroxygenases. These findings not only present new possibilities for development of novel aromatic hydroxylases and dehalogenases, but also challenge the conventional concept that heme-dependent H2O2-utilizing enzymes necessitate a carboxyl group of different origins in the active site for acid-base catalytic activation of H_2_O_2_ (Figures 1A, 1B). The structure-function relationship (Figure 3) indicates the new activities derive from the phenolic hydroxyl group activated by electron-withdrawing substituent(s), which functionally replaces the carboxyl group to maintain the acid-base catalysis.

Mechanistically, this unusual catalysis is initiated by the involvement of a water molecule, which plays a crucial role in abstracting a hydrogen atom from the phenolic O−H bond, forming a phenoxy radical and Cpd II. Then, the phenoxy radical undergoes a rearrangement to form the semiquinone radical. Following the mediation of water cleavage by Cpd II, the oxygen atom of the newly formed hydroxyl group originating from water, bypasses the conventional oxygen rebound mechanism. To the best of our knowledge, this is the first report detailing the water-dependent hydroxylation and dehalogenation reactions of halophenols catalyzed by a P450 enzyme so far, representing a mechanism fundamentally distinct from the canonical hydroxylation of aliphatic carboxylic acid by P450_BSβ_ via typical C_β_-H abstraction._[49]_ Our finding underscores the emergence of CYP152 peroxygenases as a key enzyme resource for delving into catalytic versatility and promiscuity. The continuous discovery of new activities in this P450 family highlights the immense potential for innovative reactions and applications.

Of note, P450_BSβ_-F46A achieved a considerable TON of 1832 towards **18** with the *in situ* H_2_O_2_ releasing system, showcasing its potential as a novel and useful biocatalyst for degradation of environmental pollutants. The catechol products of halophenols derived from *ortho*-hydroxylation or *ortho*-dehalogenation/hydroxylation (Figure 3) are vital substrates in aromatic ring cleavage reactions in biodegradation pathways of many aromatic pollutants_[60]_ (Figure 1C). In contrast to previously reported P450 enzymes, which often suffer from limited efficiency and instability,_[61]_ the engineered P450 peroxygenases showed considerable substrate scope and superior catalytic activities, rendering it a more practical and eco-friendly tool for enzyme/substrate engineering, biocatalysis, and the removal of recalcitrant and xenobiotic pollutants. This significant expansion in biodegradation capability paves a promising way for future application of P450 peroxygenases in the field of bioremediation.

Despite the fact that the efficiencies of the new reaction types fall short of that of normal fatty acid hydroxylation and decarboxylation reactions,_[47,48]_ it nonetheless serves as a motivating impetus for the future endeavors to enhance activity and reusability through classical enzyme engineering and enzyme immobilization, as well as to expand the competence of P450 peroxygenases by incorporating non-canonical amino acids,_[62,63]_ thereby pushing the boundaries of enzymatic catalysis.

## Acknowledgments

This work was financially supported by the National Natural Science Foundation of China (32025001 to S.L., 32300021 to Y.J., 32200017 to Z.L., 22403094 to W.P.), China Postdoctoral Science Foundation (2022M710080 to Y.J.), Postdoctoral Innovation Project of Shandong Province (SDCX-ZG-202201005 to Y.J.), Shandong Provincial Natural Science Foundation (ZR2023QC021 to Y.J., ZR2022QC070 to Z.L.), and Qingdao Natural Science Foundation (23-2-1-25-zyyd-jch to Y.J.). We thank Zhifeng Li, Jingyao Qu, Guannan Lin and Haiyan Sui from the State Key Laboratory of Microbial Technology at Shandong University for their assistance in GC-MS, LC-MS and NMR data collection. We also thank the staff of the BL19U1 beamline at the National Facility for Protein Science Shanghai (NFPS) for their help with data collection, and Xiaoju Li from the Core Facilities for Life and Environmental Sciences at Shandong University for her assistance with XRD.

## Table of Contents

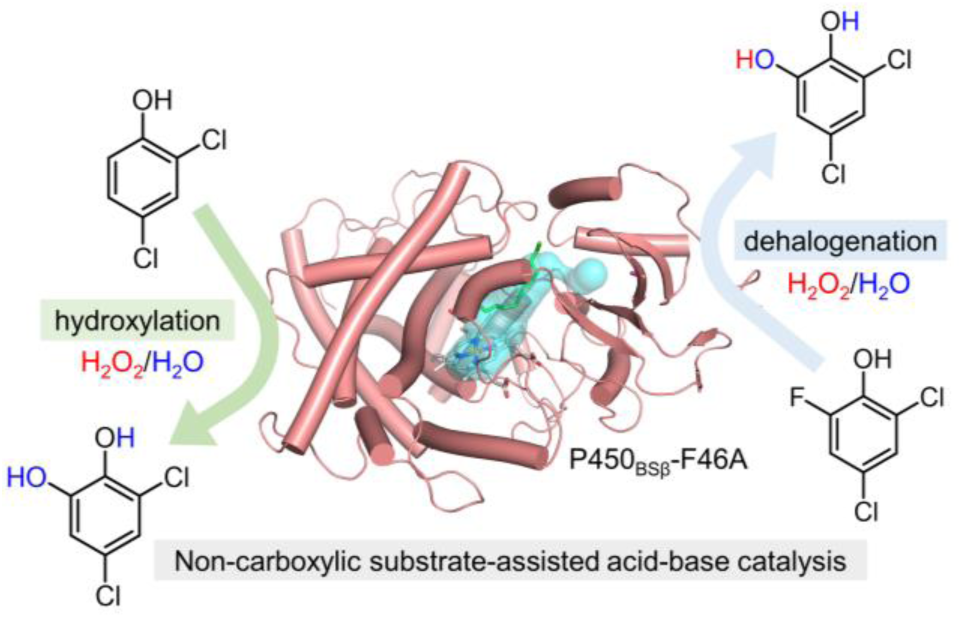

## Notes

### Competing Interest Statement

The authors have declared no competing interest.

