## Supplemental information for "Unexpected Activities of CYP152 Peroxygenases towards Non-carboxylic Substrates Reveal Novel Substrate Recognition Mechanism and Catalytic Versatility"

##### Table of Contents

Experimental Procedures.........................................................................................2-8

Results...........................................................................................................................9

Supporting tables.........................................................................................9-17

Supporting figures.......................................................................................18-67

References..................................................................................................68-69

#### Experimental Procedures

##### 1.1 General materials

The strains of *Escherichia coli* DH5*α* and BL21(DE3) and plasmid pET28b were preserved by our laboratory. All antibiotics and chemicals including 2,4-dichlorophenol (**1**), 3,5-dichlorocatechol (**2**), phenol (**3**), catechol (**4**), 2,4-dimethylphenol (**5**), 2-methoxy-phenol (**6**), 1,3-dichlorobenzene (**7**), 2,4-dichloranisole (**8**), 2,4-dichlorobenzenethiol (**9**), 2,4-dichlorobenzoic acid (**10**), 2-chlorophenol (**11**), 3-chlorophenol (**12**), 4-chlorophenol (**13**), 2-chloro-4-methylphenol (**14**), 4-chloro-2-methylphenol (**15**), 2,4-dibromophenol (**16**), 2,4-diiodophenol (**17**), 2,4,5-trichlorophenol (**18**), 2,4-dichloro-6-fluorophenol (**19**), 2,4,6-trichlorophenol (**20**), 2,4-dichloro-6-bromophenol (**21**), 2,4-dichloro-6-iodophenol (**22**), 2-hydroxybenzonitrile (**23**), 3-fluro-4-hydroxybenzonitrile (**24**), 2-nitrophenol (**25**), 1,3,5-trichlorobenzene, 2-chlorobenzenethiol, benzoic acid, 4-hydroxybenzoic acid, pyrocatechol, 2-hydroxybenzoic acid, 2-chlorobenzoic acid, aniline, 2,4-dichloranisole, 2-chlorobenzenethiol, 2-chlorohydroquinone, 2-nitro-4-hydroxyphenol, 3-chlorocatechol, 4-chlorocatechol, 3-chloro-5-methylcatechol, 3,4,6-trichlorobenzene-1,2-diol, 3-bromo-5-chlorobenzene-1,2-diol, 2,5-dihydroxybenzonitrile, 2,3-dihydroxybenzonitrile, 3,4-dihydroxybenzonitrile, $H_2{}^{18}O$, $H_2{}^{18}O_2$ were purchased from Solarbio (Beijing, China), Sigma Aldrich (St. Louis, MO, USA), TCI (Shanghai, China), Macklin (Shanghai, China), Aladdin Biochemical Technology (Shanghai, China), Bidepharm (Shanghai, China), or Shanghai yuanye Bio-Technology Co Ltd (Shanghai, China). Purification of DNA fragments was performed using the MonPure™ Gel & PCR Clean Kit from Baisai Biotechnology (Qingdao, China). Ni-NTA resin used for protein purification was obtained from Sangon Biotech (Shanghai, China). PD-10 desalting columns were purchased from GE Healthcare (Piscataway, NJ, USA). Millipore Amicon Ultra centrifugal filters were bought from Millipore (Billerica, MA, USA). The 10 × QuickRun™ Fast Running Buffer and FlexiRun™ premixed gel solution for SDS-PAGE analysis, GelRed, Luria-Bertani broth (LB) and Terrific broth (TB) were supplied by MDBio (Xinbei, China). I-5™ 2 × High-Fidelity Master Mix (TP001) and 2 × T5 Super PCR Mix (Colony, TSE005) were purchased from Beijing Tsingke Biotech Co., Ltd. (Beijing, China).

##### 1.2 Molecular cloning and protein purification

The constructs of single mutants including $P450_{BS\beta}$-F46A, $P450_{BS\beta}$-F79A, $P450_{BS\beta}$-F173A, $P450_{BS\beta}$-F289A and $P450_{BS\beta}$-F292A, and double mutants including $P450_{BS\beta}$-F46A-R242A,

$P450_{BS\beta}$-F46A-R242E, $P450_{BS\beta}$-F46A-R242S and $P450_{BS\beta}$-F46A-R242K were prepared by site-directed mutagenesis via overlap extension PCR using $P450_{BS\beta}$ and $P450_{BS\beta}$-F46A encoding genes as templates, respectively. The sequences of primers used in this study are listed in Table S1. All cloned sequences were confirmed by DNA sequencing at Sangon Biotech (Shanghai, China), and then used to transform *E. coli* BL21(DE3) for protein expression. The plasmids pET28b-*bsβ*, pET28b-*oleT,* pET28b-*spα,* pET28b-*aldo* were constructed by this laboratory previously.[1]

The *E. coli* BL21(DE3) cells carrying a certain recombinant expression vector were grown at 37 °C for 12 h with shaking at 220 rpm and then used as a seed culture to inoculate (1:50 ratio) a modified Terrific Broth medium containing rare salt solution. Cells were grown at 37 °C for 3-4 h until the optical density at 600 nm ($OD_{600}$) reached 0.8 to 1.0, to which 0.2 mM isopropyl *β*-D-1-thiogalactopyranoside (IPTG) was added. For P450 expression, 1 mM *δ*-aminolevulinic acid (5-ALA) and 1 mM thiamine were also supplemented. Afterwards, the cultivation continued for another 24 h at 18 °C. The cells were harvested (6,000 *g*, 4 °C, 10 min) and stored at -80°C for later use. Purification of the $His_6$-tagged AldO, $P450_{SP\alpha}$, $OleT_{JE}$, $P450_{BS\beta}$ and its mutants were performed by following the previously established procedure.[1b]

**1.3 Enzyme concentration determination**

Analysis of the UV-visible spectroscopic properties of $P450_{BS\beta}$ and its mutants were carried out as described previously.[2] The P450 protein concentrations were calculated based on their CO-bound reduced difference spectra using the reduced differential extinction coefficient $\varepsilon_{450\text{-}490}$ of 91,000 $M^{-1}$ $cm^{-1}$.[3]The concentration of AldO was determined at 452 nm with the reported extinction coefficient of 12,500 $M^{-1}$ $cm^{-1}$.[4]

**1.4 *In vitro* enzymatic assays**

A typical assay for CYP152 peroxygenases and related mutants containing 1 μM P450, 0.5 mM substrate, 5 μM P450 AldO, 10% glycerol and 5% EtOH as co-solvent in 200 μL reaction buffer (pH 7.4, 50 mM $NaH_2PO_4$) was carried out at 30 °C for 6 h.

**Determination of turnover number (TON):** The AldO/glycerol-based *in situ* releasing $H_2O_2$ reaction mixture contained 1 μM P450, 0.5-5 mM substrates, 5 μM AldO, 10% glycerol, and 5% EtOH as co-solvent in 200 μL reaction buffer. The reactions were performed at 30 °C for 6 h.

**Determination of substrate dissociation constants ($K_D$):** 1 mL reaction mixture containing 1 μM of P450 ($P450_{BS\beta}$, $P450_{BS\beta}$-F46A, or $P450_{BS\beta}$-F46A-R242A) in the reaction buffer was titrated with sequential additions of 1-20 mM substrate stock (lauric acid, **1**, **5**, **7**, and **10-20**) dissolved in 10% ethanol (1-20 μL). The amount of ethanol did not exceed 1% *vol*/*vol*. The additive absorbance changes at 390 nm and 420 nm were fit as a function of substrate concentrations using OriginPro 8.5 program to calculate the dissociation constant ($K_D$) values.

**Time-course experiments:** The reaction mixture contained 1 μM P450, 0.5 mM substrate (**1** or **19**), 10% glycerol, and 5% EtOH as co-solvent in a 1.2 mL reaction system. The reaction was initiated by adding 5 μM AldO. Aliquots (150 μL) of reactions were taken and quenched at different time points (0, 0.5, 1, 2, 4, and 6 h) by thoroughly mixing with 150 μL acetonitrile.

**The $^{18}$O-tracing experiments using $H_2{}^{18}O_2$:** A 200 μL reaction containing 1 μM P450$_{BS\beta}$-F46A, 500 μM **1** or **19** as substrate, 500 μM $H_2{}^{18}O_2$ or $H_2O_2$ as cofactor, and 5% EtOH as co-solvent was carried out at 30 °C for 6 h.

**The $^{18}$O-tracing experiments using $H_2{}^{18}O$:** The 200 μL reaction containing 1 μM P450$_{BS\beta}$-F46A, 500 μM **1** or **19** as substrate, 500 μM $H_2O_2$, and 5% EtOH as co-solvent in 200 μL reaction buffer made of $H_2O$ or $H_2{}^{18}O$. The reactions were performed at 30°C for 6 h.

**1.5 Up-scaling reaction of the P450$_{BS\beta}$-F46A/AldO system using non-carboxylic substrates (15, 16, 17, 22, 24) for the preparation of related products.**

Conversion of 10 mg (~500 μM) each non-carboxylic substrates (**15**, **16**, **17**, **22**, or **24**) was achieved using 1 μM P450$_{BS\beta}$-F46A and 5 μM AldO on a 100-mL reaction scale. The reaction was quenched by adding 0.5 mL 10 M HCl after 6 h reaction time at 30 °C and then extracted with equal volume of ethyl acetate for three times. The organic extracts were combined and concentrated by vacuum rotary evaporation and re-dissolved in 2 mL acetonitrile (ACN) as the working sample for semi-preparative HPLC purification on a Waters XBridgeTM C-18 column (10 × 250 mm, 5 μm).

For preparation of compound **15**-P1: 37% ACN-63% $H_2O$ over 35 min at a flow rate of 2.5 mL/min ($\lambda$ = 220 nm, retention time: 23.5 min).

For preparation of compound **16**-P1: 48% ACN-52% $H_2O$ over 35 min at a flow rate of 2.5 mL/min ($\lambda$ = 220 nm, retention time: 16.6 min).

For preparation of compound **17**-P1: 48% ACN-52% $H_2O$ over 35 min at a flow rate of 2.5 mL/min ($\lambda$ = 220 nm, retention time: 23.8 min).

For preparation of compound **22**-P2: 50% ACN-50% $H_2O$ over 30 min at a flow rate of 2.5

mL/min ($\lambda$ = 220 nm, retention time: 16.1 min).

For preparation of compound **24**-P2: 25% ACN-75% $H_2O$ over 30 min at a flow rate of 2.5 mL/min ($\lambda$ = 220 nm, retention time: 16.5 min).

**1.6 Protein expression, purification, crystallization and structure determination**

The constructs were cloned using standard PCR methods and assembled via Gibson assembly. The gene was inserted into a pET-28a vector, while incorporating a 6 × His tag along with a SUMO protease site at the *N*-terminus of $P450_{BS\beta}$-F46A to facilitate purification. Subsequently, the error-free plasmids were isolated and transformed into *E. coli* BL21(DE3) competent cells. The cultures were incubated with shaking at 200 rpm at 37 °C until $OD_{600}$ reached to 0.8, then induced for protein expression with 0.5 mM IPTG and 10 mM 5-ALA at 18 °C for 15 h. Bacterial cells were then collected by centrifugation. The cell pellet from a 1 L culture was resuspended in 30 mL lysis buffer containing 50 mM Tris-HCl (pH 8.0) and 150 mM NaCl, 20 μg $mL^{-1}$ DNase I and 1 mM PMSF, and lysed using a high-pressure cell disruptor (Union-Biotech co, LTD, shanghai, China). For purification of $P450_{BS\beta}$-F46A, the supernatant was incubated with 2 mL Ni-NTA resin at 4 °C for 2 h. The resin was then washed with 50 mL buffer containing 50 mM Tris-HCl (pH 8.0), 150 mM NaCl, and 20 mM imidazole. The SUMO tag was removed with purified His-tagged ULP1 protease at room temperature for 2 h. The protein was further purified using size-exclusion chromatography (SEC) on an ÄKTA FPLC system equipped with a Superdex 200 Increase 10/300 GL column (GE Healthcare Life Sciences). The purity of the main peak protein sample was analyzed by SDS-PAGE, followed by aliquoting and flash-freezing with liquid nitrogen for storage at -80 °C.

The crystallization screening of $P450_{BS\beta}$-F46A (5 mg $mL^{-1}$) was conducted using the hanging-drop vapor diffusion method at 20 °C, with a 1:1 mixture of 1 μL protein and 1 μL reservoir solution. Crystals were successfully obtained in a solution containing 0.45 M $Li_2SO_4$, 50 mM MES (pH 5.8), and 10% PEG3350. Prior to data collection, the crystals were cryo-protected by immersing them in a drop of reservoir solution supplemented with 25% glycerol. X-ray diffraction experiments were performed at the BLl9U beamline of the Shanghai Synchrotron Radiation Facility (SSRF, China) under cryogenic conditions at 100 K. Data indexing, integration, and scaling were carried out using HKL3000.[5] The phase of $P450_{BS\beta}$-F46A was determined by molecular replacement using PHENIX[6] with the $P450_{BS\beta}$ wild-type structure (PDB accession code:1IZO) as the searching template. The model was manually built with Coot[7]. The structure was refined with PHENIX and manual checking was performed between refinement cycles. The final data collection and refinement statistics were summarized in Supplementary Table 7. All structural figures were generated using PyMOL version 2.3.2 (https://pymol.org/2).

##### 1.7 AutoDock analysis

The initial structures of **1** and **19** were generated using ChemDraw 22.0, and subsequently optimized through density functional theory (DFT). Automated molecular docking was performed using the DFT-optimized structures with the AutoDock Vina program (version 1.5.6) integrated with AutoDockTools (ADT).[8] Receptor-ligand complexes preparation followed the guidelines provided in AutoDock manual, with polar hydrogens added using AutoDockTools. All side chains remained fixed during docking calculations while a suitable grid enclosing heme's active site was defined through the model visualization in AutoDockTools. Default parameter settings along with top-scoring docked conformations served as input for subsequent analyses in this study. Visualization and analysis of resulting docked models employed ADT alongside PyMOL version 2.3.2 (https://pymol.org/2/). The substrate access channels were determined using the Caver 3.0.3 plugin in PyMOL 2.3.2.[9]

##### 1.8 MD, QM/MM and QM calculations

The initial 3D structure of $P450_{BS\beta}$-F46A, crystallized in the present study, was prepared based on the X-ray structure (PDB accession code: 9IY1) for MD, QM/MM, and QM calculations. Herein, we assigned the protonation states of titratable residues (His, Glu, Asp) on the basis of pKa values from the PROPKA software[10] in combination with careful visual inspection of local hydrogen-bond networks. The force field of Fe(III)-$H_2O_2$ complex was parameterized using the "MCPB.py" modeling tool[11] of AmberTools18[12]. The Amber ff14SB force field[13] was employed for the protein residues. The general AMBER force field (GAFF)[14] was used for substrates, while the partial atomic charges were obtained from the RESP method[15], using HF/6-31G* level of theory. The parmchk utility from AmberTools18 was used to generate the missing parameters for substrates and co-substrates. Sodium ions were added to the protein surface to neutralize the total charge of the systems. Finally, the resulting three systems were solvated in a rectangular box of TIP3P waters extending up to a minimum distance of 15 Å from the protein surface. After setup, the whole system was fully minimized using combined steepest descent and conjugate gradient method. Then, the system was gently annealed from 10 to 300 K under canonical ensemble for 50 ps with a weak restraint of 15 kcal/mol/Å on protein. To achieve a uniform density after heating dynamics, 1 ns of density equilibration was performed under the NPT ensemble at the target temperature of 300 K and target pressure of 1.0 atm. Afterwards, we removed all restraints on protein and further equilibrated the system for 10 ns under the NPT ensemble to get the well settled pressure and temperature. Finally, a productive MD simulation under the NPT ensemble was conducted for 100-200 ns for the enzyme system. During all MD simulations, the covalent bonds containing hydrogen were constrained using SHAKE and an integration step of 2 fs was used. All MD simulations were performed with GPU version of Amber 18 package[12].

The representative snapshot extracted from each classical MD trajectory was used for the subsequent QM/MM calculations. All QM/MM calculations were performed using ChemShell[16], combining turbomole[17] for the QM region and DL_POLY[18] for the MM region. The electronic embedding scheme[19] was used to account for the polarizing effect of the enzyme environment on the QM region. Hydrogen link atoms with the charge-shift model were applied to treat the QM/MM boundary. During QM/MM geometry optimizations, the QM region was studied with the hybrid UB3LYP[20] density functional with two levels of theory. For geometry optimization, the double-ζ basis set def2-SVP was used. The energies were further corrected with the larger basis set def2-TZVP for all atoms. Dispersion corrections computed with Grimme's D3 method[21] were included in all QM calculations. The QM region in the system included the substrate, the Fe(III)---$H_2O_2$ complex of $P450_{BS\beta}$-F46A, the coordinated S atom of C363 and the side chain of Arg242. All the transition states (TSs) were located by relaxed potential energy surface (PES) scans followed by full TS optimizations using the DL-FIND code[22].

All QM model calculations were performed with the Gaussian 16 software[23]. For the P450-mediated O-O cleavage of $H_2O_2$, the geometries of species were optimized in conjunction with the SMD continuum solvation model (the solvent chlorobenzene is used)[24] at the B3LYP/def2-SVP level of theory, while the energies were further refined with the larger basis set def2-TZVP for all atoms. As for the geometries of ene-ketone intermediate species, they were fully optimized in conjunction with the SMD continuum solvation model at the B3LYP/6-31G(d) level of theory. The energies were further refined with the larger basis set 6-311++g(d,p) for all atoms. Dispersion corrections computed with Grimme's D3 method[21-22, 25] were included in all QM calculations.

##### 1.9 Analytical methods

**LC-MS analysis.** LC-MS analysis was performed on a YMC HPLC column (150 × 4.6 mm, $C_{18}$) using the negative mode electrospray ionization with a linear gradient of 10% to 100% ACN in dd$H_2O$ with 0.1% formic acid at a flow rate of 1 mL/min. The high-resolution mass spectra (HRMS) were recorded on a Bruker Maxis UHR-TOF.

**GC analysis.** The samples (*e.g.*, **3**, **4**, **7**) were analyzed by the methods reported previously[26]. The Agilent 7890B gas chromatograph equipped with a capillary column HP-5 (Agilent Technologies, Santa Clara, CA, USA; cross-linked polyethylene glycerol, i.d. 0.25 μm film thickness, 30 m by 0.32 mm) was used for GC analyses. The oven program was set initially at 40 °C for 4 min, then increased to 280 °C by the rate of 10 °C per min and held for 5 min. The injecting temperature was set to 280 °C under splitless injection conditions with 1 μL injection volume.

**NMR analysis.** The nuclear magnetic resonance (NMR) spectra were recorded on a Bruker Avance III 600 MHz spectrometer.

### 2. Results

#### 2.1 Supporting tables

**Table S1.** Primers used in this study.

| **Primer** | **Sequence (5'→3')** |
|---|---|
| P450$_{BS\beta}$-F | GGAATTC**CATATG**GCTAGCATGAATGAGCAG |
| P450$_{BS\beta}$-R | CCG**CTCGAG**TTAACTTTTTCGTCTGATTCCG |
| P450$_{BS\beta}$-F46A-F | AAAAACGCAATTTGCATGACTGGC |
| P450$_{BS\beta}$-F46A-R | GCAAATTGCGTTTTTTCCCAACAA |
| P450$_{BS\beta}$-F79A-F | TCGCTGGCAGGTGTTAATGCGATT |
| P450$_{BS\beta}$-F79A-R | AACACCTGCCAGCGATTTCTGCAC |
| P450$_{BS\beta}$-F173A-F | GACGCGGCAGGTGCTGTGGGACCG |
| P450$_{BS\beta}$-F173A-R | AGCACCTGCCGCGTCGACCATGTC |
| P450$_{BS\beta}$-F289A-F | TATCCGGCAGGCCCGTTTTTAGGG |
| P450$_{BS\beta}$-F289A-R | CGGGCCTGCCGGATAATATCTGCG |
| P450$_{BS\beta}$-F292A-F | GGCCCGGCATTAGGGGCGCTTGTC |
| P450$_{BS\beta}$-F292A-R | CCCTAATGCCGGGCCGAACGGATA |
| P450$_{BS\beta}$-F46A-R242A-F | GTACTGGCACCTATTGTCGCCATTTCT |
| P450$_{BS\beta}$-F46A-R242A-R | AATAGGTGCCAGTACATTAATCAGCTC |
| P450$_{BS\beta}$-F46A-R242E-F | GTACTGGAACCTATTGTCGCCATTTCT |
| P450$_{BS\beta}$-F46A-R242E-R | AATAGGTTCCAGTACATTAATCAGCTC |
| P450$_{BS\beta}$-F46A-R242S-F | GTACTGTCACCTATTGTCGCCATTTCT |
| P450$_{BS\beta}$-F46A-R242S-R | AATAGGTGACAGTACATTAATCAGCTC |
| P450$_{BS\beta}$-F46A-R242K-F | GTACTGAAGCCTATTGTCGCCATTTCT |
| P450$_{BS\beta}$-F46A-R242K-R | AATAGGCTTCAGTACATTAATCAGCTC |
| AldO-F | GGAATTC**CATATG**AGCGATATTACCGTGACC |
| AldO-R | CCG**CTCGAG**TTAACCCGCTAACACGCCACGC |

Note: The bold nucleotides denote the restriction sites of *Nde*I and *Xho*I. The red nucleotides denote the mutated codons and the sequences of complementary bases are underlined.

**Table S2.** [1]H NMR (600 MHz, $CD_3CN$) and [13]C NMR (151 MHz, $CD_3CN$) data of **15**-P1.

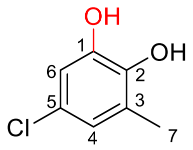


| **Position** | **$\delta_C$, type** | **$\delta_H$, mult., ($J$ in Hz)** |
|---|---|---|
| 1 | 145.8, C | |
| 2 | 142.9, C | |
| 3 | 127.2, C | |
| 4 | 122.3, CH | 6.66, m |
| 5 | 124.0, C | |
| 6 | 113.7, CH | 6.70, d, (2.2) |
| 7 | 15.8, $CH_3$ | 2.15, s |

**Table S3.** [1]H NMR (600 MHz, $CD_3CN$) and [13]C NMR (151 MHz, $CD_3CN$) data of **16**-P1.

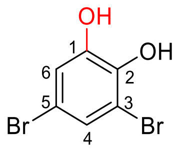


| **Position** | **$\delta_C$, type** | **$\delta_H$, mult., (*J* in Hz)** |
|---|---|---|
| 1 | 147.0, C | |
| 2 | 143.2, C | |
| 3 | 110.7, C | |
| 4 | 126.6, CH | 7.19, d, (2.2) |
| 5 | 112.0, C | |
| 6 | 118.5, CH | 6.98, d, (2.2) |

**Table S4.** $^{1}H$ NMR (600 MHz, $CD_3CN$) and $^{13}C$ NMR (151 MHz, $CD_3CN$) data of **17**-P1.

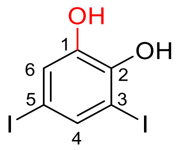


| **Position** | ***δ*$_C$, type** | ***δ*$_H$, mult., (*J* in Hz)** |
|---|---|---|
| 1 | 145.8, C | |
| 2 | 146.7, C | |
| 3 | 84.7, C | |
| 4 | 138.0, CH | 7.54, d, (1.9) |
| 5 | 82.1, C | |
| 6 | 124.9, CH | 7.14, d, (1.9) |

**Table S5.** [1]H NMR (600 MHz, $CD_3CN$) and [13]C NMR (151 MHz, $CD_3CN$) data of **22**-P2.

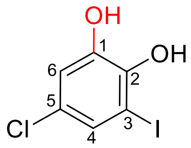


| **Position** | ***δ*<sub></sub>C, type** | ***δ*H, mult., (*J* in Hz)** |
|---|---|---|
| 1 | 145.6, C | |
| 2 | 145.7, C | |
| 3 | 83.7, C | |
| 4 | 129.3, CH | 7.23, d, (2.3) |
| 5 | 125.8, C | |
| 6 | 116.8, CH | 6.89, d, (2.3) |

**Table S6.** [1]H NMR (600 MHz, DMSO-$d_6$) and [13]C NMR (151 MHz, DMSO-$d_6$) data of **24**-P2.

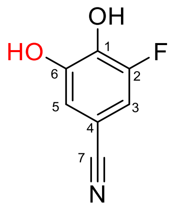


| **Position** | **$\delta_C$, mult., (*J* in Hz),** | **$\delta_H$, mult., (*J* in Hz)** |
|---|---|---|
| 1 | 138.0, d, (13.6), C | |
| 2 | 150.1, d, (239.8), C | |
| 3 | 110.4, d, (22.6), CH | 7.24, d, (11.5) |
| 4 | 98.6, d, (11.9), C | |
| 5 | 113.9, s, CH | 6.98, s |
| 6 | 147.0, d, (6.7), C | |
| 7 | 117.5, d, (3.2), C | |

**Table S7.** Data collection and refinement statistics.

| | P450$_{BS\beta}$-F46A |
|---|---|
| PDB ID | 9IY1 |
| **Data collection** | |
| Wavelength | 0.987 |
| Resolution range | 49.49 - 2.29 (2.372 - 2.29) |
| Space group | I222 |
| a b c | 172.42 190.114 226.571 |
| α β γ | 90.00 90.00 90.00 |
| Unique reflections | 165978 (16442) |
| Completeness (%) | 97.45 (93.20) |
| Mean I/sigma(I) | 1.47 |
| Wilson B-factor | 28.37 |
| R-means | 0.306(1.488) |
| CC1/2 | 0.983(0.636) |
| Data redundancy | 12.9(12.0) |
| **Refinement** | |
| Resolution range | 49.49 - 2.29 (2.372 - 2.29) |
| Reflections used in refinement | 162139 |
| Reflections used for R-free | 1997 |
| R-work | 0.1978 (0.2519) |
| R-free | 0.2403 (0.2949) |
| Number of non-hydrogen atoms | 21795 |
| macromolecules | 20192 |
| ligands | 651 |
| solvent | 952 |
| Protein residues | 2489 |
| RMS (bonds) | 0.004 |
| RMS (angles) | 0.57 |
| Ramachandran favored (%) | 98.14 |
| Ramachandran allowed (%) | 1.61 |
| Ramachandran outliers (%) | 0.24 |
| Rotamer outliers (%) | 0.52 |
| Clash score | 3.77 |
| Average B-factor | 32.06 |
| macromolecules | 31.76 |
| ligands | 39.40 |
| solvent | 33.40 |

Values in parentheses are for highest-resolution shell.

**Table S8.** The catalytic activities of P450$_{BS\beta}$ and P450$_{BS\beta}$-F46A towards non-carboxylic substrates.

| Enzyme | Substrate | mM | TON | Conversion/% | Reaction types |
|---|---|---|---|---|---|
| P450$_{BS\beta}$ | **1** | 0.5 | 121 ± 17 | 24.2 | hydroxylation |
| | **16** | 0.5 | 168 ± 12 | 33.5 | hydroxylation |
| | **17** | 0.5 | 75 ± 8 | 15.0 | hydroxylation |
| | **18** | 0.5 | 296 ± 40 | 59.2 | hydroxylation |
| | **19** | 0.5 | 67 ± 9 | 13.5 | dehalogenation/hydroxylation |
| | **20** | 0.5 | 15 ± 1 | 3.0 | dehalogenation/hydroxylation |
| P450$_{BS\beta}$-F46A | **11** | 0.5 | 257 ± 30 | 51.4 | hydroxylation |
| | **12** | 0.5 | 262 ± 7 | 52.4 | hydroxylation |
| | **13** | 0.5 | 153 ± 11 | 30.6 | hydroxylation |
| | **14** | 0.5 | 41 ± 6 | 8.3 | hydroxylation |
| | **15** | 0.5 | 88 ± 6 | 17.6 | hydroxylation |
| | **1** | 0.5 | 479 ± 21 | 95.9 | hydroxylation |
| | | 1 | 875 ± 10 | 87.5 | |
| | | 2 | 1607 ± 139 | 80.3 | |
| | | 5 | 917 ± 17 | 18.4 | |
| | **16** | 0.5 | 450 ± 18 | 90.0 | hydroxylation |
| | | 1 | 875 ± 29 | 87.5 | |
| | | 2 | 1522 ± 93 | 76.1 | |
| | | 5 | 489 ± 49 | 9.8 | |
| | **17** | 0.5 | 380 ± 28 | 76.1 | hydroxylation |
| | | 1 | 593 ± 54 | 59.3 | |
| | | 2 | 259 ± 12 | 13.0 | |
| | **18** | 0.5 | 483 ± 1 | 96.6 | hydroxylation |
| | | 1 | 975 ± 11 | 97.5 | |
| | | 2 | 1832 ± 25 | 91.6 | |
| | | 5 | 892 ± 126 | 17.8 | |
| | **19** | 0.5 | 295 ± 2 | 58.9 | dehalogenation/hydroxylation |
| | **20** | 0.5 | 173 ± 19 | 34.6 | dehalogenation/hydroxylation |
| | **21** | 0.5 | 160 ± 5 | 31.9 | dehalogenation/hydroxylation |
| | **22** | 0.5 | 192 ± 4 | 38.4 | dehalogenation/hydroxylation |
| | **23** | 0.5 | 276 ± 4 | 55.1 | hydroxylation |
| | **24** | 0.5 | 315 ± 19 | 62.9 | hydroxylation |
| | **25** | 0.5 | 242 ± 18 | 48.3 | hydroxylation |

Reaction conditions: 1 μM P450 (P450$_{BS\beta}$ or P450$_{BS\beta}$-F46A), 0.5-5 mM substrates, and 5 μM AldO + 10% glycerol as *in situ* $H_2O_2$ releasing system, at 30 °C for 6 h. All experiments were performed in triplicate.

**Table S9.** The substrate dissociation constants ($K_D$) of non-carboxylic substrates with P450$_{BS\beta}$, P450$_{BS\beta}$-F46A and P450$_{BS\beta}$-F46A-R242A.

| Enzyme | Substrate | $K_D$/μM | $\Delta A_{max}$/$\mu M^{-1}\cdot cm^{-1}$ | Ref |
|---|---|---|---|---|
| P450$_{BS\beta}$ | lauric acid | 1.8 ± 0.2 | 0.024 ± 0.001 | [1b] |
| | **1** | 73.2 ± 26.6 | 0.018 ± 0.002 | this study |
| P450$_{BS\beta}$-F46A | lauric acid | 4.0 ± 0.8 | 0.029 ± 0.002 | this study |
| | **1** | 7.6 ± 2.8 | 0.016 ± 0.001 | this study |
| | **5** | 2.5 ± 0.7 | 0.014 ± 0.001 | this study |
| | **7** | 2.6 ± 1.3 | 0.012 ± 0.001 | this study |
| | **10** | 2.6 ± 0.8 | 0.012 ± 0.001 | this study |
| | **11** | 2.1 ± 0.5 | 0.011 ± 0.001 | this study |
| | **12** | 4.2 ± 2.0 | 0.012 ± 0.001 | this study |
| | **13** | 7.1 ± 2.4 | 0.012 ± 0.001 | this study |
| | **14** | 4.0 ± 1.0 | 0.012 ± 0.001 | this study |
| | **15** | 4.6 ± 2.1 | 0.019 ± 0.002 | this study |
| | **16** | 10.5 ± 3.4 | 0.020 ± 0.002 | this study |
| | **17** | 10.9 ± 2.7 | 0.023 ± 0.001 | this study |
| | **18** | 12.9 ± 4.3 | 0.024 ± 0.002 | this study |
| | **19** | 1.2 ± 0.3 | 0.010 ± 0.001 | this study |
| | **20** | 4.5 ± 1.3 | 0.016 ± 0.001 | this study |
| P450$_{BS\beta}$-F46A-R242A | lauric acid | 14.7 ± 2.7 | 0.027 ± 0.001 | this study |
| | **1** | 19.6 ± 5.8 | 0.045 ± 0.004 | this study |
| | **19** | 3.3 ± 1.3 | 0.021 ± 0.002 | this study |

All experiments were performed in triplicate.

#### 2.2 Supporting figures

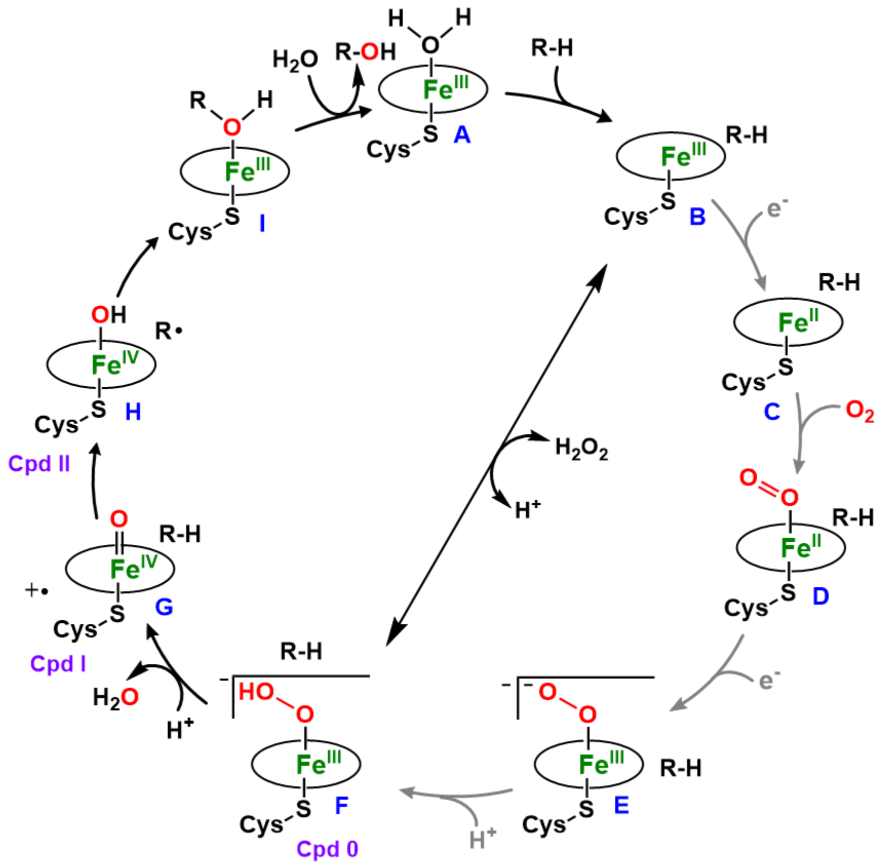


**Figure S1.** P450 catalytic cycle. The peroxide shunt pathway of P450s is indicated by black arrows. The grey arrows indicate the normal $O_2$-activation steps of classical P450 monooxygenases.

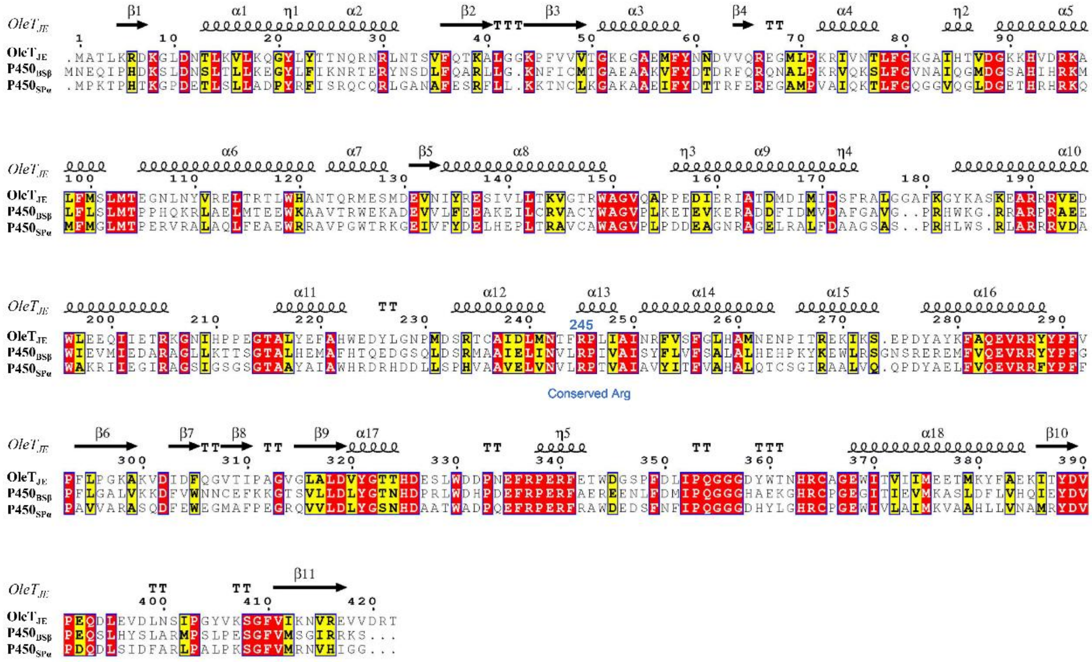


**Figure S2.** Protein sequence alignment of the fatty acid decarboxylase OleT$_{JE}$ from *Jeotgalicoccus sp.* ATCC 8456 (GenBank accession number: ADW41779.1), the fatty acid $\beta$-hydroxylase P450$_{BS\beta}$ from *Bacillus subtilis* (GenBank accession number: NP_388092.1), and the fatty acid $\alpha$-hydroxylase P450$_{SP\alpha}$ from *Sphingomonas paucimobilis* (GenBank accession number: WP_017980797.1). The conserved arginine (R245 for OleT$_{JE}$, R242 for P450$_{BS\beta}$, R241 for P450$_{SP\alpha}$) is crucial for interacting with the substrate carboxylate[27]. P450$_{SP\alpha}$ and P450$_{BS\beta}$ show 37% and 41% amino acid sequence identity to OleT$_{JE}$, respectively.

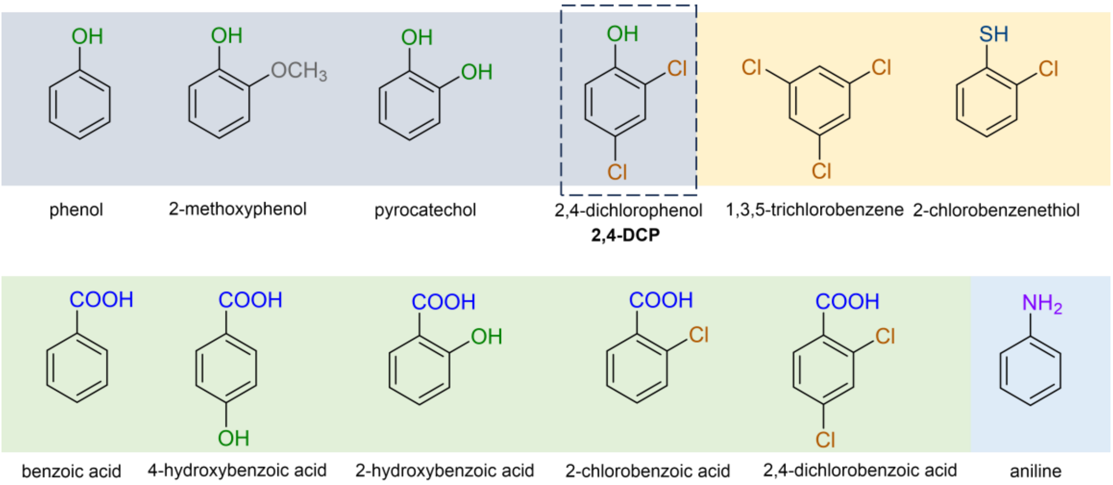


**Figure S3.** Selected benzene derivatives used as potential modulators for the catalytic activity of CYP152 peroxygenases.

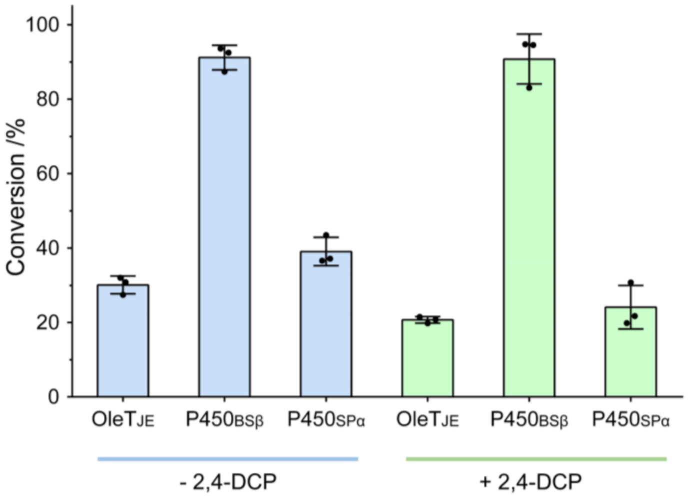


**Figure S4.** The activities of three representative CYP152 peroxygenases towards myristic acid ($C_{14}$) in the presence and absence of 0.5 mM 2,4-dichlorophenol (2,4-DCP, **1**). Reaction conditions: 1 μM P450, 2 mM myristic acid, with or without 0.5 mM **1**, and the AldO/glycerol system to generate $H_2O_2$ as cofactor at 30 °C for 6 h.

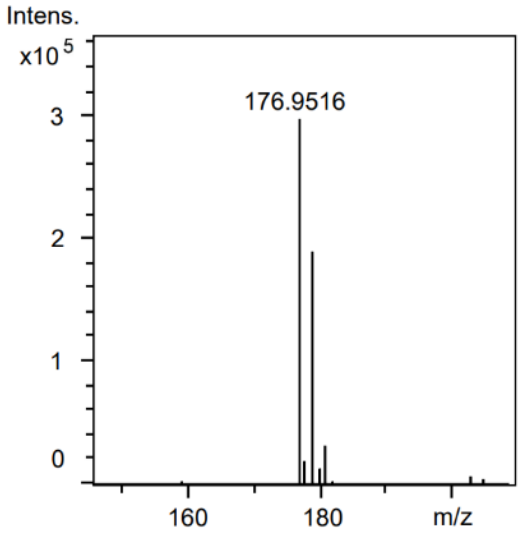


**Figure S5.** HR-ESI-MS of the product of the reaction of **1** catalyzed by $P450_{BS\beta}$ ([M-H]$^-$: *calc.* 176.9516; *obs.* 176.9516).

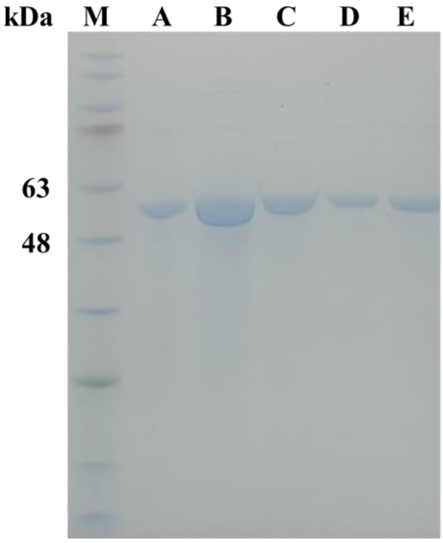


**Figure S6.** SDS-PAGE analysis of the purified His$_6$-tagged P450$_{BS\beta}$-F46A (lane A), P450$_{BS\beta}$-F79A (lane B), P450$_{BS\beta}$-F173A (lane C), P450$_{BS\beta}$-F289A (lane D), P450$_{BS\beta}$-F292A (lane E), and protein marker (M).

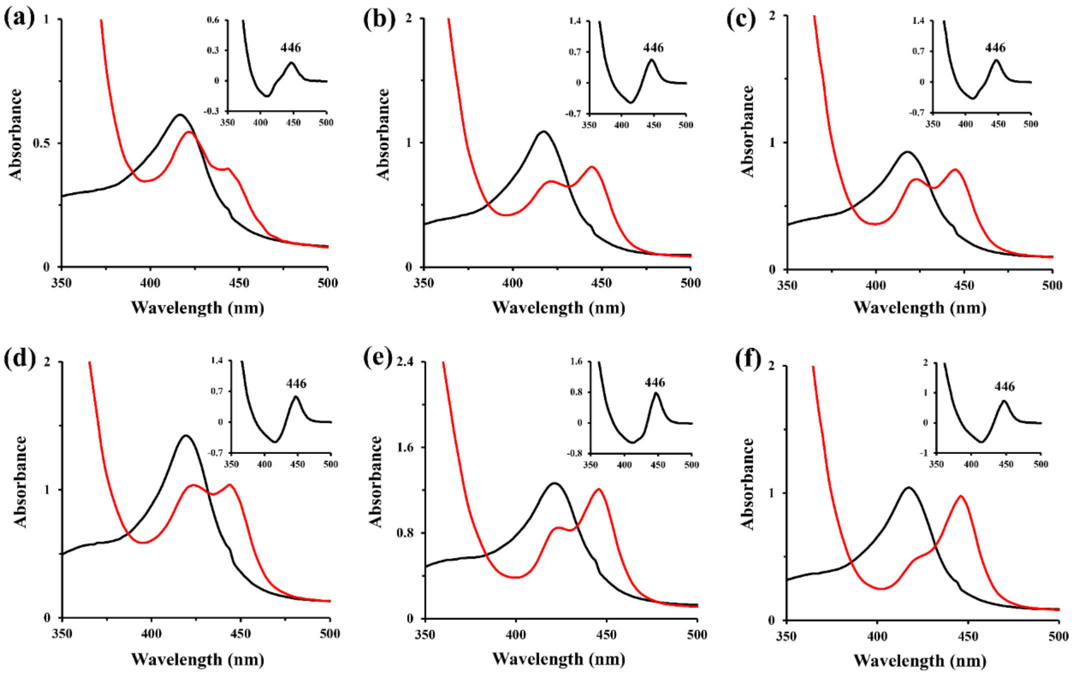


**Figure S7.** UV-visible spectra of P450$_{BS\beta}$ (**a**), P450$_{BS\beta}$-F46A (**b**), P450$_{BS\beta}$-F79A (**c**), P450$_{BS\beta}$-F173A (**d**), P450$_{BS\beta}$-F289A (**e**) and P450$_{BS\beta}$-F292A (**f**). (Black lines show the spectra for the oxidized ferric form of CYPs and red lines show the spectra for the $Na_2S_2O_4$-reduced ferrous-CO complex of CYPs; Insets exhibit the CO-bound reduced difference spectra of P450 enzymes).

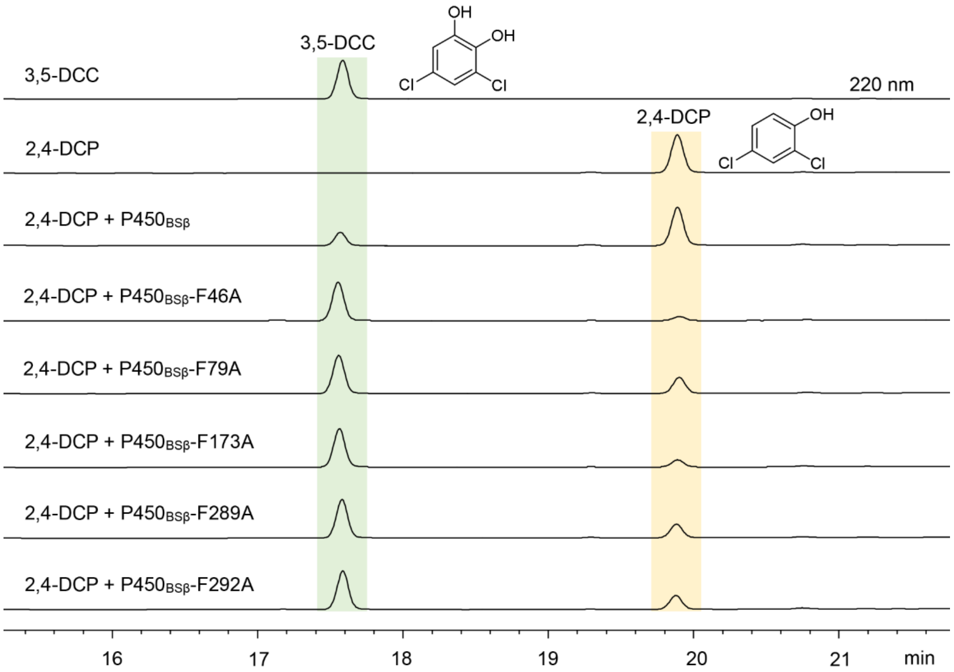


**Figure S8.** HPLC analysis of the reactions of 2,4-dichlorophenol (2,4-DCP, **1**) catalyzed by P450$_{BS\beta}$ mutants and 3,5-dichlorocatechol (3,5-DCC, **2**) standard. Reaction conditions: 1 μM P450, 500 μM 2,4-DCP, and the AldO/glycerol system to generate $H_2O_2$ as cofactor at 30 °C for 6 h.

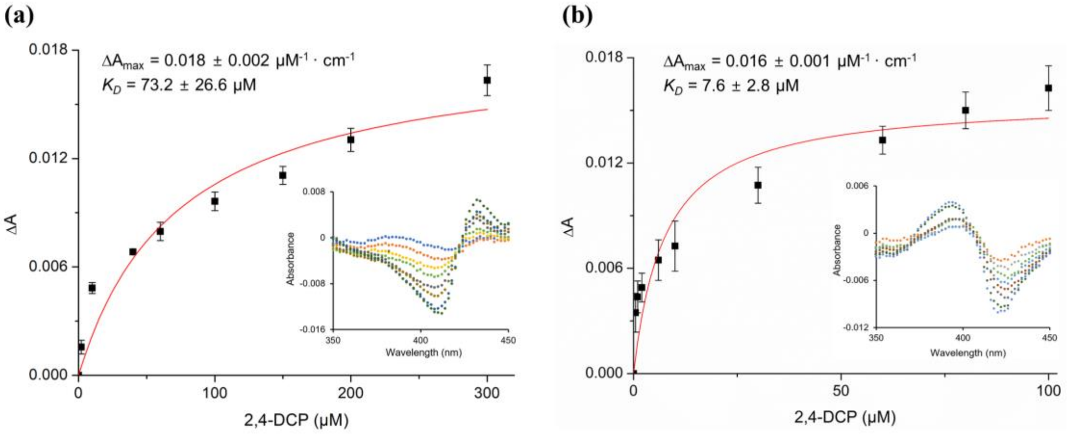


**Figure S9.** Substrate binding curves of P450$_{BS\beta}$ (**a**) and P450$_{BS\beta}$-F46A (**b**) towards 2,4-DCP (**1**). The insets show the Type I difference spectra (**a**) and Type II difference spectra (**b**). All experiments were performed in triplicate.

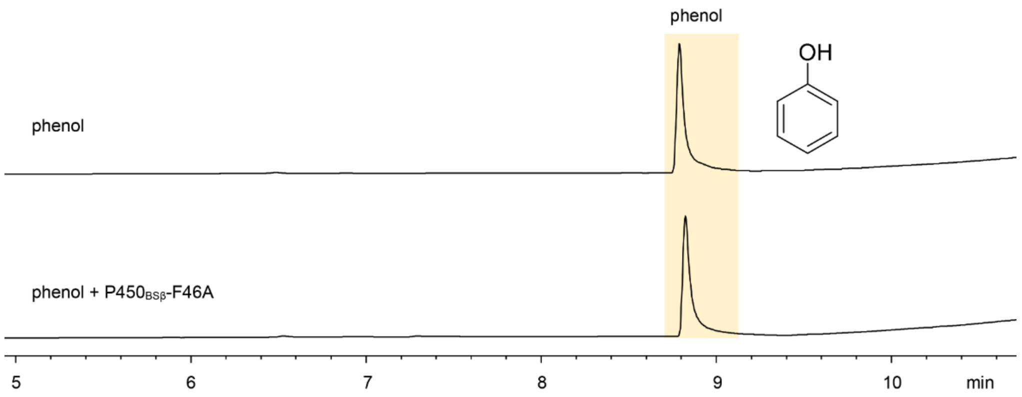


**Figure S10.** GC analysis of the reaction of phenol (**3**) catalyzed by P450$_{BS\beta}$-F46A. Reaction conditions: 1 μM P450, 500 μM substrate, and the AldO/glycerol system to generate $H_2O_2$ as cofactor at 30 °C for 6 h.

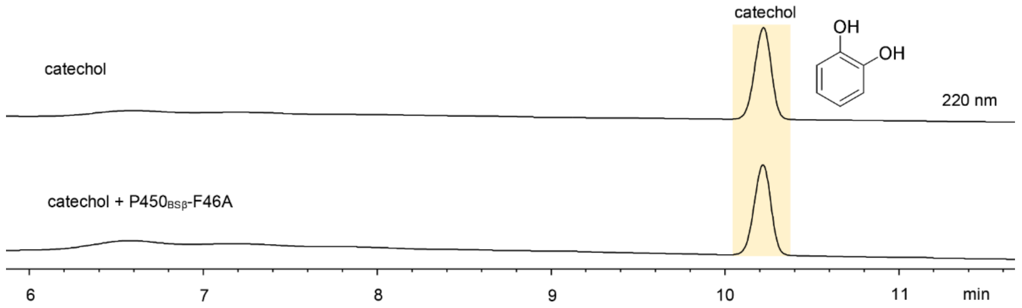


**Figure S11.** HPLC analysis of the reaction of catechol (**4**) catalyzed by P450$_{BS\beta}$-F46A. Reaction conditions: 1 μM P450, 500 μM substrate, and the AldO/glycerol system to generate $H_2O_2$ as cofactor at 30 °C for 6 h.

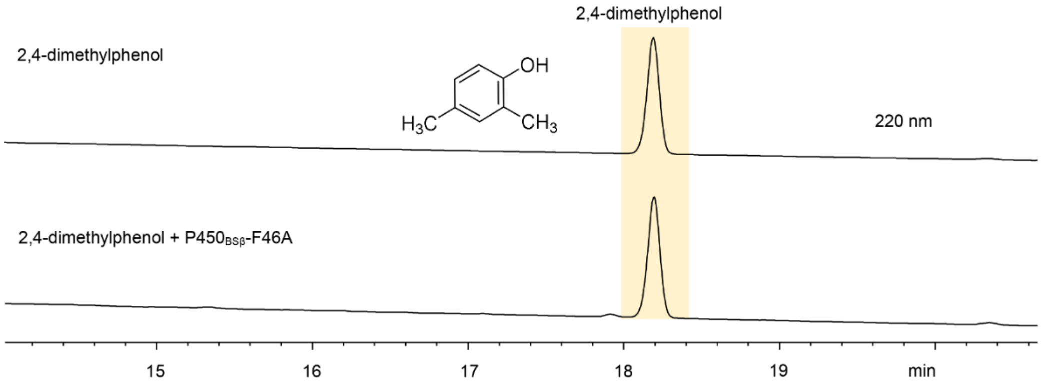


**Figure S12.** HPLC analysis of the reaction of 2,4-dimethylphenol (**5**) catalyzed by P450$_{BS\beta}$-F46A. Reaction conditions: 1 μM P450, 500 μM substrate, and the AldO/glycerol system to generate $H_2O_2$ as cofactor at 30 °C for 6 h.

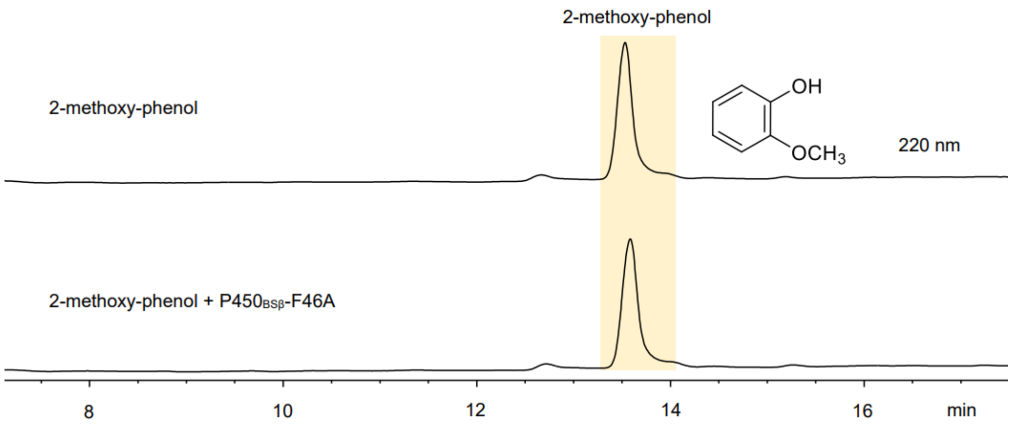


**Figure S13.** HPLC analysis of the reaction of 2-methoxy-phenol (**6**) catalyzed by P450$_{BS\beta}$-F46A. Reaction conditions: 1 μM P450, 500 μM substrate, and the AldO/glycerol system to generate $H_2O_2$ as cofactor at 30 °C for 6 h.

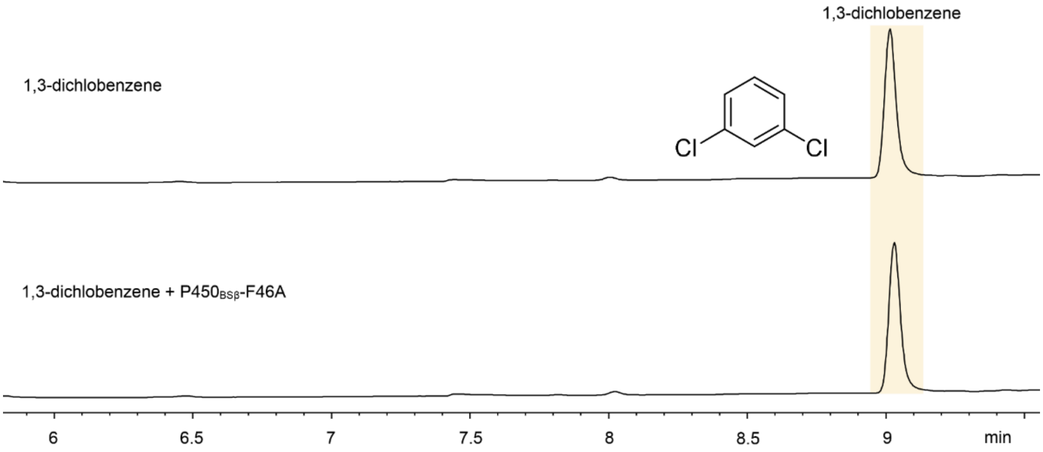


**Figure S14.** GC analysis of the reaction of 1,3-dichlobenzene (**7**) catalyzed by P450$_{BS\beta}$-F46A. Reaction conditions: 1 μM P450, 500 μM substrate, and the AldO/glycerol system to generate $H_2O_2$ as cofactor at 30 °C for 6 h.

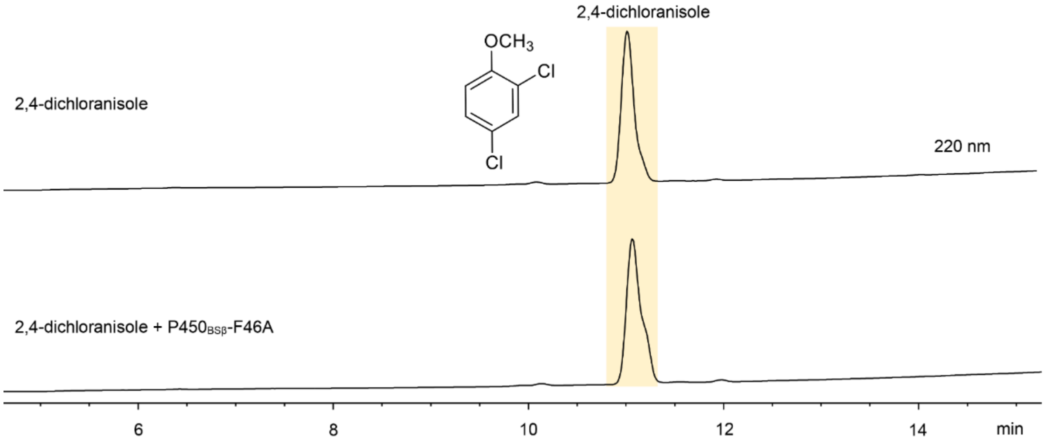


**Figure S15.** HPLC analysis of the reaction of 2,4-dichloranisole (**8**) catalyzed by P450$_{BS\beta}$-F46A. Reaction conditions: 1 μM P450, 500 μM substrate, and the AldO/glycerol system to generate $H_2O_2$ as cofactor at 30 °C for 6 h.

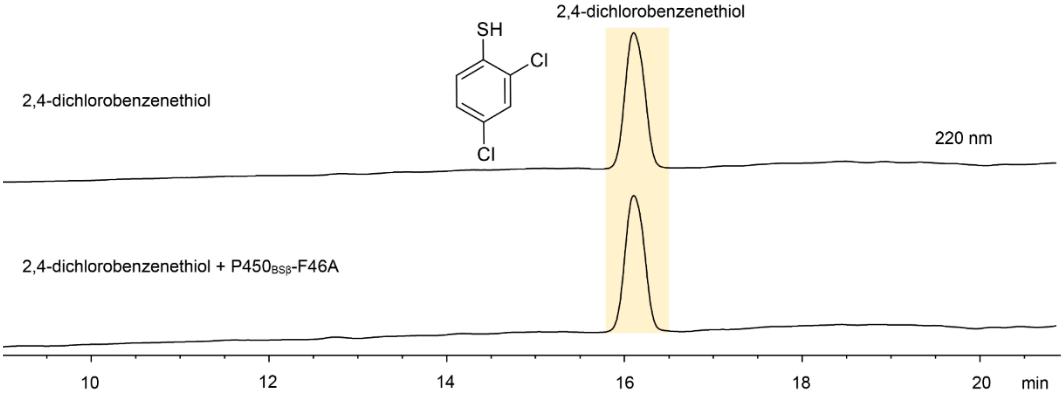


**Figure S16.** HPLC analysis of the reaction of 2,4-dichlorobenzenethiol (**9**) catalyzed by P450$_{BS\beta}$-F46A. Reaction conditions: 1 μM P450, 500 μM substrate, and the AldO/glycerol system to generate $H_2O_2$ as cofactor at 30 °C for 6 h.

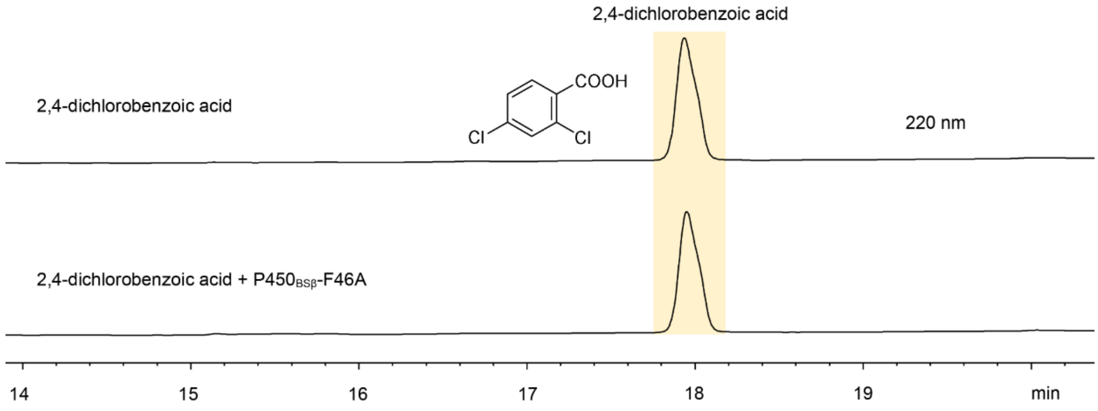


**Figure S17.** HPLC analysis of the reaction of 2,4-dichlorobenzoic acid (**10**) catalyzed by P450$_{BSβ}$-F46A. Reaction conditions: 1 μM P450, 500 μM substrate, and the AldO/glycerol system to generate $H_2O_2$ as cofactor at 30 °C for 6 h.

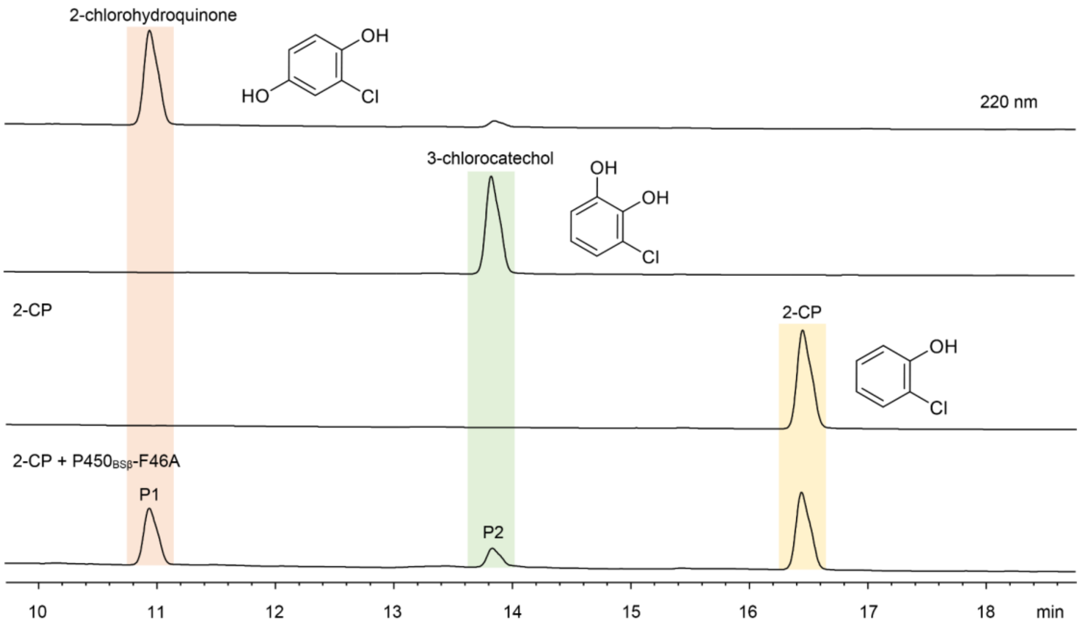


**Figure S18.** HPLC analysis of the reaction of 2-chlorophenol (2-CP, **11**) catalyzed by P450$_{BSβ}$-F46A. Reaction conditions: 1 μM P450, 500 μM substrate, and the AldO/glycerol system to generate $H_2O_2$ as cofactor at 30 °C for 6 h.

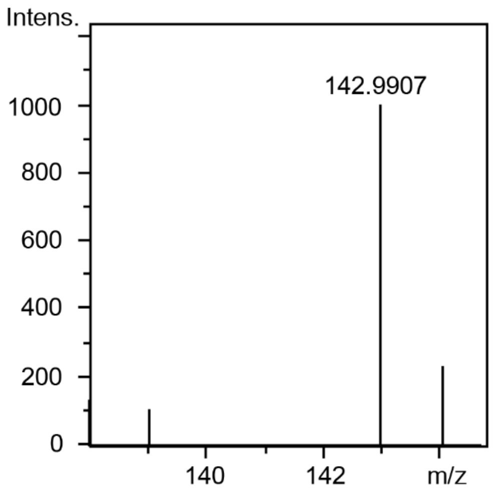


**Figure S19.** HR-ESI-MS of **11**-P1 ([M-H]$^-$: *calc.* 142.9905; *obs.* 142.9907).

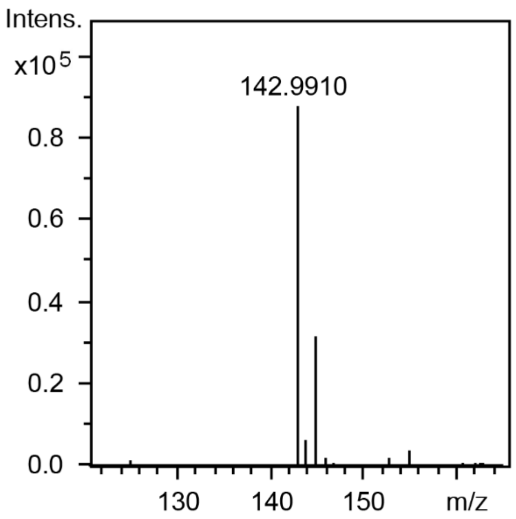


**Figure S20.** HR-ESI-MS of **11**-P2 ([M-H]$^-$: *calc.* 142.9905; *obs.* 142.9910).

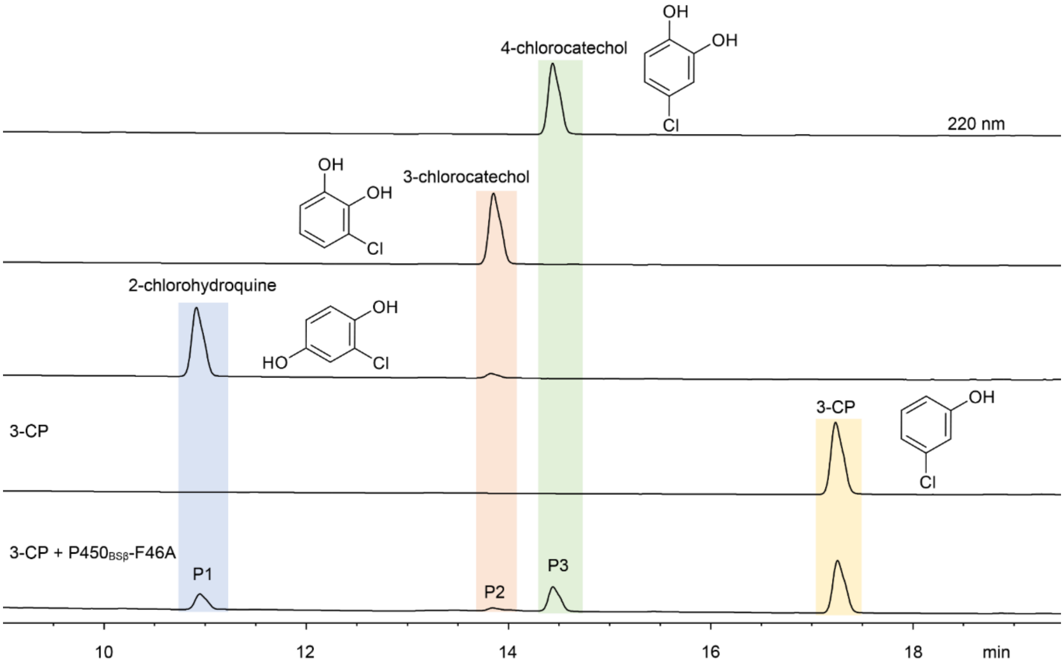


**Figure S21.** HPLC analysis of the reaction of 3-chlorophenol (3-CP, **12**) catalyzed by P450$_{BS\beta}$-F46A. Reaction conditions: 1 µM P450, 500 µM substrate, and the AldO/glycerol system to generate $H_2O_2$ as cofactor at 30 °C for 6 h.

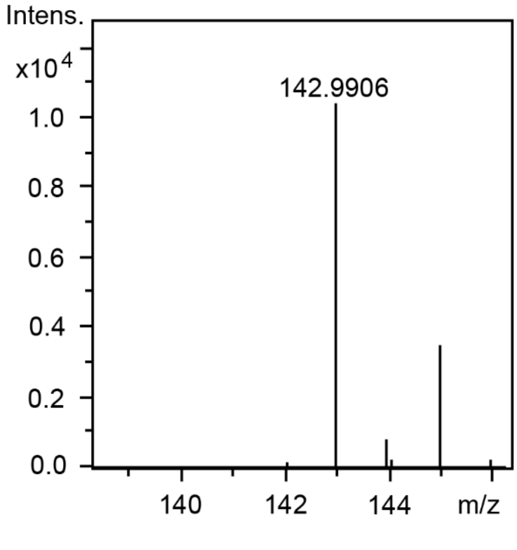


**Figure S22.** HR-ESI-MS of **12-**P1 ([M-H]$^-$: *calc.* 142.9905; *obs.* 142.9906).

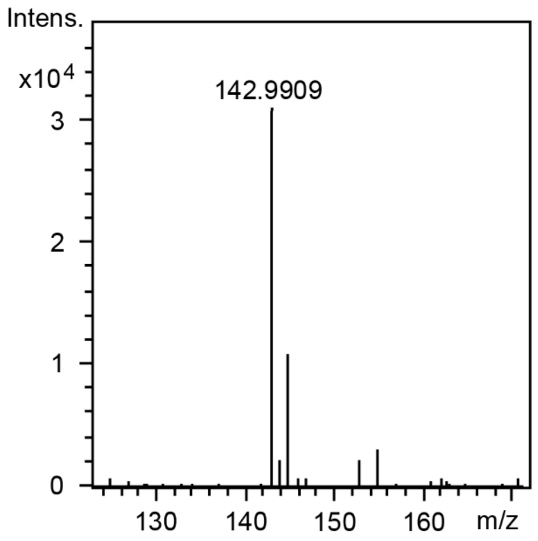


**Figure S23.** HR-ESI-MS of **12**-P2 ([M-H]$^{-}$: *calc.* 142.9905; *obs.* 142.9909).

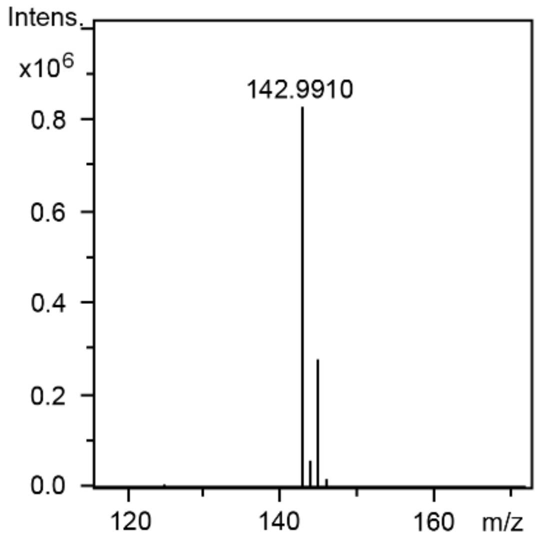


**Figure S24.** HR-ESI-MS of **12**-P3 ([M-H]$^{-}$: *calc.* 142.9905; *obs.* 142.9910).

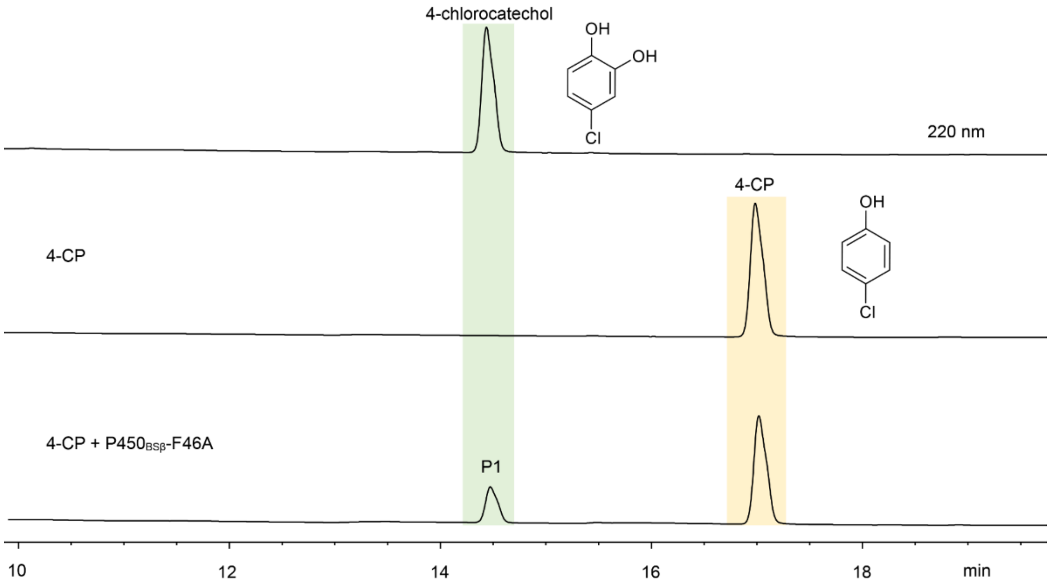


**Figure S25.** HPLC analysis of the reaction of 4-chlorophenol (4-CP, **13**) catalyzed by P450$_{BS\beta}$-F46A. Reaction conditions: 1 µM P450, 500 µM substrate, and the AldO/glycerol system to generate $H_2O_2$ as cofactor at 30 °C for 6 h.

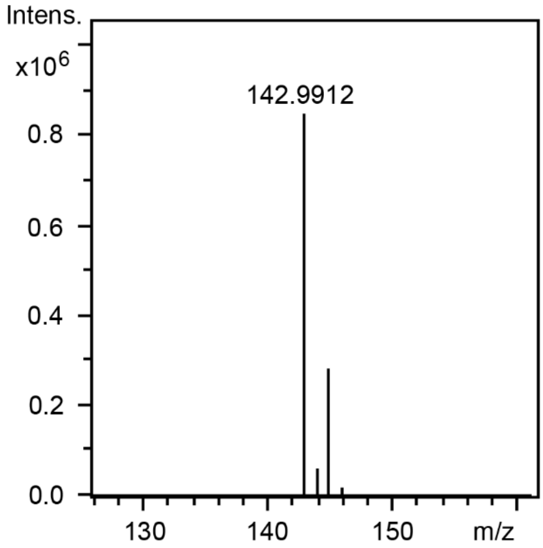


**Figure S26.** HR-ESI-MS of **13-**P1 ($[M-H]^-$: *calc.* 142.9905; *obs.* 142.9912).

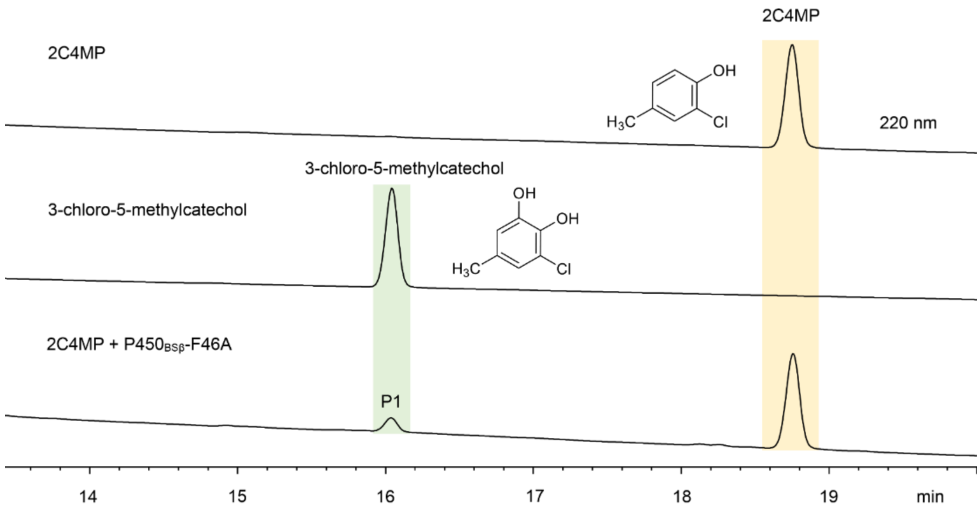


**Figure S27.** HPLC analysis of the reaction of 2-chloro-4-methylphenol (2C4MP, **14**) catalyzed by $P450_{BS\beta}$-F46A and 3-chloro-5-methylcatechol standard. Reaction conditions: 1 µM P450, 500 µM substrate, and the AldO/glycerol system to generate $H_2O_2$ as cofactor at 30 °C for 6 h.

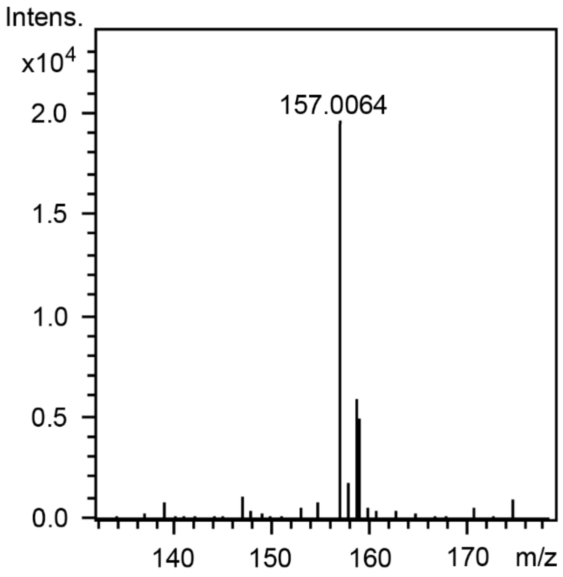


**Figure S28.** HR-ESI-MS of **14**-P1 ($[M-H]^-$: *calc.* 157.0062; *obs.* 157.0064).

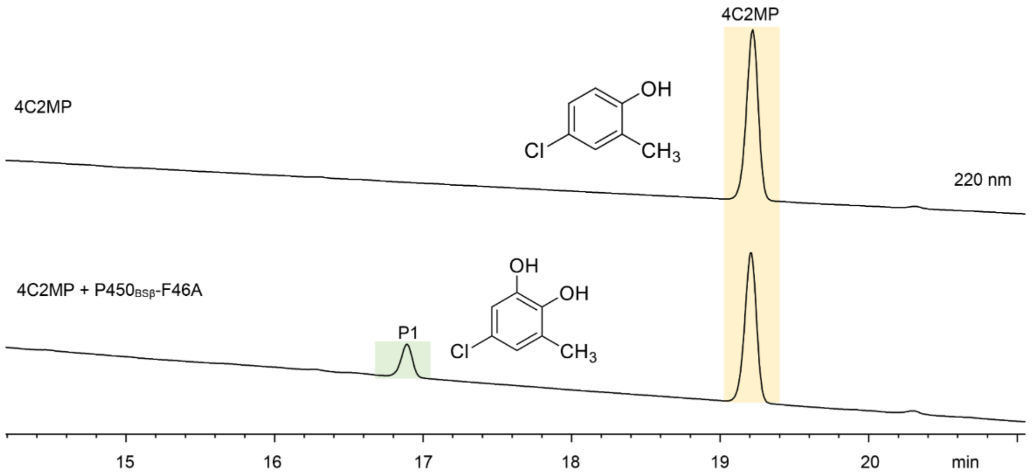


**Figure S29.** HPLC analysis of the reaction of 4-chloro-2-methylphenol (4C2MP, **15**) catalyzed by P450$_{BS\beta}$-F46A. Reaction conditions: 1 μM P450, 500 μM substrate, and the AldO/glycerol system to generate $H_2O_2$ as cofactor at 30°C for 6 h.

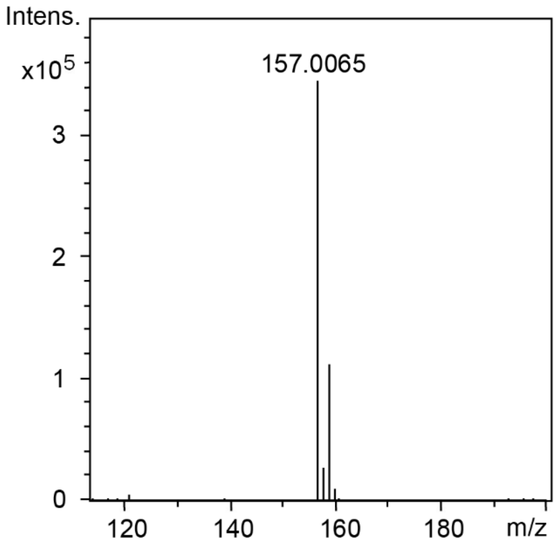


**Figure S30.** HR-ESI-MS of **15-**P1 ([M-H]$^-$: *calc.* 157.0062; *obs.* 157.0065).

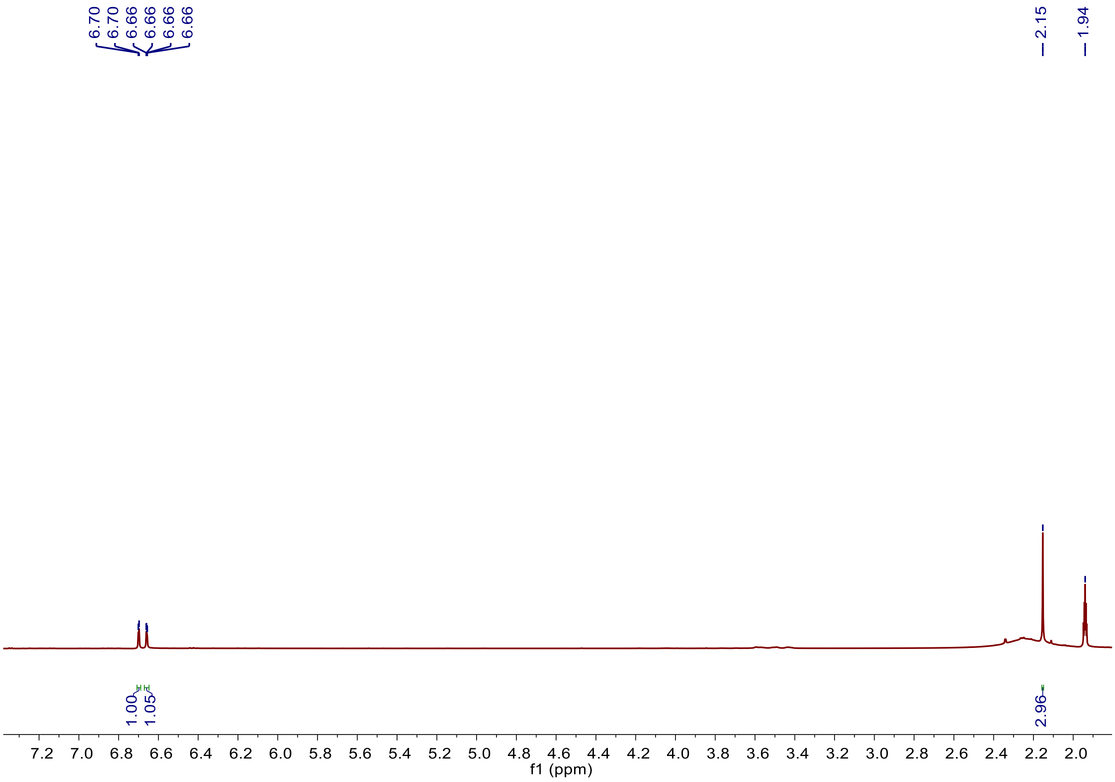


**Figure S31.** $^{1}$H NMR spectrum of **15-**P1 in $CD_3CN$.

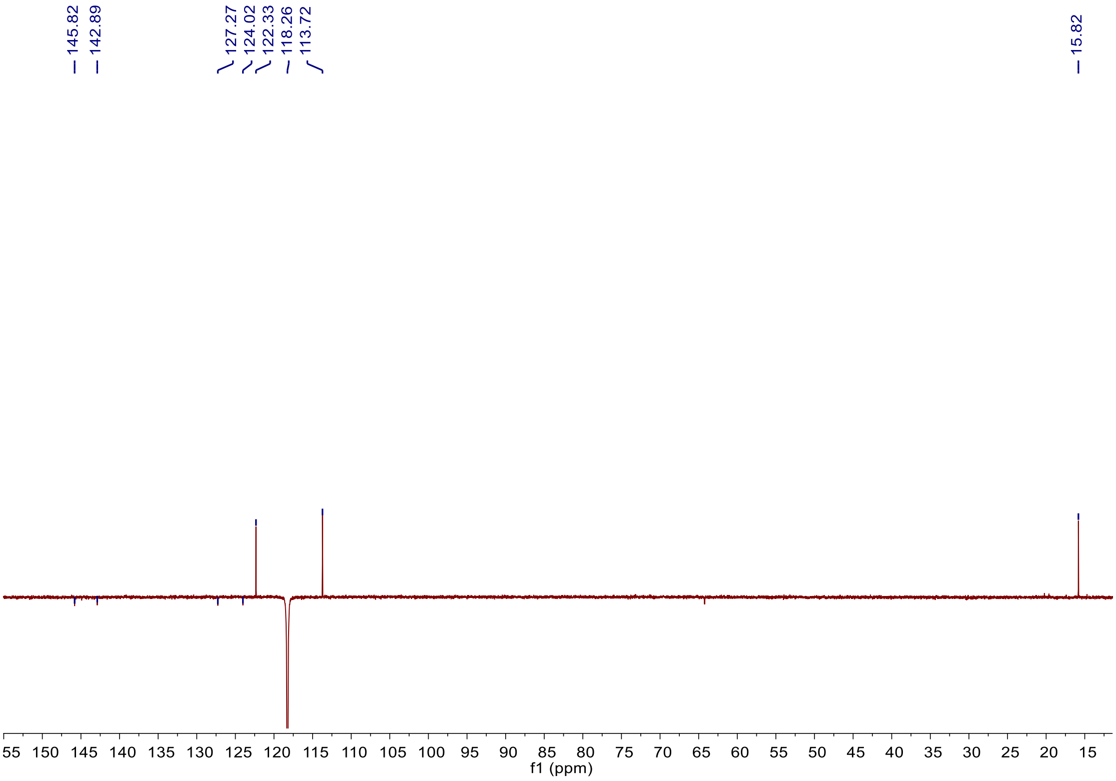


**Figure S32.** $^{13}$C-DEPTQ spectrum of **15-**P1 in $CD_3CN$.

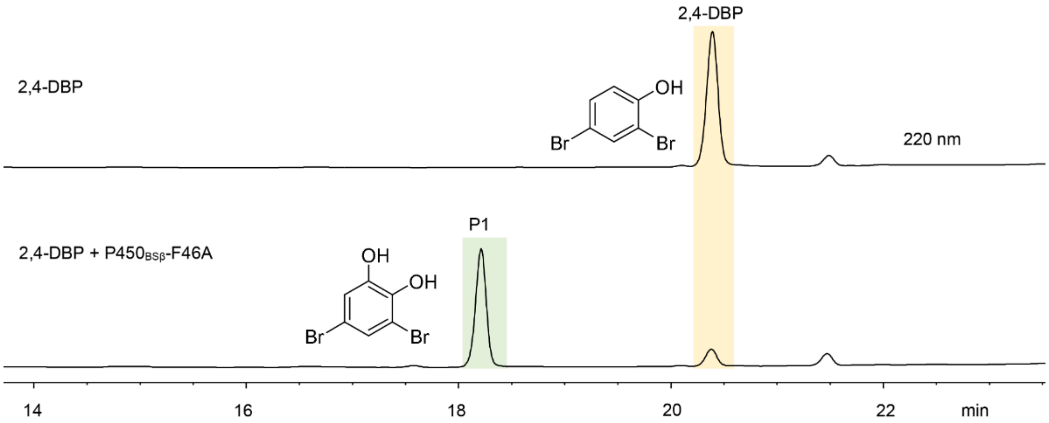


**Figure S33.** HPLC analysis of the reaction of 2,4-dibromophenol (2,4-DBP, **16**) catalyzed by P450$_{BS\beta}$-F46A. Reaction conditions: 1 µM P450, 500 µM substrate, and the AldO/glycerol system to generate $H_2O_2$ as cofactor at 30 °C for 6 h.

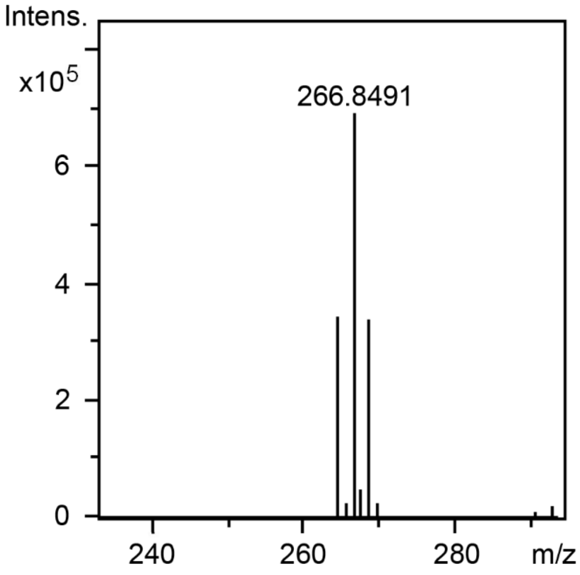


**Figure S34.** HR-ESI-MS of **16**-P1 ([M-H]$^-$: *calc.* 266.8485; *obs.* 266.8491).

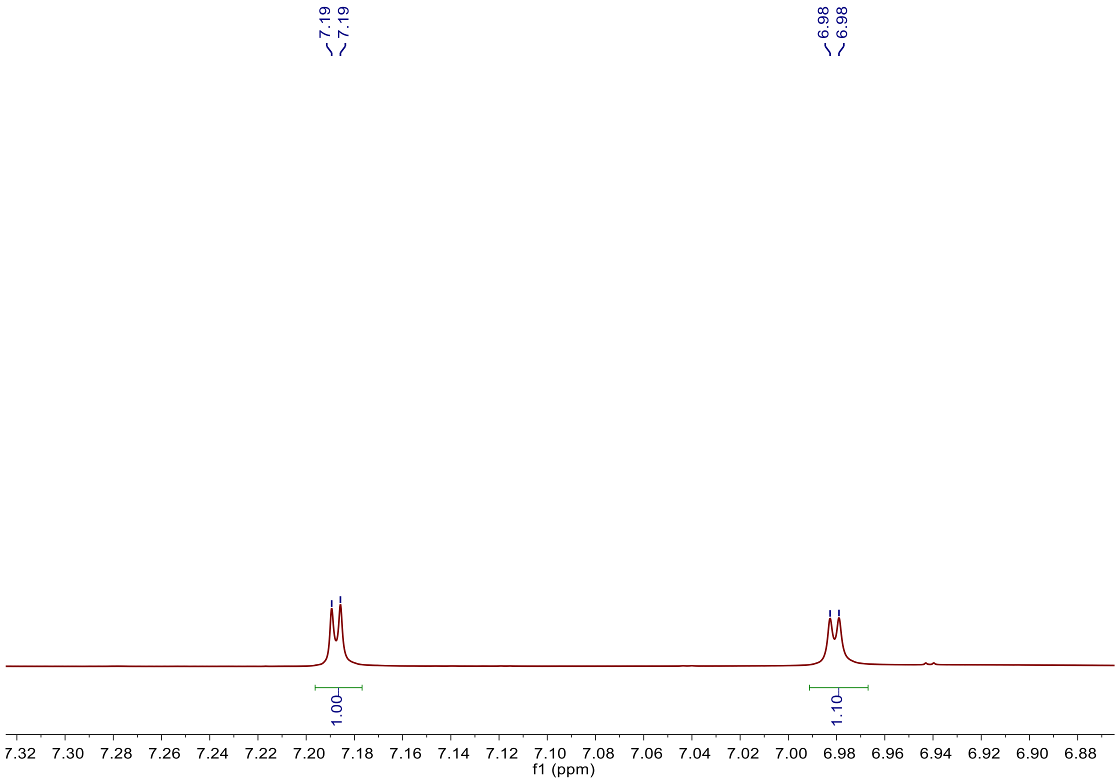


**Figure S35.** $^{1}$HNMR spectrum of **16**-P1 in $CD_3CN$.

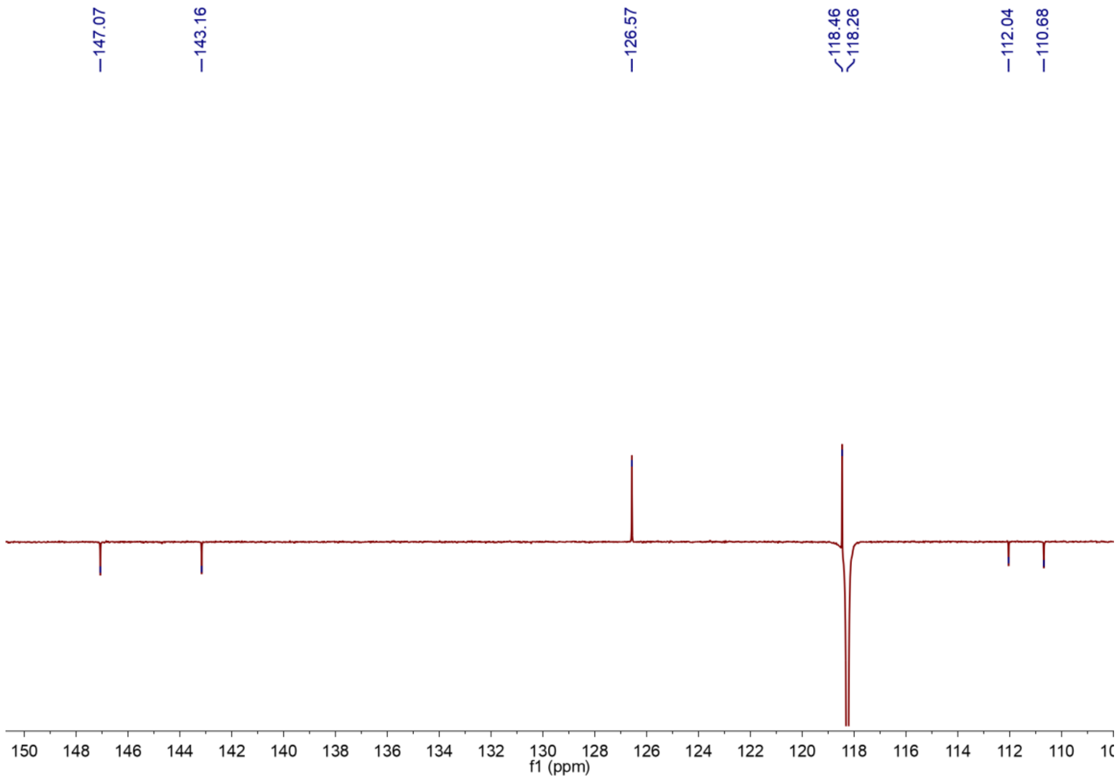


**Figure S36.** $^{13}$C-DEPTQ spectrum of **16**-P1 in $CD_3CN$.

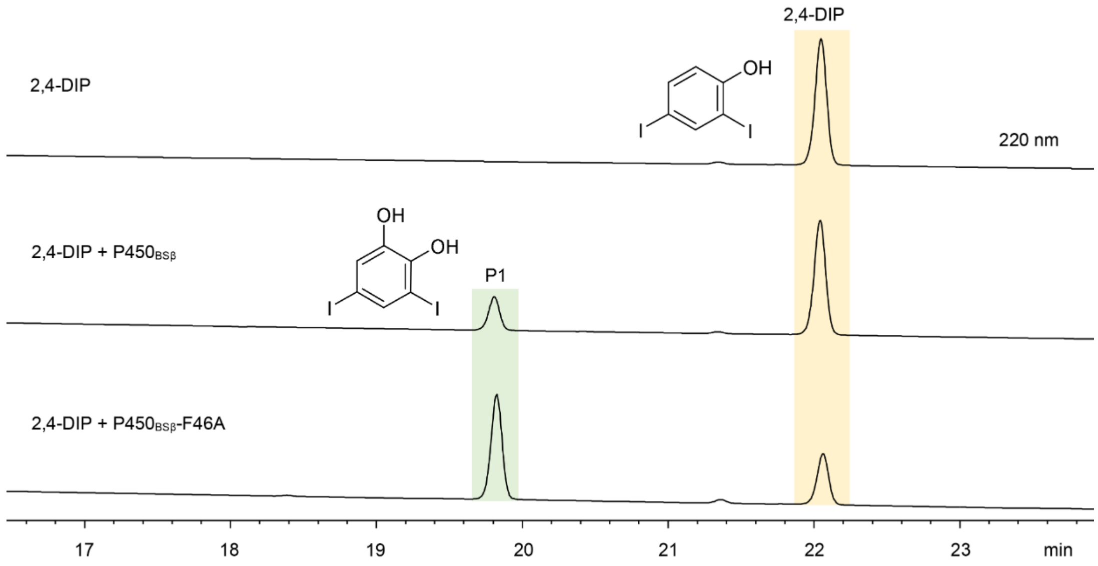


**Figure S37.** HPLC analysis of the reactions of 2,4-diiodophenol (2,4-DIP, **17**) catalyzed by P450$_{BS\beta}$ and P450$_{BS\beta}$-F46A. Reaction conditions: 1 μM P450, 500 μM substrate, and the AldO/glycerol system to generate $H_2O_2$ as cofactor at 30 °C for 6 h.

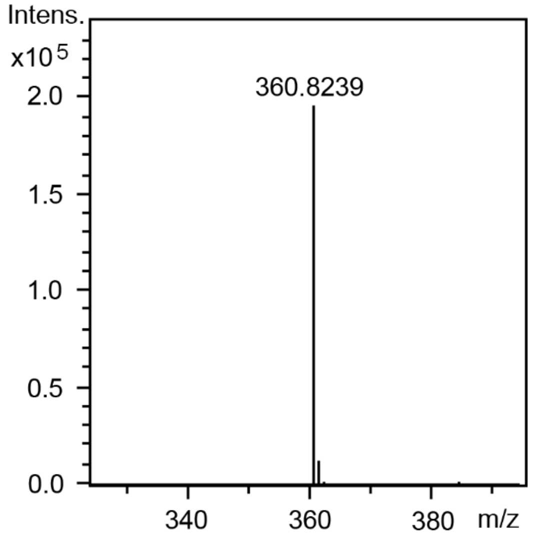


**Figure S38.** HR-ESI-MS of **17**-P1 ([M-H]$^-$: *calc.* 360.8228; *obs.* 360.8239).

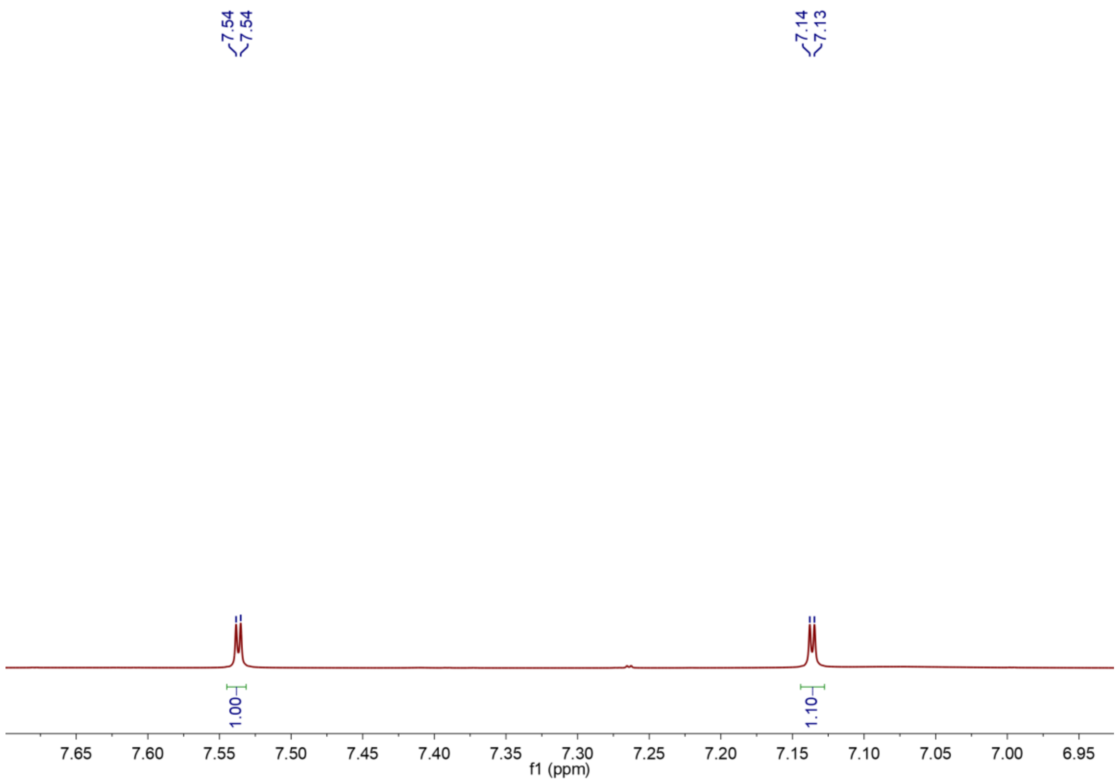


**Figure S39.** $^{1}H$ NMR spectrum of **17-**P1 in $CD_3CN$.

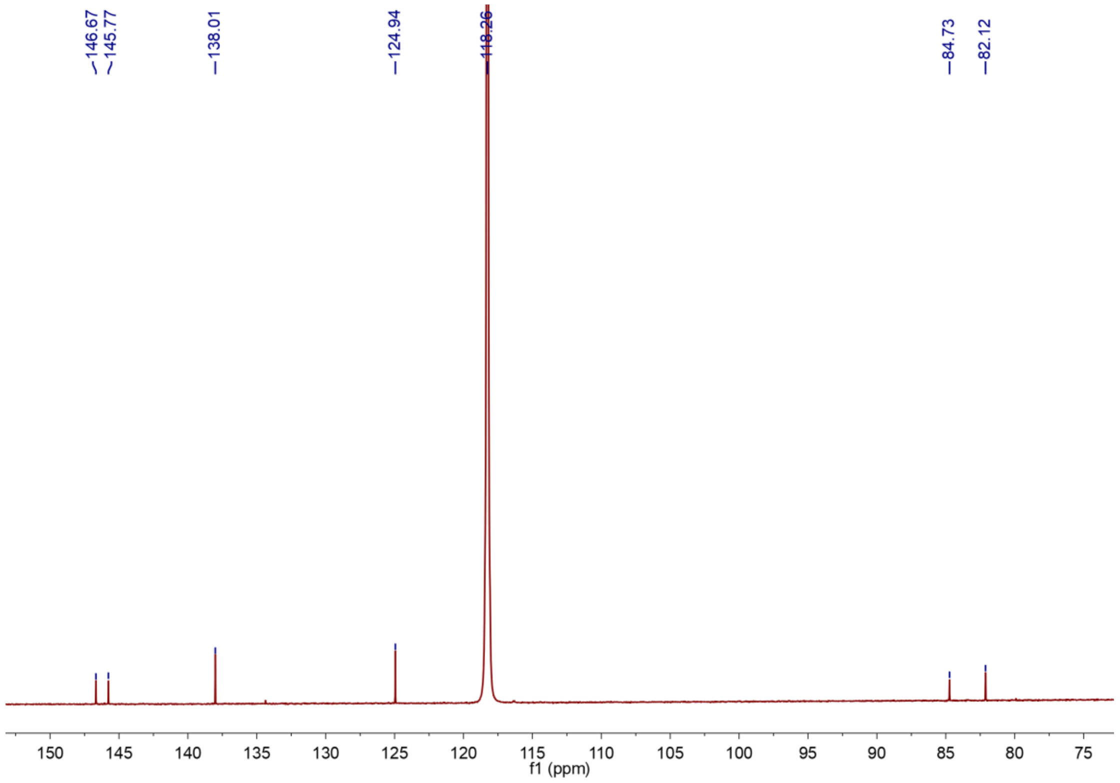


**Figure S40.** $^{13}C$-DEPTQ spectrum of **17**-P1 in $CD_3CN$.

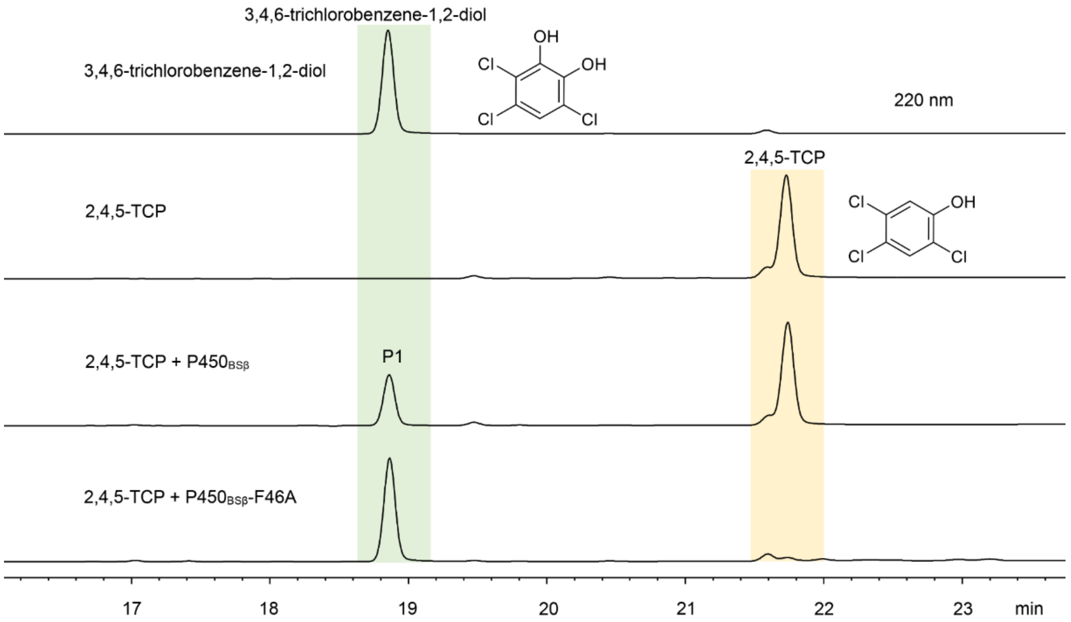


**Figure S41.** HPLC analysis of the reactions of 2,4,5-trichlorophenol (2,4,5-TCP, **18**) catalyzed by $P450_{BS\beta}$ and $P450_{BS\beta}$-F46A and 3,4,6-trichlorobenzene-1,2-diol standard. Reaction conditions: 1 μM P450, 500 μM substrate, and the AldO/glycerol system to generate $H_2O_2$ as cofactor at 30 °C for 6 h.

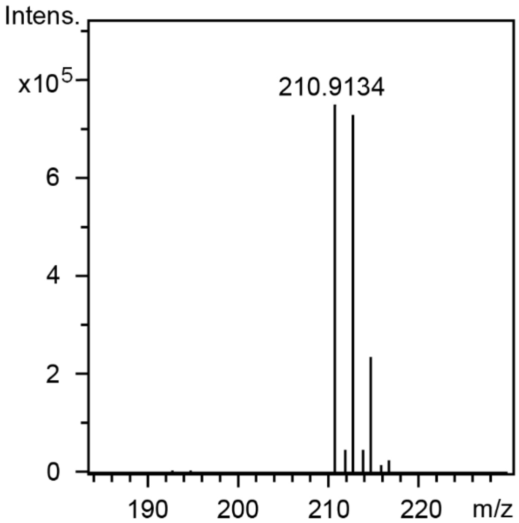


**Figure S42.** HR-ESI-MS of **18**-P1 ($[M-H]^-$: *calc.* 210.9126; *obs.* 210.9134).

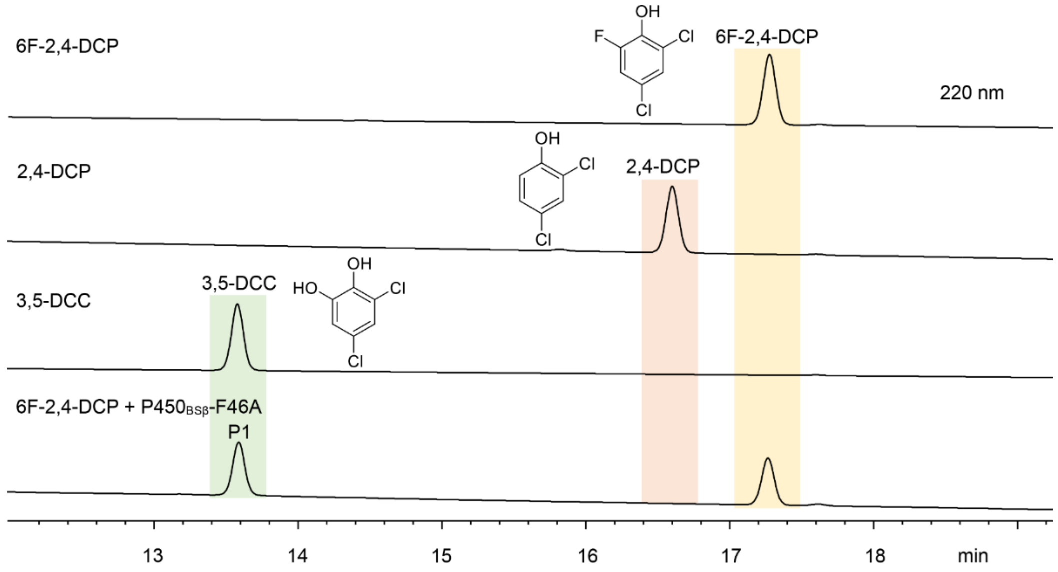


**Figure S43.** HPLC analysis of the reaction of 2,4-dichloro-6-fluorophenol (6F-2,4-DCP, **19**) catalyzed by $P450_{BS\beta}$-F46A and 2,4-dichlorophenol (2,4-DCP, **1**) and 3,5-dichlorocatechol (35-DCC, **2**) standards. Reaction conditions: 1 μM P450, 500 μM substrate, and the AldO/glycerol system to generate $H_2O_2$ as cofactor at 30 °C for 6 h.

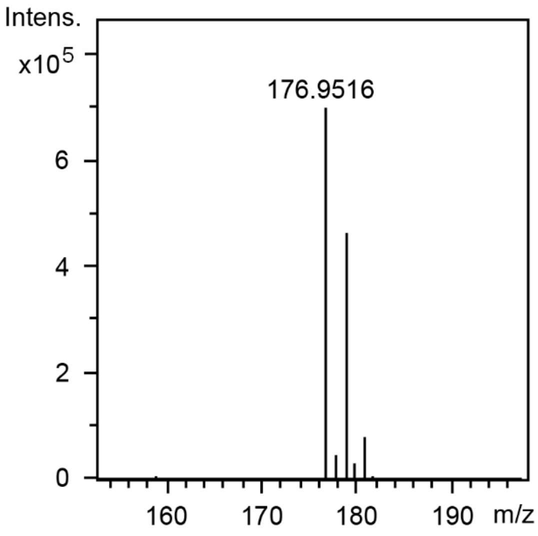


**Figure S44.** HR-ESI-MS of **19**-P1 ($[M-H]^-$: *calc.* 176.9516; *obs.* 176.9516).

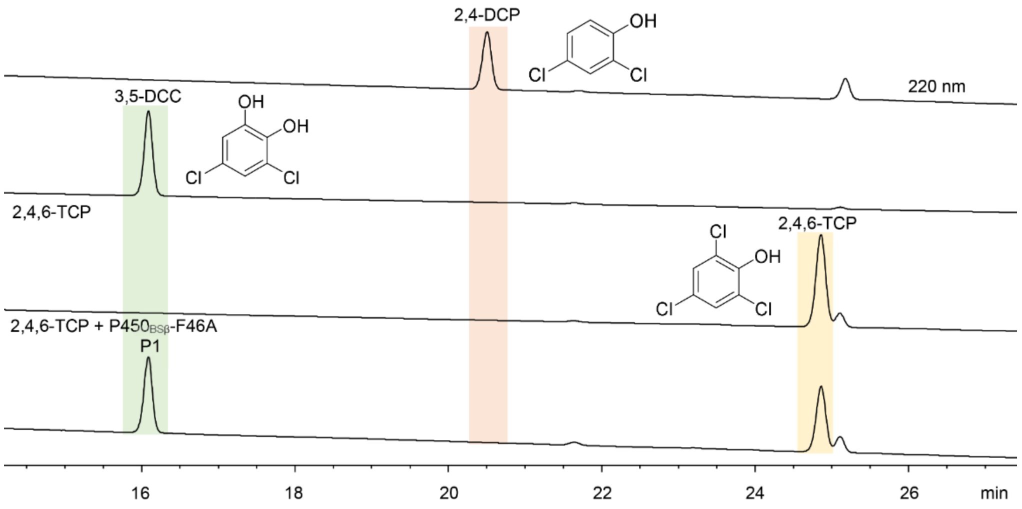


**Figure S45.** HPLC analysis of the reaction of 2,4,6-trichlorophenol (2,4,6-TCP, **20**) catalyzed by P450$_{BS\beta}$-F46A and 3,5-dichlorocatechol (3,5-DCC, **2**) standard. Reaction conditions: 1 μM P450, 500 μM substrate, and the AldO/glycerol system to generate $H_2O_2$ as cofactor at 30 °C for 6 h.

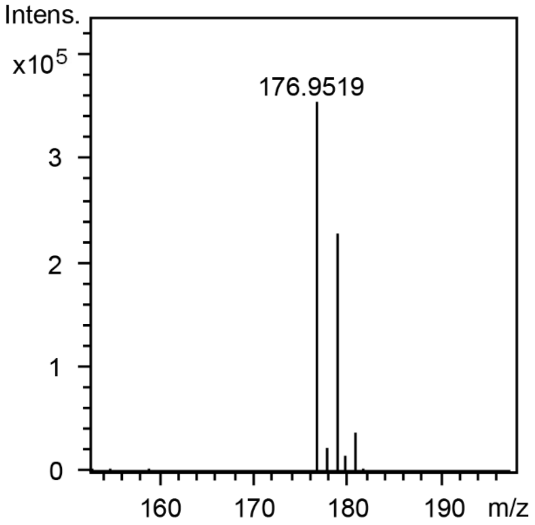


**Figure S46.** HR-ESI-MS of **20**-P1 ([M-H]$^-$: *calc.* 176.9516; *obs.* 176.9519).

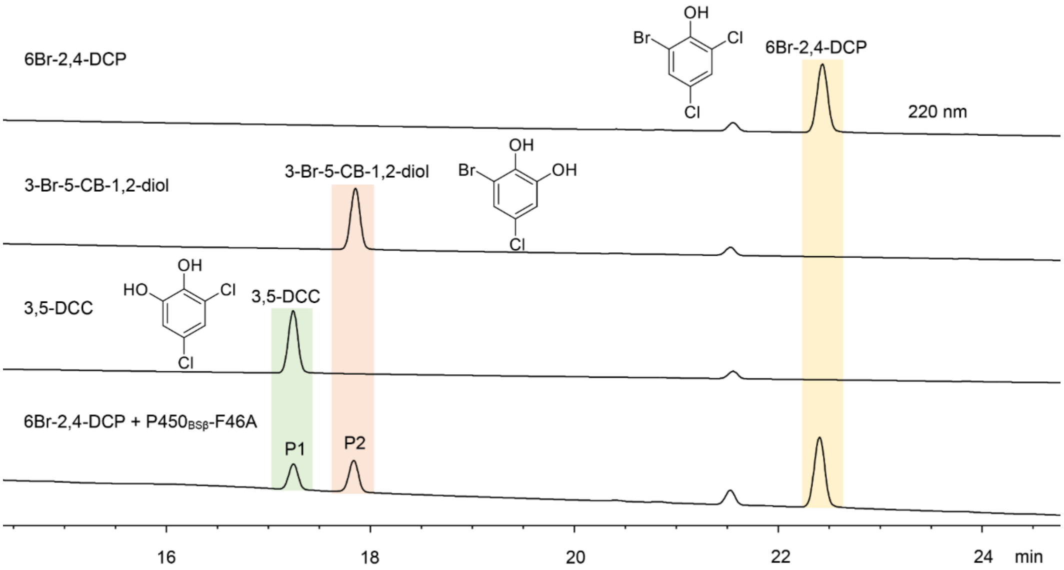


**Figure S47.** HPLC analysis of the reaction of 2,4-dichloro-6-bromophenol (6Br-2,4-DCP, **21**) catalyzed by $P450_{BS\beta}$-F46A and 3-bromo-5-chlorobenzene-1,2-diol (3-Br-5-CB-1,2-diol) and 3,5-dichlorocatechol (35-DCC, **2**) standards. Reaction conditions: 1 μM P450, 500 μM substrate, and the AldO/glycerol system to generate $H_2O_2$ as cofactor at 30 °C for 6 h.

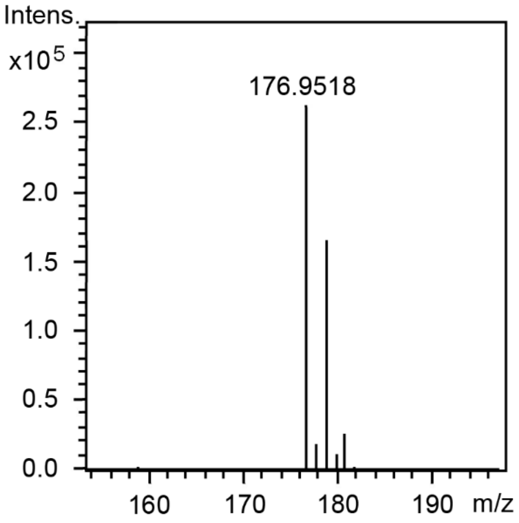


**Figure S48.** HR-ESI-MS of **21**-P1 ($[M-H]^-$: *calc.* 176.9516; *obs.* 176.9518).

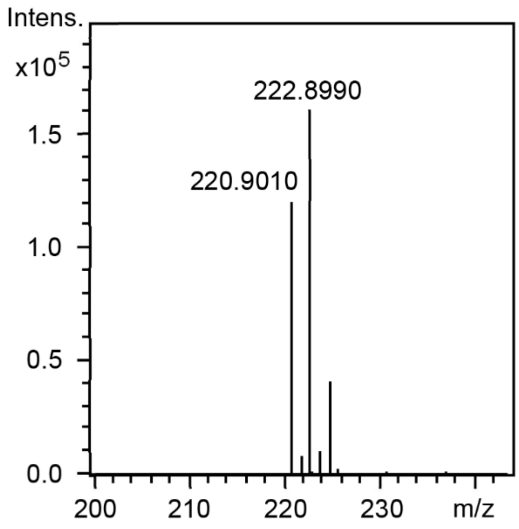


**Figure S49.** HR-ESI-MS of **21**-P2 ([M-H]$^-$: *calc.* 220.9010; *obs.* 220.9010).

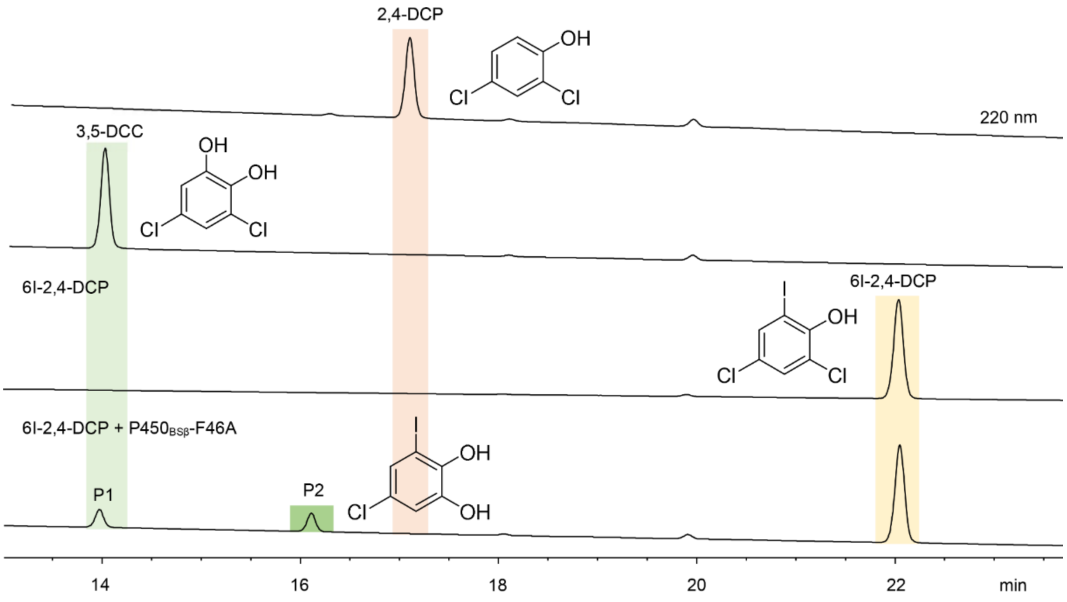


**Figure S50.** HPLC analysis of the reaction of 2,4-dichloro-6-iodophenol (6I-2,4-DCP, **22**) catalyzed by P450$_{BS\beta}$-F46A. Reaction conditions: 1 µM P450, 500 µM substrate, and the AldO/glycerol system to generate $H_2O_2$ as cofactor at 30 °C for 6 h.

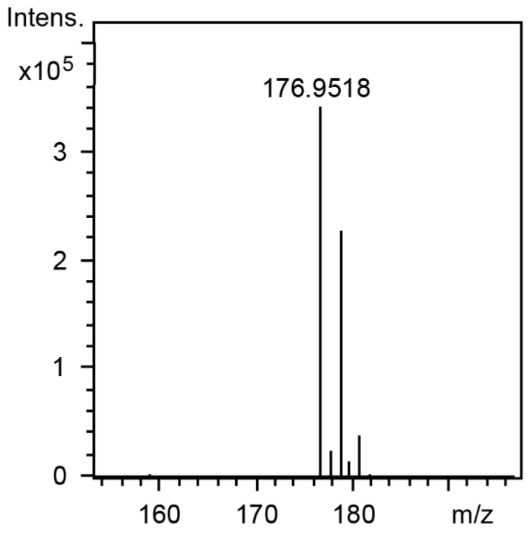


**Figure S51.** HR-ESI-MS of **22**-P1 ([M-H]$^-$: *calc.* 176.9516; *obs.* 176.9518).

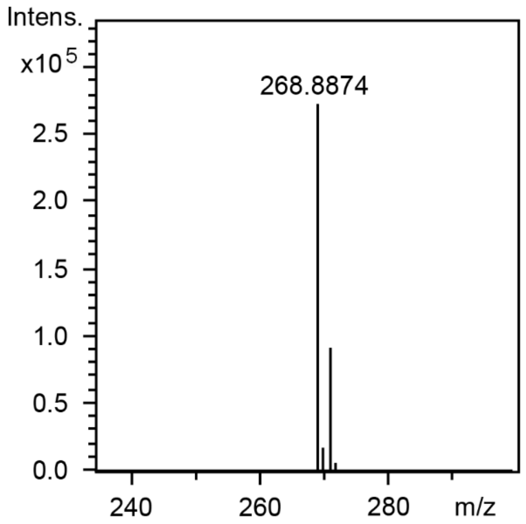


**Figure S52.** HR-ESI-MS of **22**-P2 ([M-H]$^-$: *calc.* 268.8872; *obs.* 268.8874).

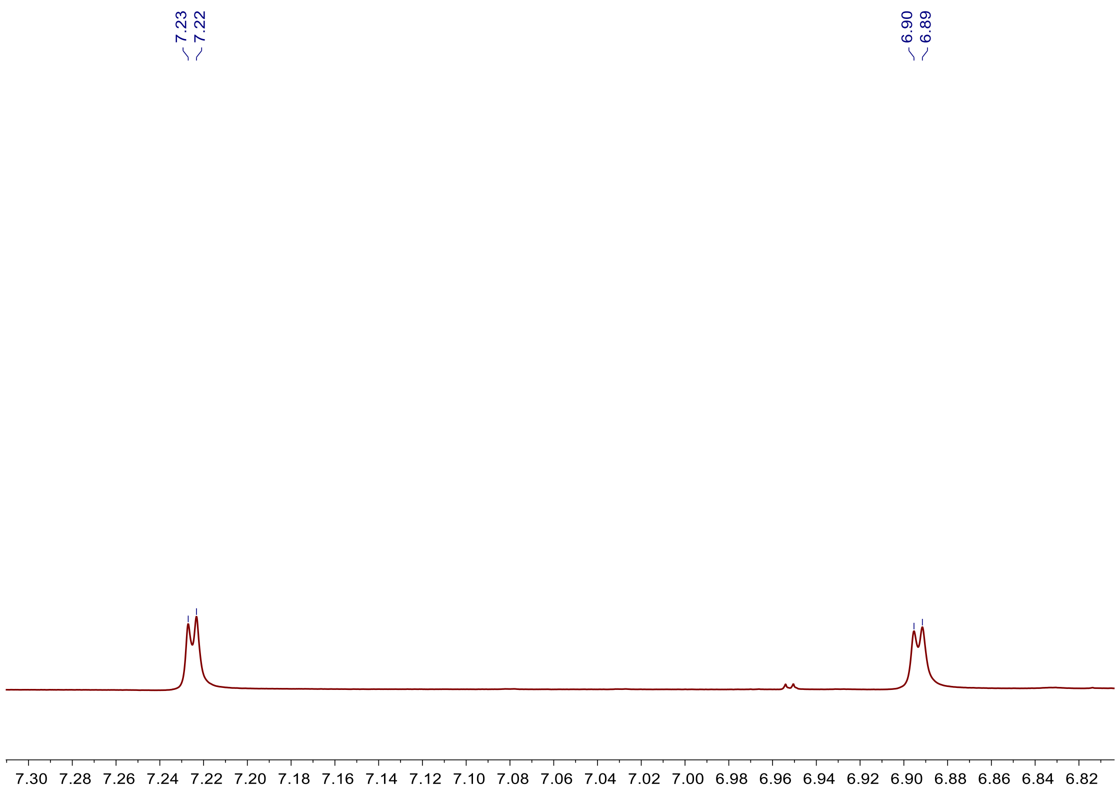


**Figure S53.** $^{1}H$ NMR spectrum of **22**-P2 in $CD_3CN$.

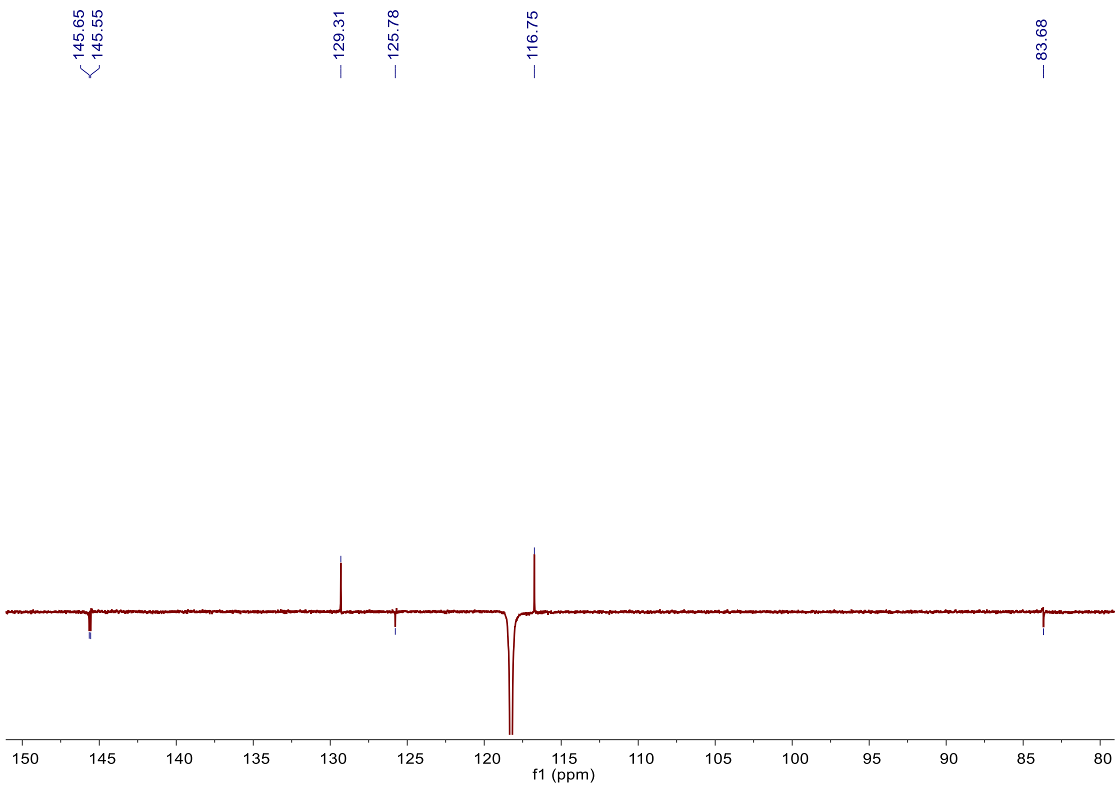


**Figure S54.** $^{13}C$-DEPTQ spectrum of **22**-P2 in $CD_3CN$.

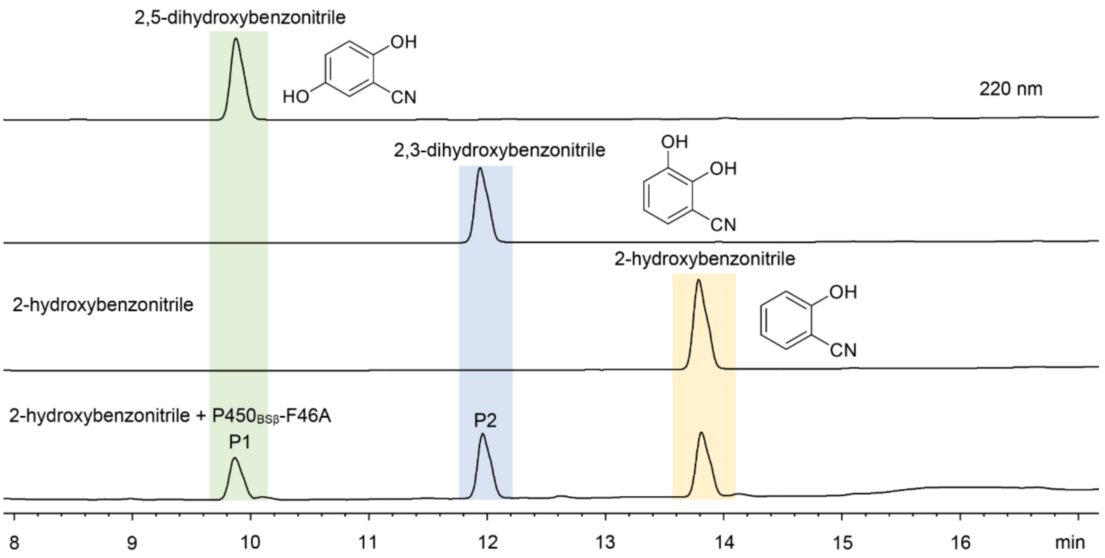


**Figure S55.** HPLC analysis of the reaction of 2-hydroxybenzonitrile (**23**) catalyzed by P450$_{BS\beta}$-F46A, 2,5-dihydroxybenzonitrile and 2,3-dihydroxybenzonitrile standards. Reaction conditions: 1 μM P450, 500 μM substrate, and the AldO/glycerol system to generate $H_2O_2$ as cofactor at 30 °C for 6 h.

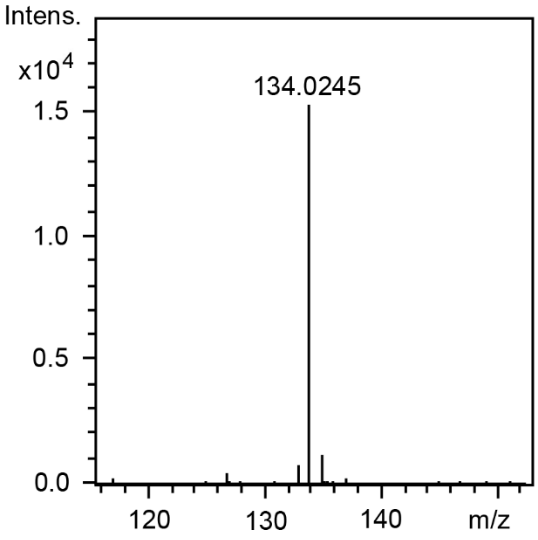


**Figure S56.** HR-ESI-MS of **23**-P1 ([M-H]$^-$: *calc.* 134.0248; *obs.* 134.0245).

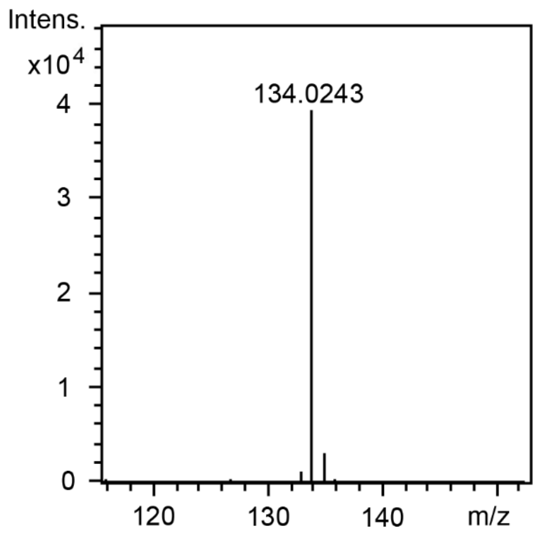


**Figure S57.** HR-ESI-MS of **23**-P2 ([M-H]$^-$: *calc.* 134.0248; *obs.* 134.0243).

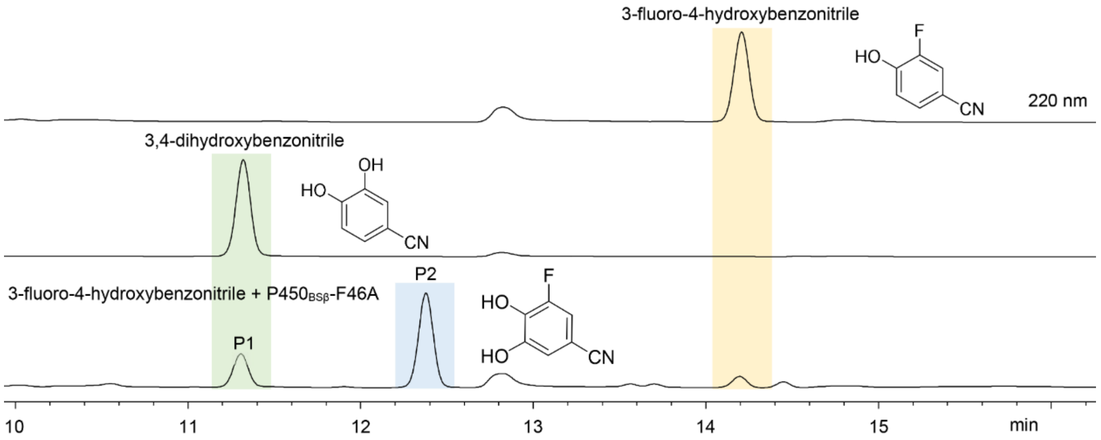


**Figure S58.** HPLC analysis of the reaction of 3-fluoro-4-hydroxybenzonitrile (**24**) catalyzed by P450$_{BS\beta}$-F46A and 3,4-dihydrobenzonitrile standard. Reaction conditions: 1 μM P450, 500 μM substrate, and the AldO/glycerol system to generate $H_2O_2$ as cofactor at 30 °C for 6 h.

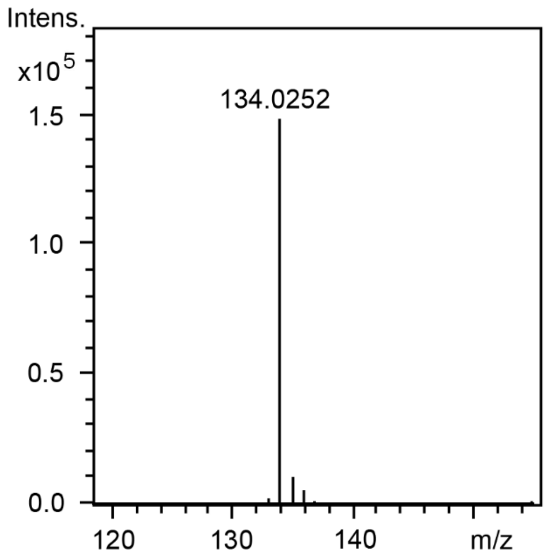


**Figure S59.** HR-ESI-MS of **24**-P1 ([M-H]$^-$: *calc.* 134.0248; *obs.* 176.9516).

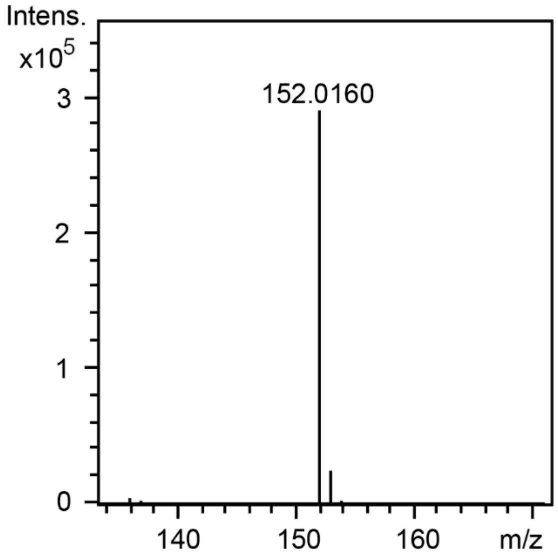


**Figure S60.** HR-ESI-MS of **24**-P2 ([M-H]$^-$: *calc.* 152.0160; *obs.* 176.9516).

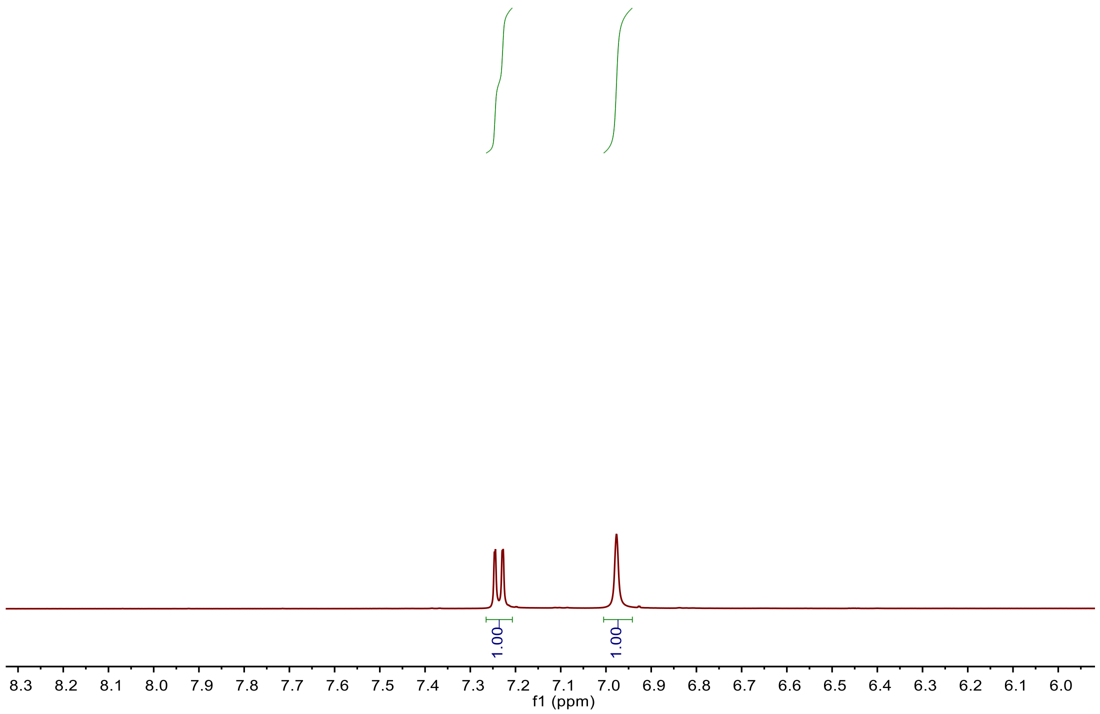


**Figure S61.** $^{1}$H NMR spectrum of **24**-P2 in DMSO-*d*6.

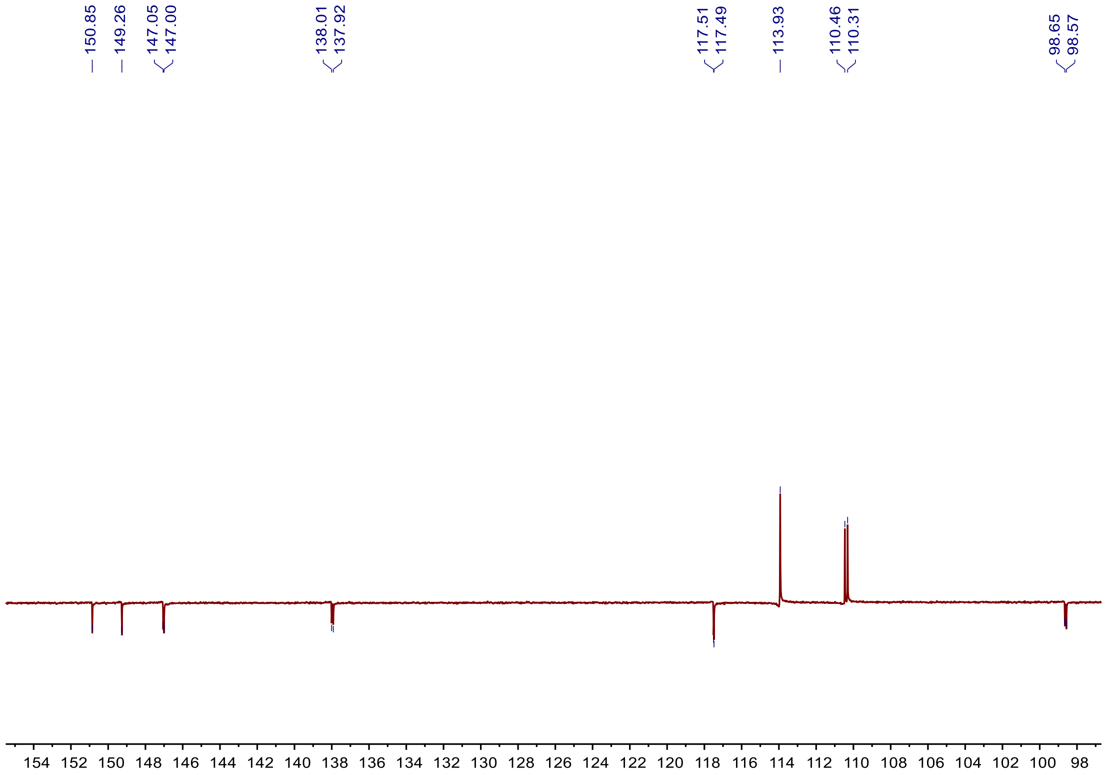


**Figure S62.** $^{13}$C-DEPTQ spectrum of **24**-P2 in DMSO-$d_6$.

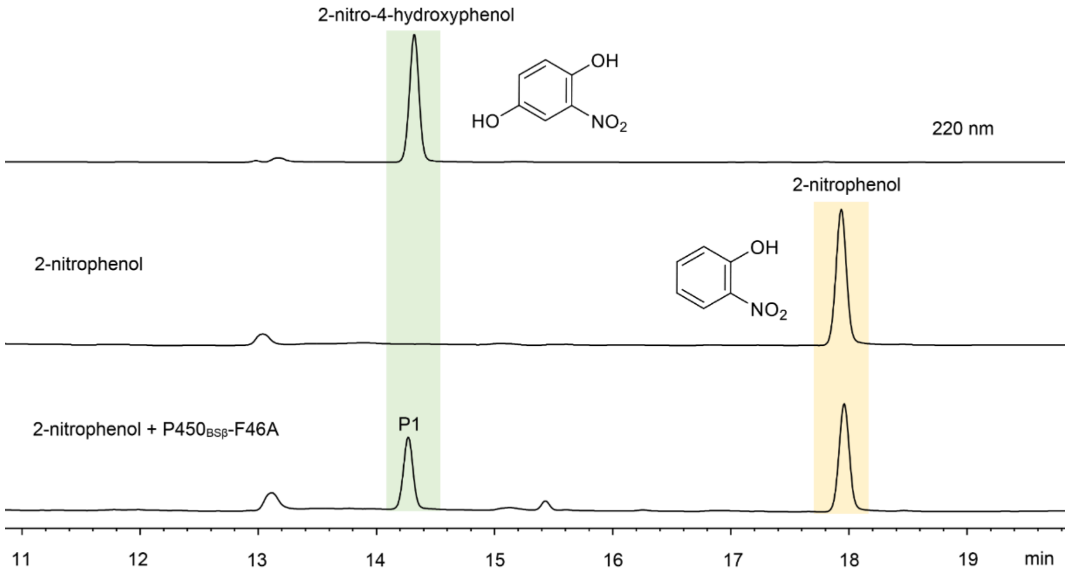


**Figure S63.** HPLC analysis of the reaction of 2-nitrophenol (**25**) catalyzed by P450$_{BSβ}$-F46A and 2-nitro-4-hydroxyphenol standard. Reaction conditions: 1 μM P450, 500 μM substrate, and the AldO/glycerol system to generate $H_2O_2$ as cofactor at 30 °C for 6 h.

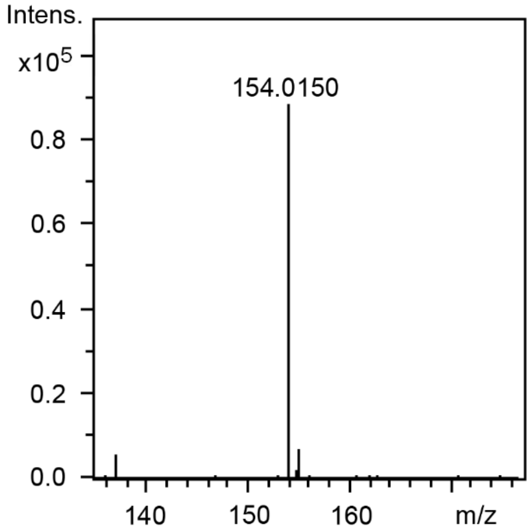


**Figure S64.** HR-ESI-MS of **25-**P1 ([M-H]$^-$: *calc.* 154.0146; *obs.* 154.0150).

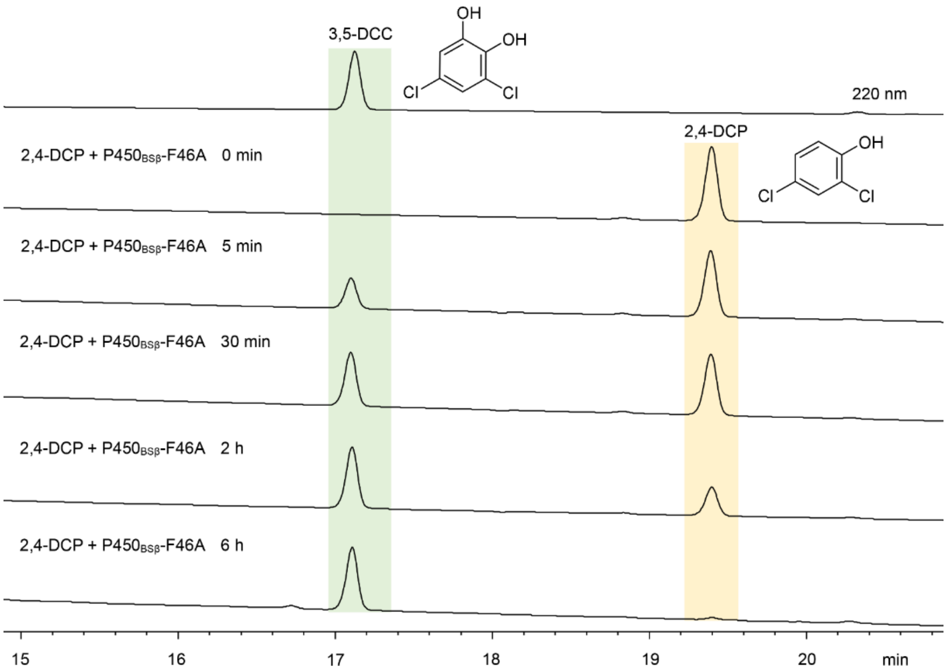


**Figure S65.** HPLC analysis of the time-course experiments of the reactions of 2,4-DCP (**1**) catalyzed by P450$_{BS\beta}$-F46A. Reaction conditions: 1 μM P450$_{BS\beta}$-F46A supported by the AldO/glycerol $H_2O_2$ generation system at 30°C.

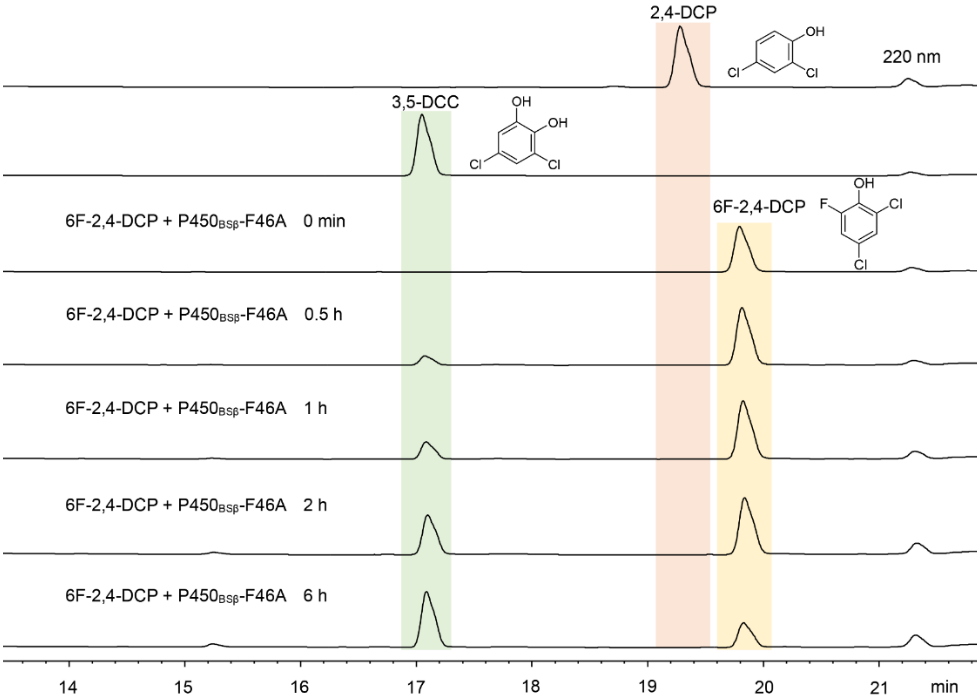


**Figure S66.** HPLC analysis of the time-course experiments of the reactions of 6F-2,4-DCP (**19**) catalyzed by P450$_{BS\beta}$-F46A. Reaction conditions: 1 μM P450$_{BS\beta}$-F46A supported by the AldO/glycerol $H_2O_2$ generation system at 30°C.

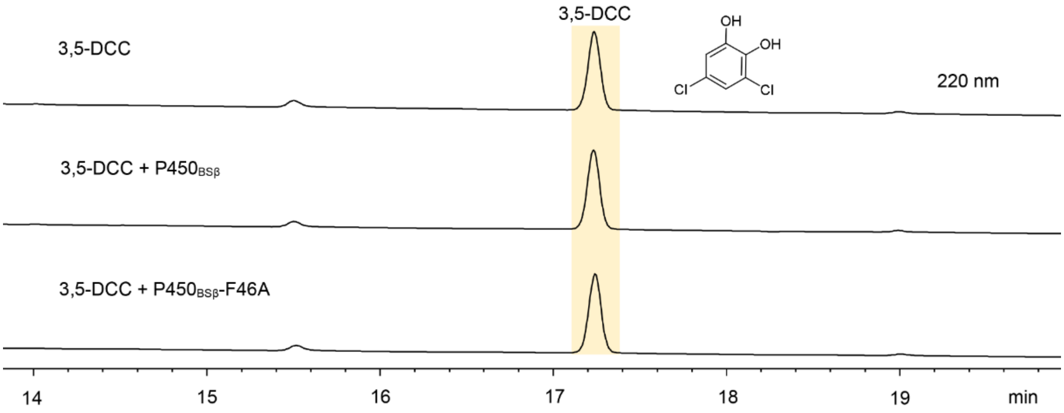


**Figure S67.** HPLC analysis of the reactions of 3,5-dichlorocatechol (3,5-DCC, **2**) catalyzed by P450$_{BSβ}$ and P450$_{BSβ}$-F46A. Reaction conditions: 1 µM P450, 500 µM substrate, and the AldO/glycerol system to generate $H_2O_2$ as cofactor at 30 °C for 6 h.

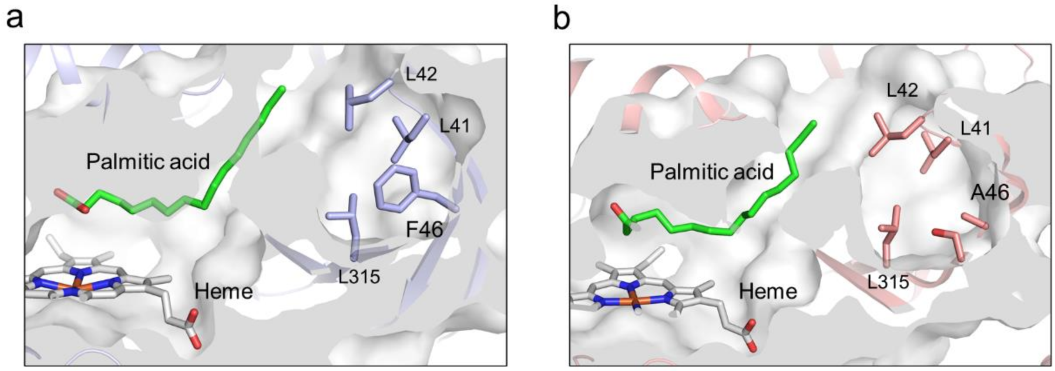


**Figure S68.** Comparative analysis of crystal structure of wild-type P450$_{BSβ}$ (**a**) and P450$_{BSβ}$-F46A (**b**) in complex with palmitic acid.

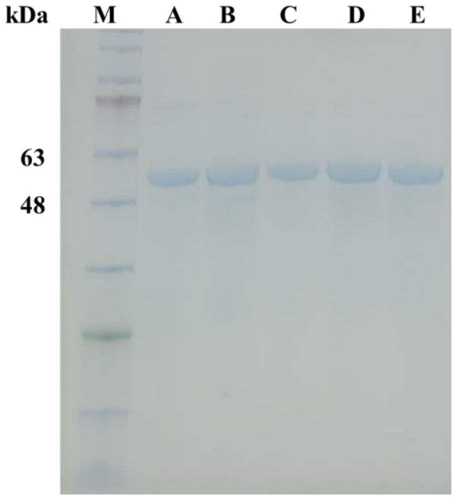


**Figure S69.** SDS-PAGE analysis of the purified $His_6$-tagged $P450_{BS\beta}$-F46A (lane A), $P450_{BS\beta}$-F46A-R242A (lane B), $P450_{BS\beta}$-F46A-R242K (lane C), $P450_{BS\beta}$-F46A-R242E (lane D), $P450_{BS\beta}$-F46A-R242S (lane E) and protein marker (M).

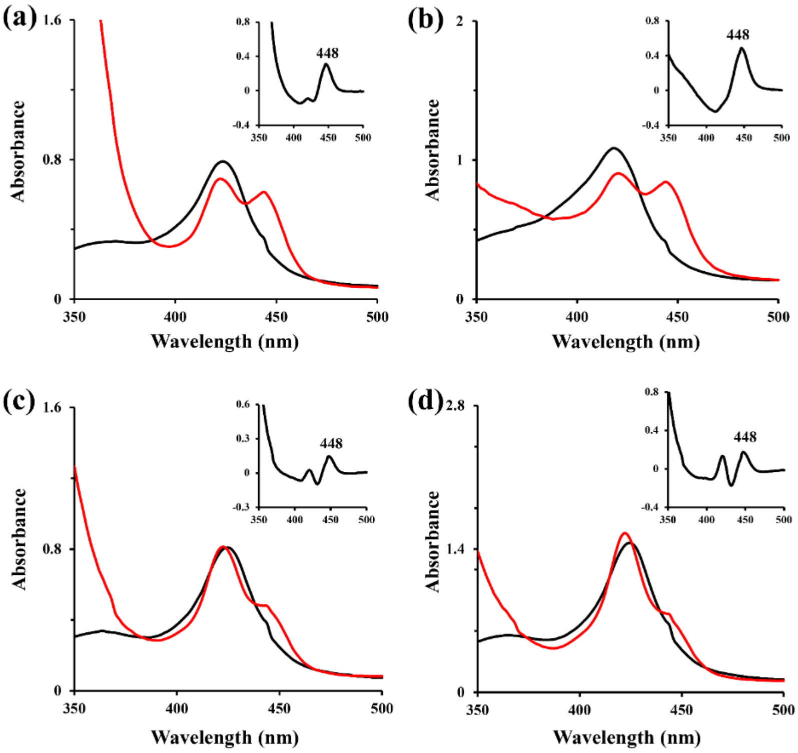


**Figure S70.** UV-visible spectra of $P450_{BS\beta}$-F46A-R242A (**a**), $P450_{BS\beta}$-F46A-R242K (**b**), $P450_{BS\beta}$-F46A-R242E (**c**), and $P450_{BS\beta}$-F46A-R242S (**d**). (Black lines show the spectra for the oxidized ferric form of CYPs and red lines show the spectra for the $Na_2S_2O_4$-reduced ferrous-CO complex of CYPs; Insets exhibit the CO-bound reduced difference spectra of P450 enzymes).

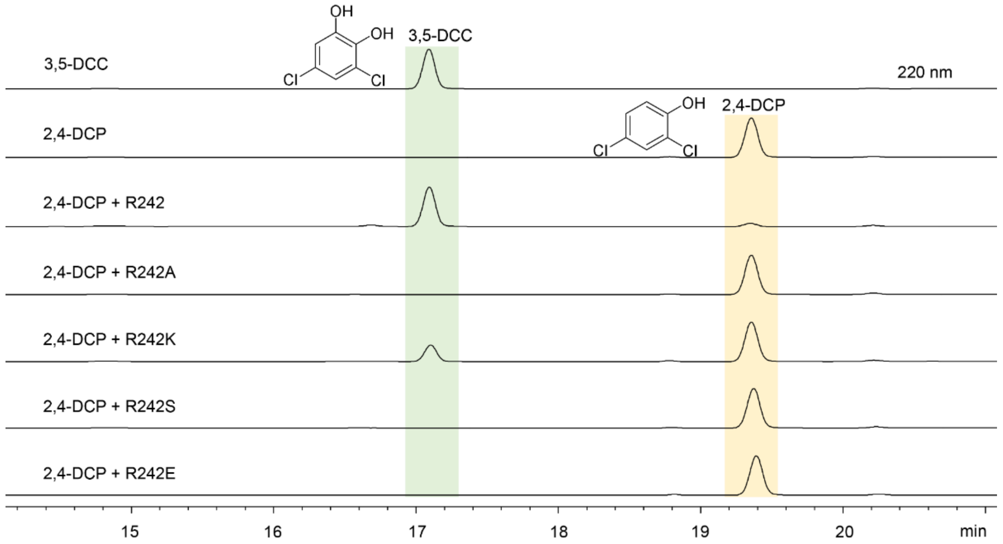


**Figure S71.** HPLC analysis of the reactions of 2,4-dichlorophenol (2,4-DCP, **1**) catalyzed by $P450_{BS\beta}$, $P450_{BS\beta}$-F46A-R242A, $P450_{BS\beta}$-F46A-R242K, $P450_{BS\beta}$-F46A-R242S and $P450_{BS\beta}$-F46A-R242E, and 3,5-dichlorocatechol (3,5-DCC, **2**) standard. Reaction conditions: 1 μM P450, 500 μM substrate, and the AldO/glycerol system to generate $H_2O_2$ as cofactor at 30 °C for 6 h.

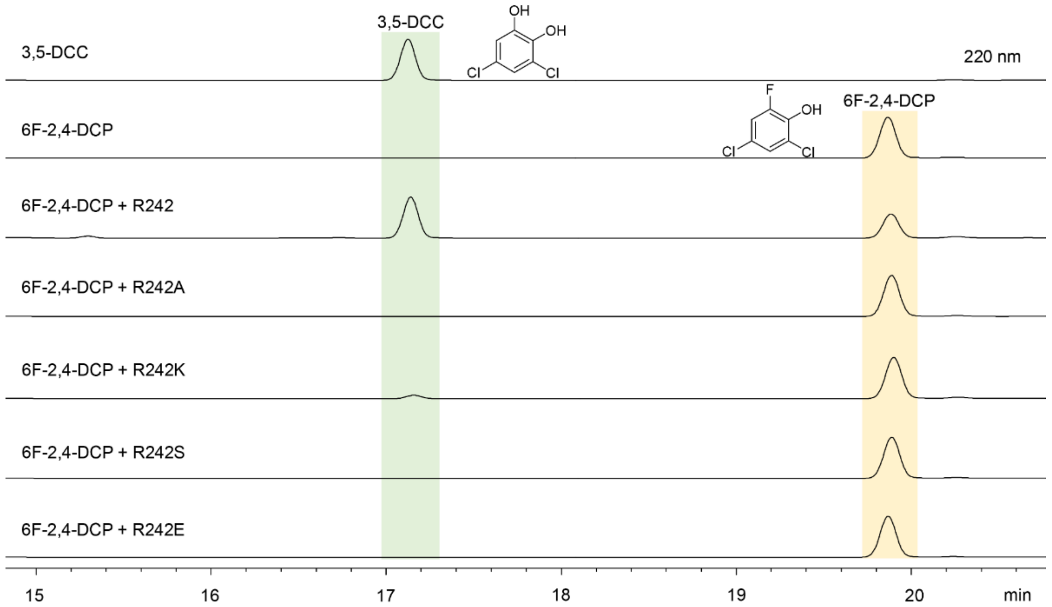


**Figure S72.** HPLC analysis of the reactions of 2,4-dichloro-6-fluorophenol (6F-2,4-DCP, **19**) catalyzed by $P450_{BS\beta}$-F46A, $P450_{BS\beta}$-F46A-R242A, $P450_{BS\beta}$-F46A-R242S, $P450_{BS\beta}$-F46A-R242K and $P450_{BS\beta}$-F46A-R242E, and 3,5-dichlorocatechol (3,5-DCC, **2**) standard. Reaction conditions: 1 μM P450, 500 μM substrate, and the AldO/glycerol system to generate $H_2O_2$ as cofactor at 30 °C for 6 h.

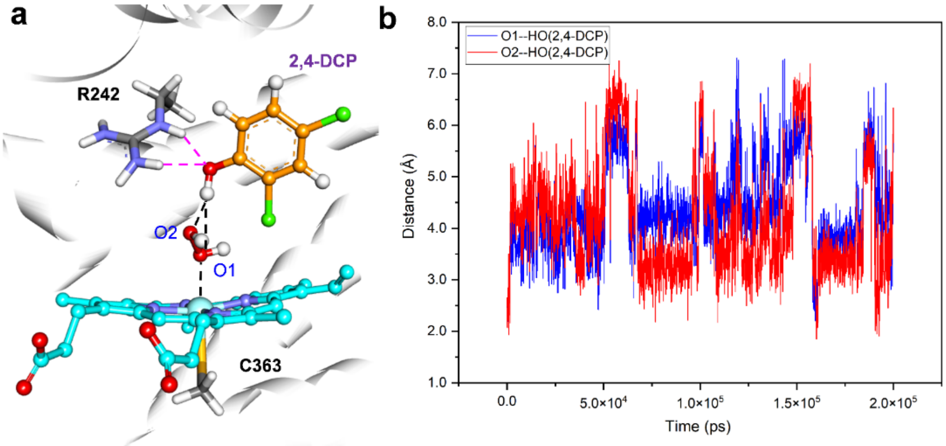


**Figure S73.** The fluctuation of the distances between O1/O2 atom of $H_2O_2$ and the hydroxyl group of substrate 2,4-DCP (**1**) during the 200 ns MD simulation.

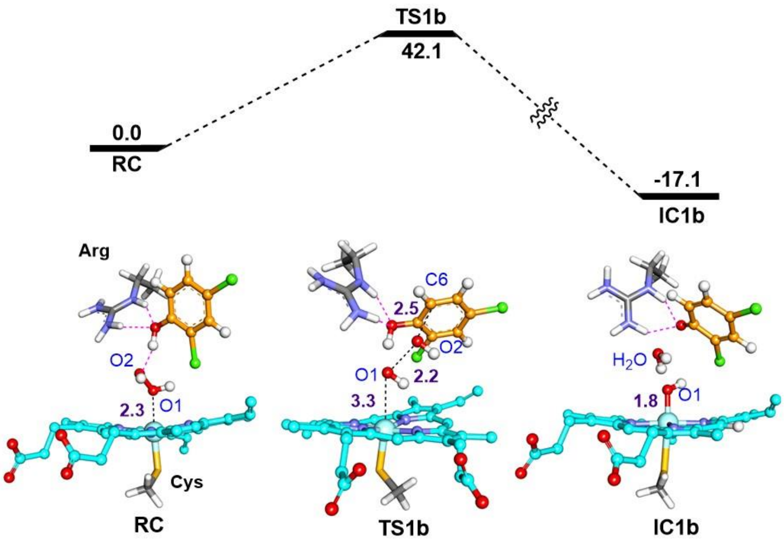


**Figure S74.** QM/MM calculated O-O homolysis and $OH^{\bullet}$ attach to **1** by $P450_{BS\beta}$-F46A, along with the QM/MM optimized structures of key species involved in the reactions. The energies are given in kcal/mol while the key distances are given in Å.

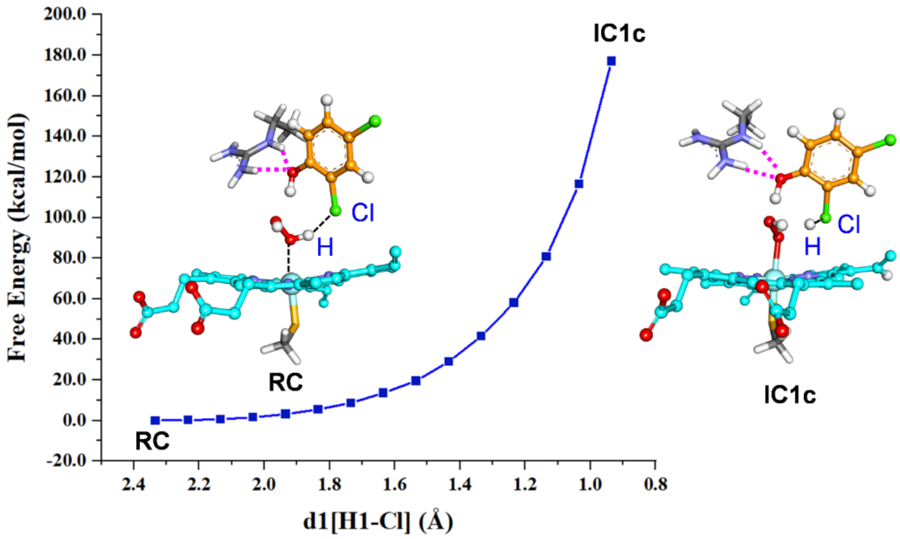


**Figure S75.** QM/MM calculated the proximal proton transfer from $H_2O_2$ to the substrate's chlorine ion in $P450_{BS\beta}$-F46A, along with the QM/MM optimized structures of key species involved in the reactions. The energies are given in kcal/mol.

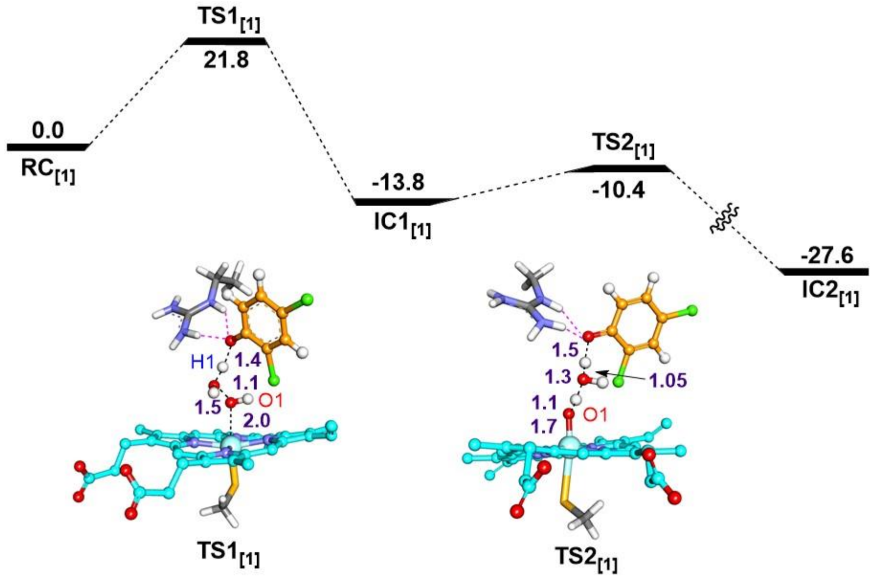


**Figure S76.** The calculated mechanism (with energies in kcal/mol) for the formation of Cpd I by $P450_{BS\beta}$-F46A in the presence of the non-carboxylic substrate 2,4-DCP (**1**). The non-carboxylic substrate 2,4-DCP phenolic hydroxyl group interacting with the guanidyl group of R242 is located above the heme. The key distances are given in angstrom (Å).

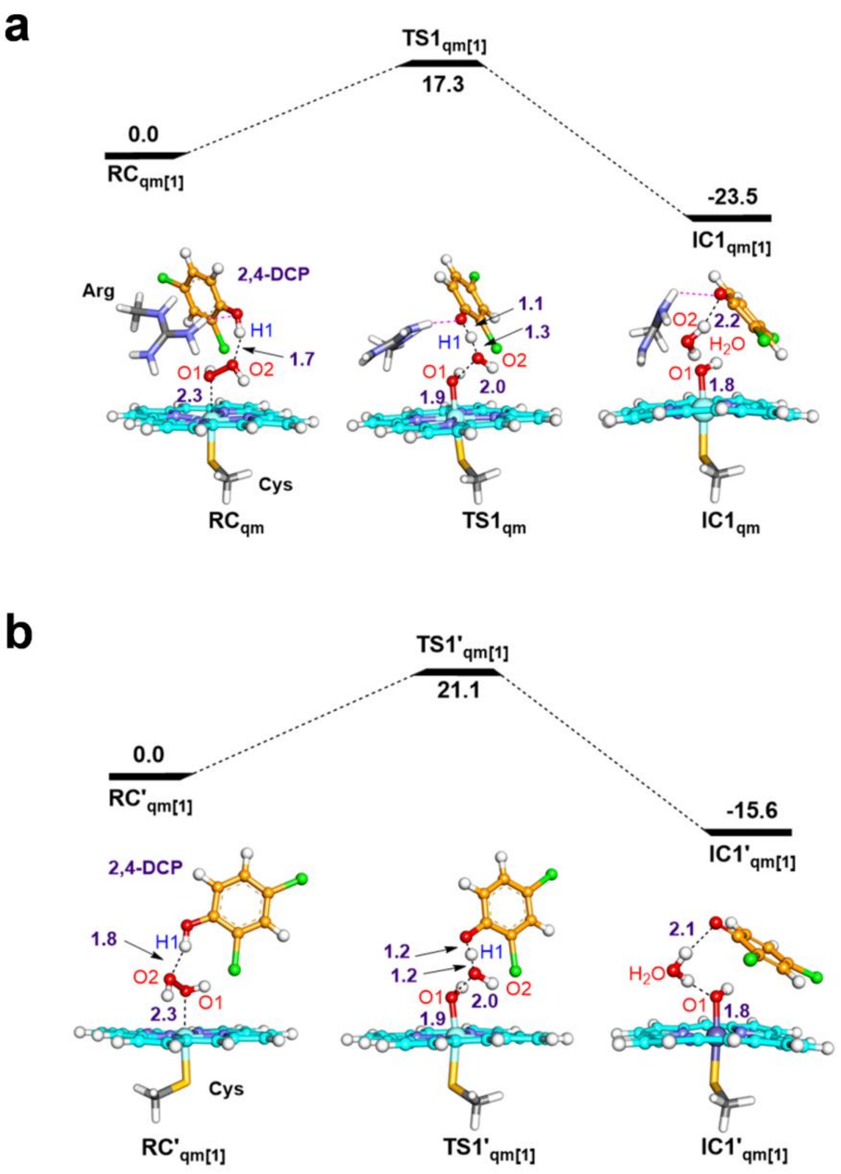


**Figure S77.** QM calculated O-O homolysis route (with energies in kcal/mol) toward Cpd I by P450$_{BS\beta}$-F46A in the presence of the non-carboxylic substrate 2,4-DCP (**1**) and conserved R242. (**a**), with the conserved R242 and (**b**), without the conserved R242, along with the QM optimized structures of key species involved in the reactions. The energies are given in kcal/mol while the key distances are given in Å.

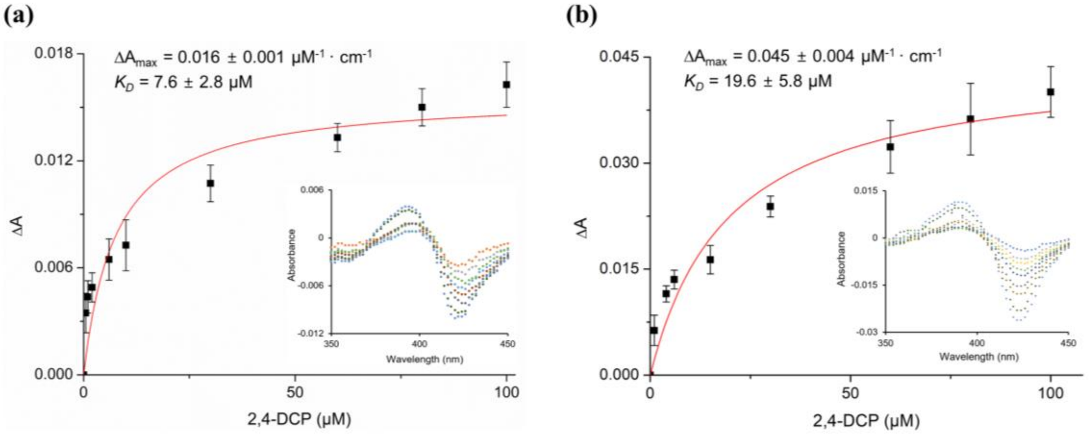


**Figure S78.** Substrate binding curves of P450$_{BS\beta}$-F46A (**a**) and P450$_{BS\beta}$-F46A-R242A (**b**) with 2,4-DCP (**1**). The insets show the Type I difference spectra. All experiments were performed in triplicate.

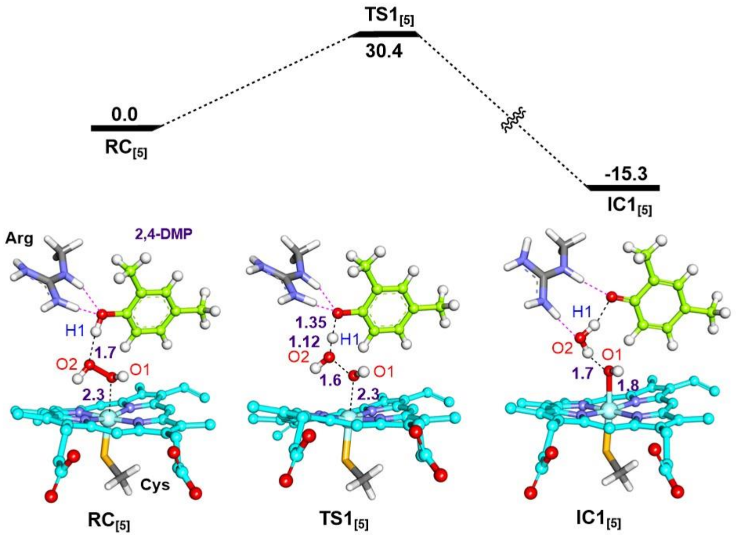


**Figure S79.** QM/MM calculated mechanism for the formation of Cpd I by P450$_{BS\beta}$-F46A in the presence of the non-carboxylic substrate 2,4-dimethylphenol (2,4-DMP, **5**), along with the QM/MM optimized structures of key species involved in the reactions. The energies are given in kcal/mol while the key distances are given in Å.

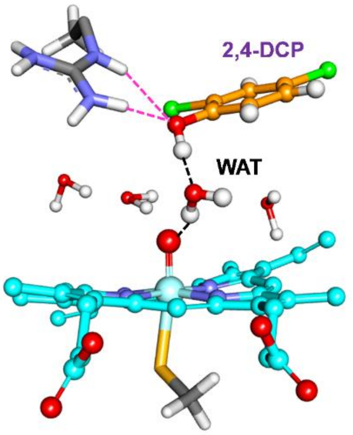


| | WAT No. | H-bond (%) |
|---|---|---|
| 2,4DCP: O atom | 18292 | 21.65 |
| | 18550 | 17.55 |
| | 4135 | 12.90 |
| | 13932 | 8.30 |
| | 923 | 4.95 |
| | 3657 | 4.45 |
| | 19121 | 3.30 |
| | 4177 | 3.25 |
| | 5342 | 2.85 |
| | 13003 | 2.50 |

**Figure S80.** The numbering and statistical proportion of water molecules forming hydrogen bonds with substrate and Cpd I. The solvent water molecules near Cpd I could easily replace the water molecule generated from $H_2O_2$ and make H-bond interactions with O1 of Cpd I and substrate 2,4-DCP (**1**) during the MD simulation.

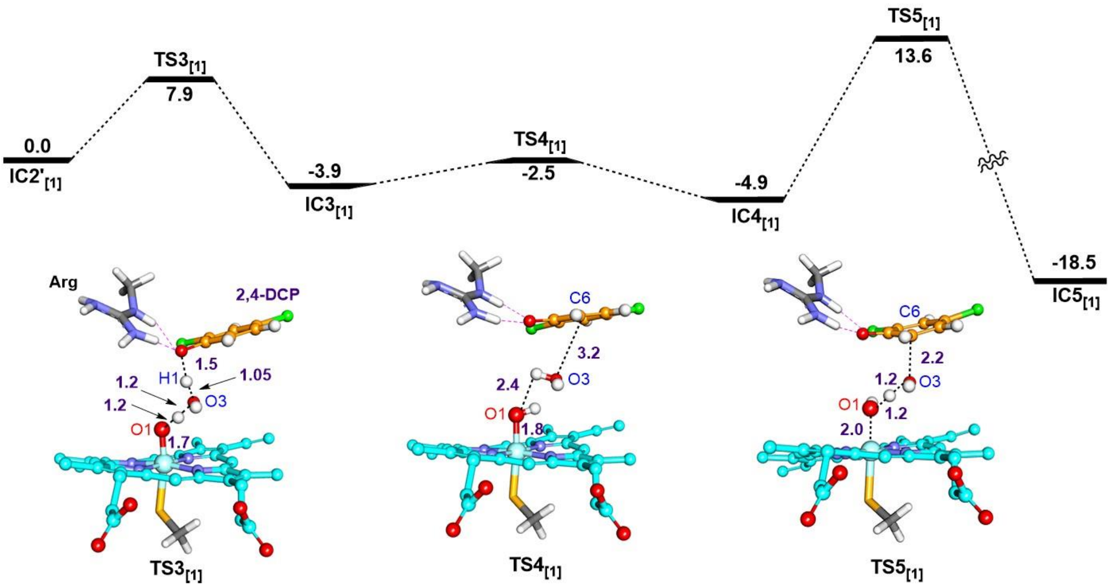


**Figure S81.** QM/MM calculated mechanism for the formation of ene-ketone intermediate starting from Cpd I of $P450_{BS\beta}$-F46A, along with the QM/MM optimized structures of TSs species involved in the reactions. The energies are given in kcal/mol while the key distances are given in Å.

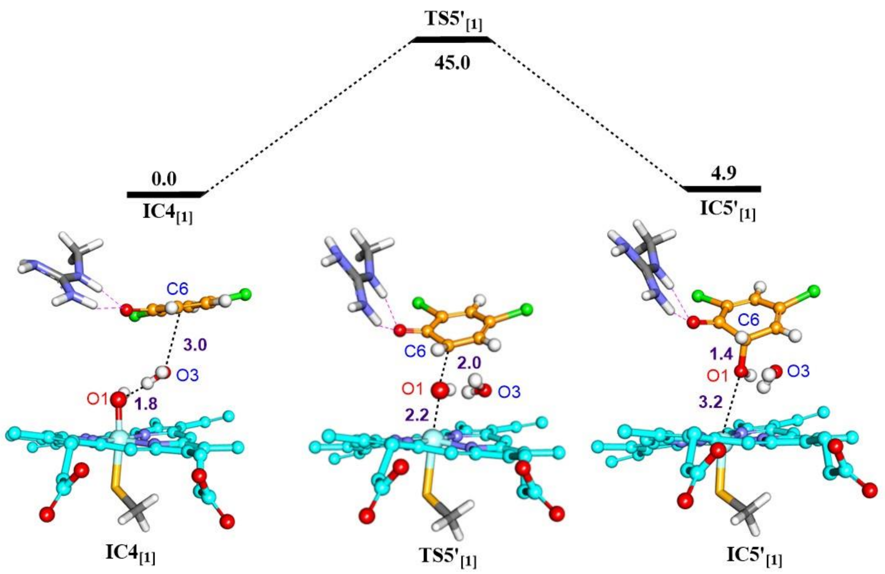


**Figure S82.** QM/MM calculated mechanism for the OH radical of Cpd II directly attack the substrate in $P450_{BS\beta}$-F46A. The energies are given in kcal/mol while the key distances are given in Å.

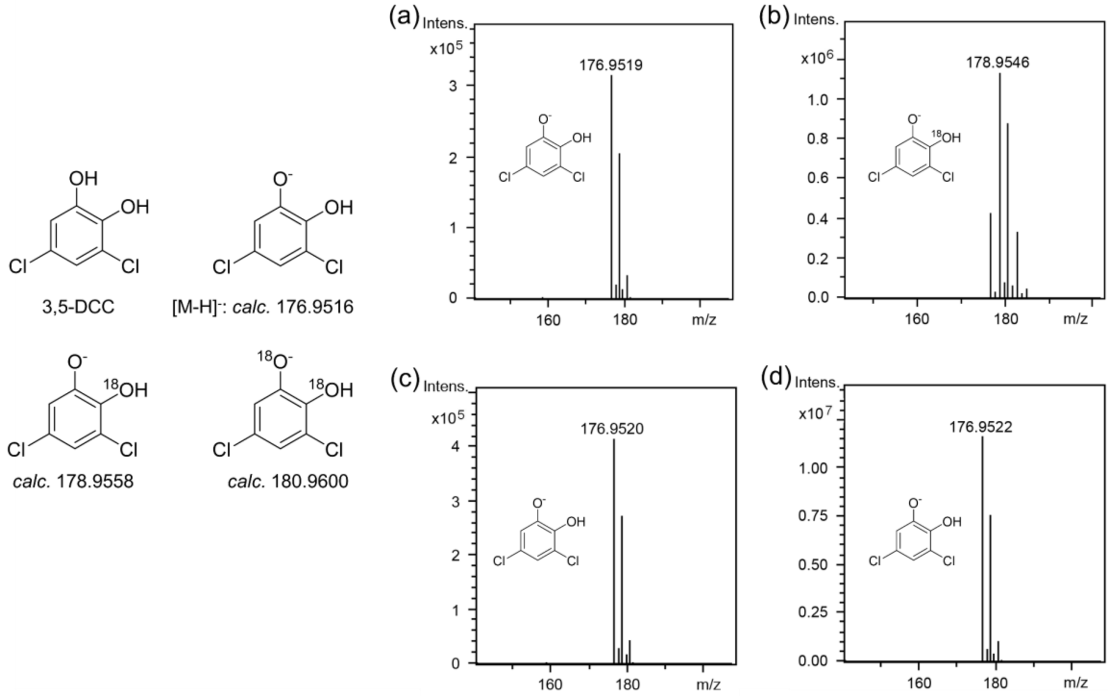


**Figure S83.** LC-MS analysis of 3,5-DCC (**2**) standard in $H_2O_2$ (**a**), $H_2{}^{18}O$ (**b**), $H_2{}^{18}O_2$ (**c**), and $H_2O$ (**d**). These results suggest that one oxygen atom in the standard product **2** underwent an exchange with the $^{18}O$ label from water (**b**), indicative of a potential scrambling event with $H_2{}^{18}O$ solvent.

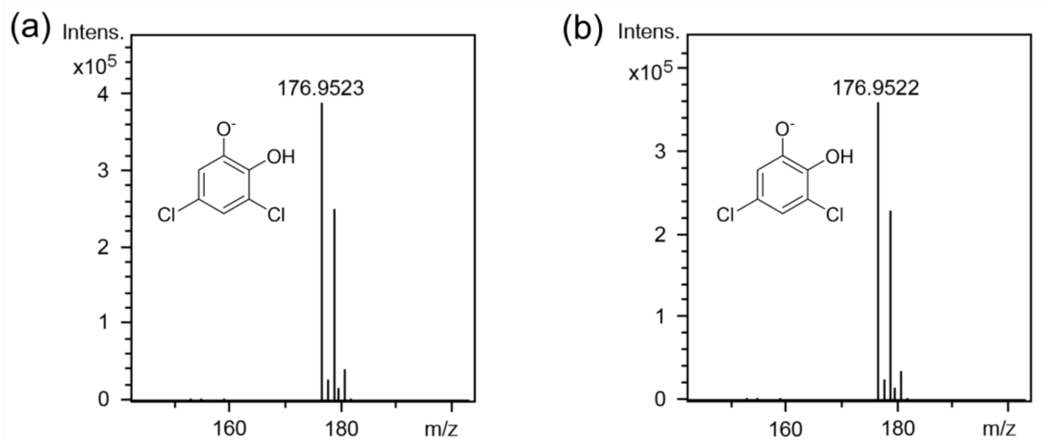


**Figure S84.** LC-MS analysis of **2** production from **1** by P450$_{BS\beta}$-F46A when supported by $H_2O_2$ (**a**) or $H_2{}^{18}O_2$ (**b**). Reaction conditions: 1 μM P450$_{BS\beta}$-F46A, 500 μM **1**, 500 μM $H_2O_2$ or $H_2{}^{18}O_2$ as cofactor at 30 °C for 6 h. These $H_2{}^{18}O_2$/$H_2O_2$ tracing experiments showed that the hydroxyl oxygen atom of **2** production from **1** by P450$_{BS\beta}$-F46A did not originate from hydrogen peroxide.

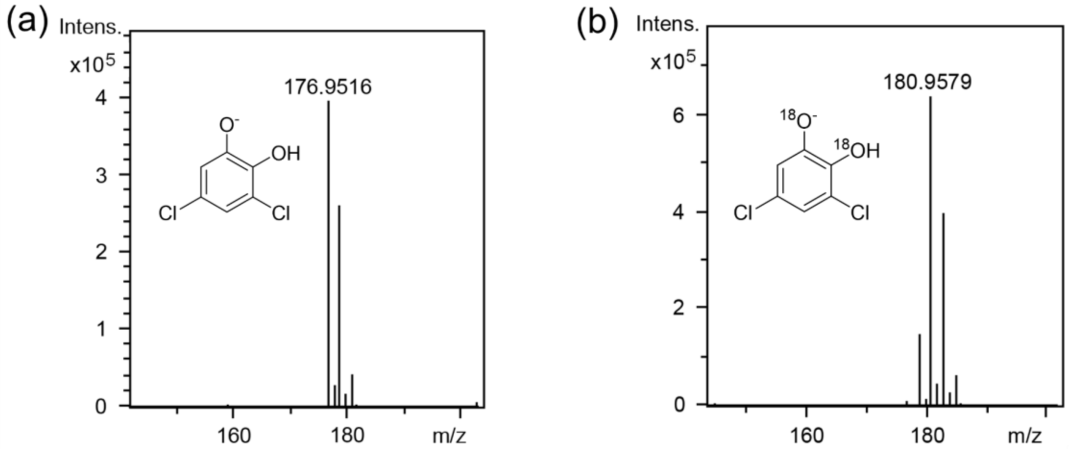


**Figure S85.** LC-MS analysis of **2** production from **1** by P450$_{BS\beta}$-F46A in $H_2O$ (**a**) or $H_2{}^{18}O$ (**b**). Reaction conditions: 1 μM P450$_{BS\beta}$-F46A, 500 μM **1**, 500 μM $H_2O_2$ as cofactor at 30 °C for 6 h. Upon comparing the $H_2{}^{18}O$ results of the standard **2** product (Figure S83b) with those from $H_2{}^{18}O$/$H_2O$ tracing experiments, it became clear that the hydroxyl oxygen atom of **2** production from **1** by P450$_{BS\beta}$-F46A exclusively derived from water.

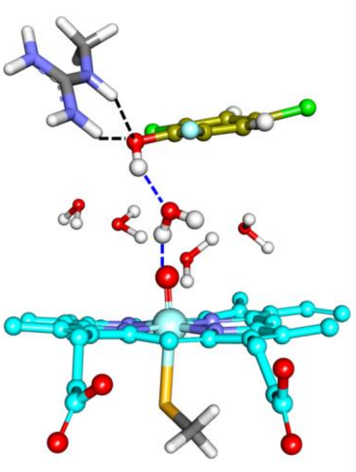

| | WAT No. | H-bond (%) |
|---|---|---|
| 6F-2,4-DCP: O atom | 8093 | 29.65 |
| | 23347 | 11.15 |
| | 6950 | 6.00 |
| | 12726 | 5.65 |
| | 5342 | 4.00 |
| | 16114 | 2.50 |
| | 2932 | 1.55 |
| | 1631 | 1.55 |
| | 6950 | 1.45 |
| | 16096 | 1.20 |

**Figure S86.** The numbering and statistical proportion of water molecules forming hydrogen bonds with the substrate and Cpd I. The solvent water molecules near Cpd I could easily replace the water molecule generated from $H_2O_2$ and make H-bond interactions with O1 of Cpd I and substrate 6F-2,4-DCP (**19**) during the MD simulation.

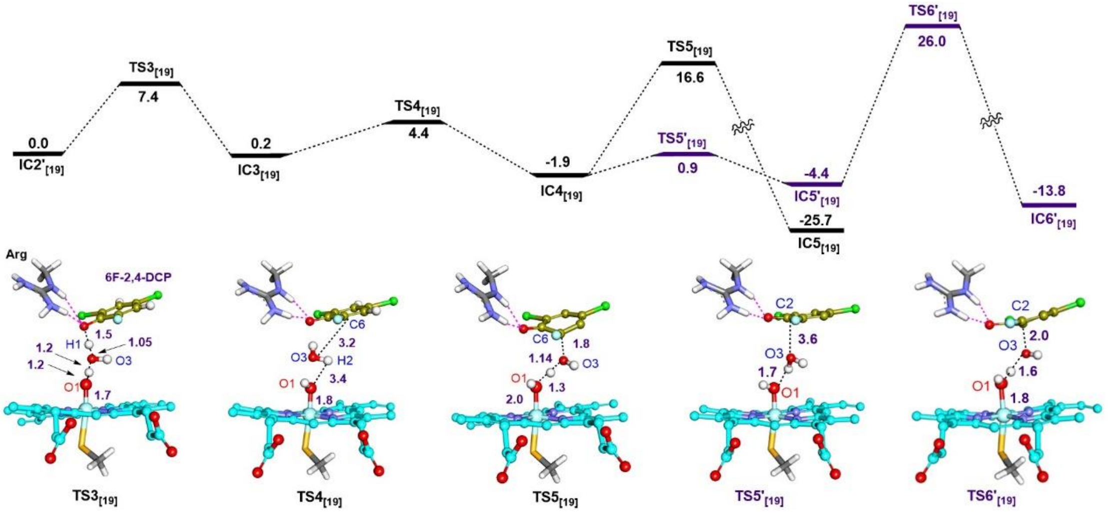


**Figure S87.** QM/MM calculated mechanism for the formation of F-substituted ene-ketone intermediate starting from Cpd I of $P450_{BS\beta}$-F46A, along with the QM/MM optimized structures of TSs species involved in the reactions. The energies are given in kcal/mol while the key distances are given in Å.

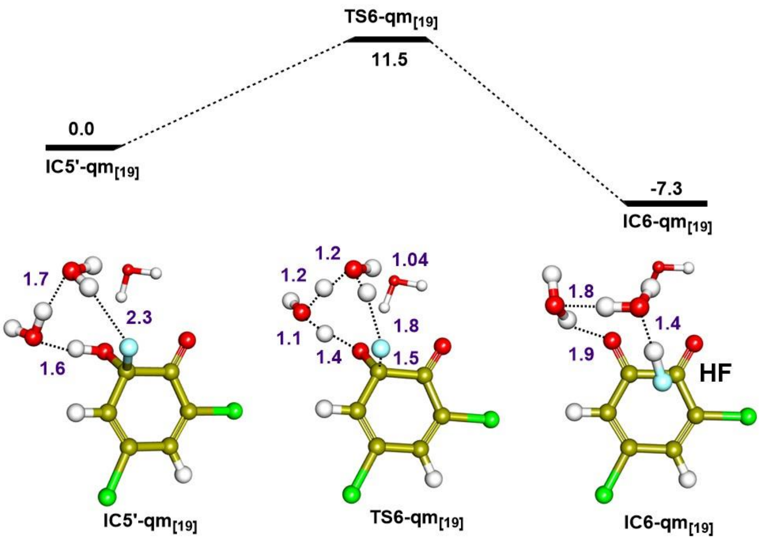


**Figure S88.** QM calculated potential energy profile (in kcal/mol) for the water-mediated proton transfer to generate the quinone intermediate (IC6-qm[**19**]) and HF.

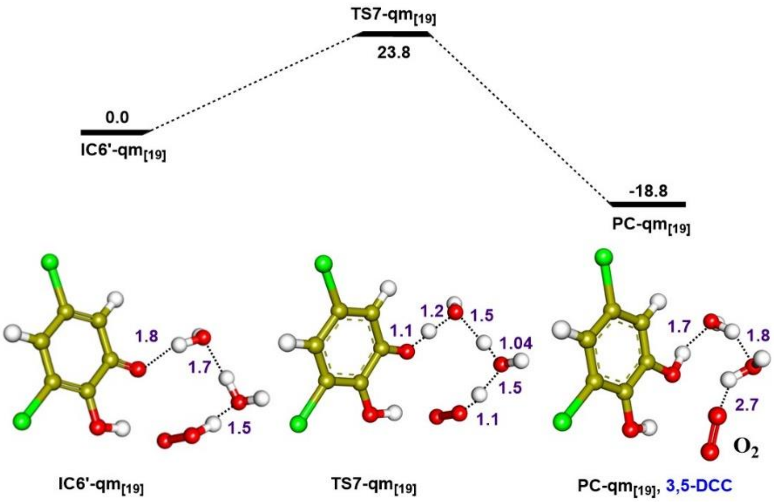


**Figure S89.** QM calculated potential energy profile (in kcal/mol) for the water-mediated hydrogen transfer from $H_2O_2$ to quinone for the generation of the product 3,5-DCC (PC-qm[**19**], **2**).

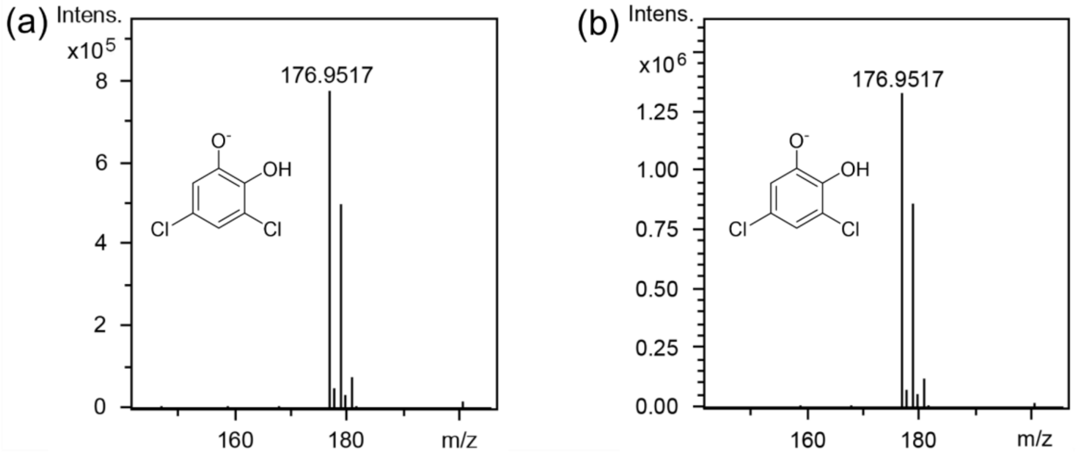


**Figure S90.** LC-MS analysis of **2** production from **19** by $P450_{BS\beta}$-F46A when supported by $H_2O_2$ (**a**) or $H_2{}^{18}O_2$ (**b**). Reaction conditions: 1 μM $P450_{BS\beta}$-F46A, 500 μM **19**, 500 μM $H_2O_2$ or $H_2{}^{18}O_2$ as cofactor at 30 °C for 6 h. These $H_2{}^{18}O_2/H_2O_2$ tracing experiments showed that the hydroxyl oxygen atom of **2** production from **19** by $P450_{BS\beta}$-F46A did not originate from hydrogen peroxide.

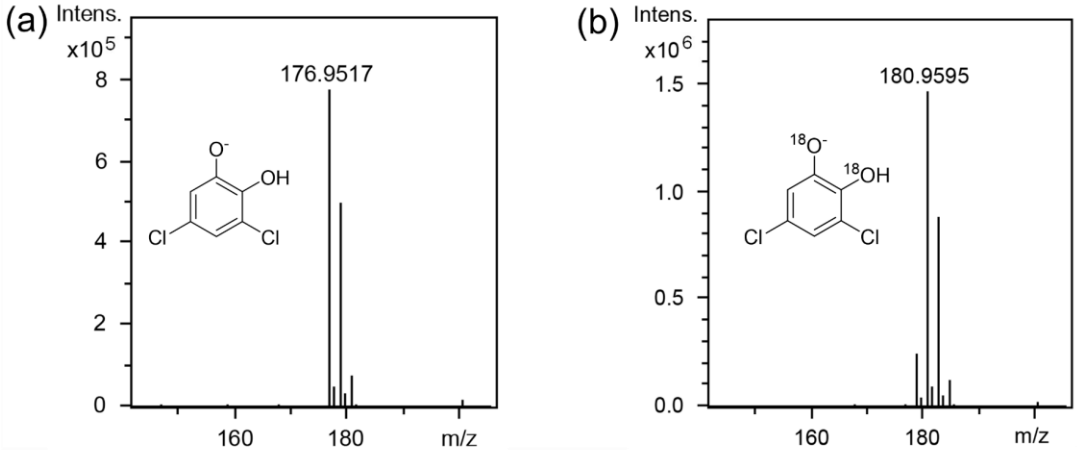


**Figure S91.** LC-MS analysis of **2** production from **19** by $P450_{BS\beta}$-F46A in $H_2O$ (**a**) or $H_2{}^{18}O$ (**b**). Reaction conditions: 1 μM $P450_{BS\beta}$-F46A, 500 μM **19**, 500 μM $H_2O_2$ as cofactor at 30 °C for 6 h. Upon comparing the $H_2{}^{18}O$ results of the standard **2** product (Figure S83b) with those from $H_2{}^{18}O/H_2O$ tracing experiments, it became clear that the hydroxyl oxygen atom of **2** production from **19** by $P450_{BS\beta}$-F46A exclusively derived from water.

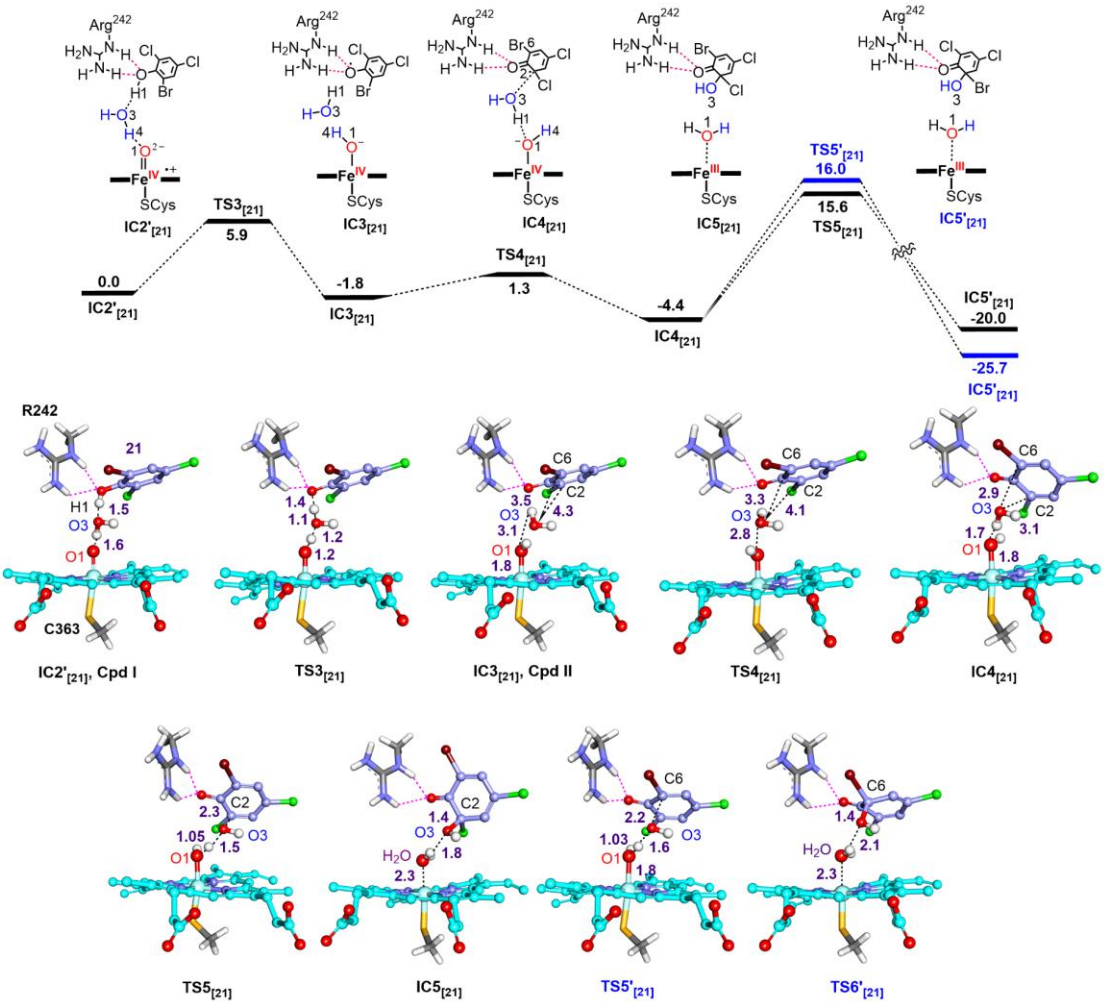

**Figure S92.** QM/MM calculated mechanism for the formation of hydroxylation intermediate starting from Cpd I of P450$_{BS\beta}$-F46A. The non-carboxylic substrate 2,4-dichloro-6-bromophenol (**21**) phenolic hydroxyl group interacting with the guanidyl group of Arg$^{242}$ is located above the heme, along with the QM/MM optimized structures of key species involved in the reactions. The key distances are given in angstrom (Å).

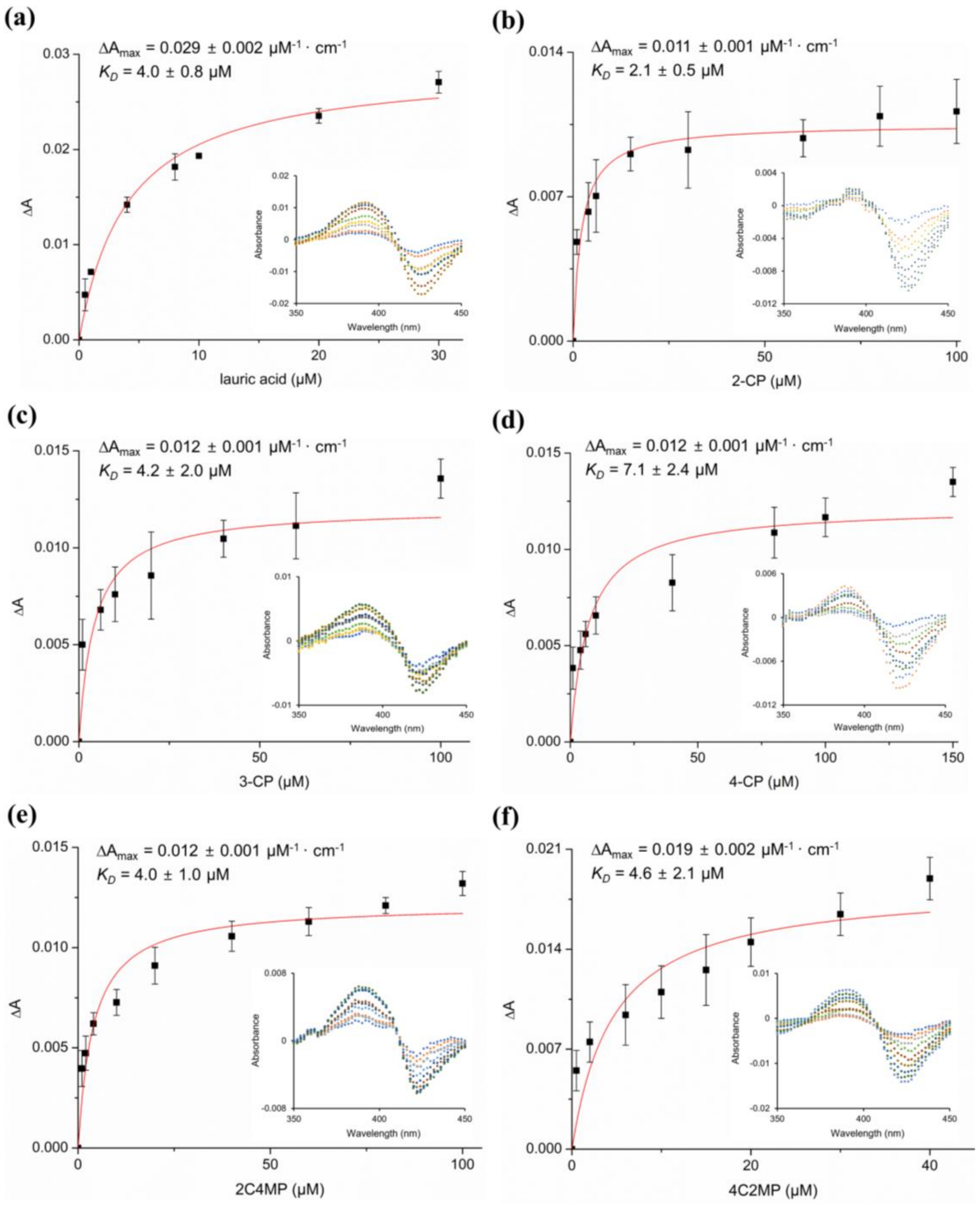


**Figure S93.** Substrate binding curves of P450$_{BS\beta}$-F46A. The substrate dissociation constant ($K_D$) measurements of lauric acid (**a**), 2-CP (**11**, **b**), 3-CP (**12**, **c**), 4-CP (**13**, **d**), 2C4MP (**14**, **e**), and 4C2MP (**15**, **f**) towards P450$_{BS\beta}$-F46A. The insets show the Type I difference spectra. All experiments were performed in triplicate.

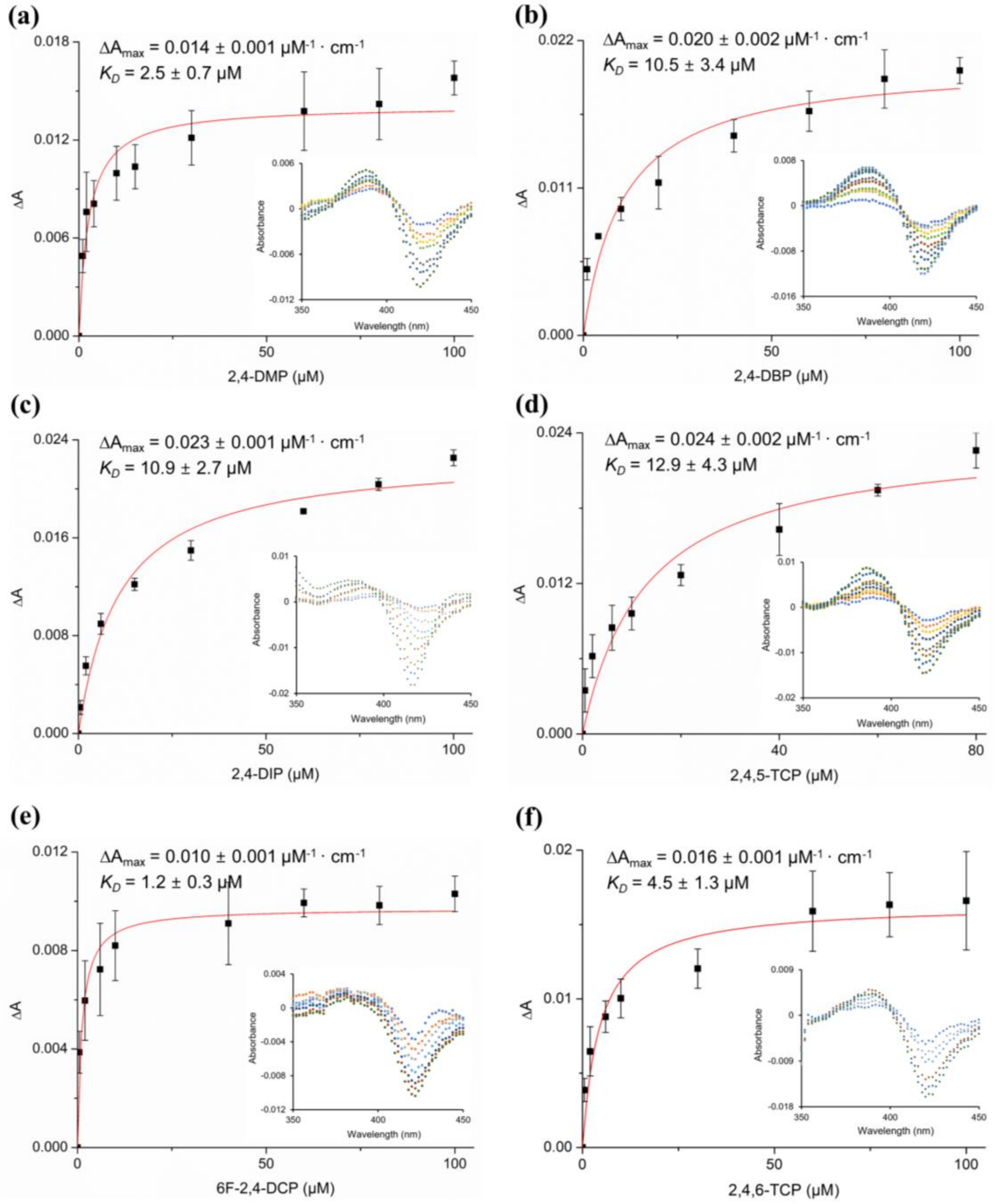


**Figure S94.** Substrate binding curves of P450$_{BS\beta}$-F46A. The substrate dissociation constant ($K_D$) measurements of 2,4-DMP (**5**, **a**), 2,4-DBP (**16**, **b**), 2,4-DIP (**17**, **c**), 2,4,5-TCP (**18**, **d**), 6F-2,4-DCP (**19**, **e**) and 2,4,6-TCP (**20**, **f**) towards P450$_{BS\beta}$-F46A. The insets show the Type I difference spectra. All experiments were performed in triplicate.

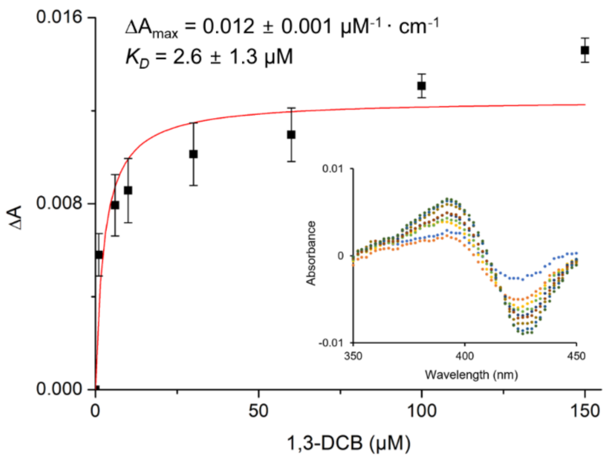


**Figure S95.** Substrate binding curve of P450$_{BSβ}$-F46A towards1,3-DCB (**7**). The inset shows the Type I difference spectra. All experiments were performed in triplicate.

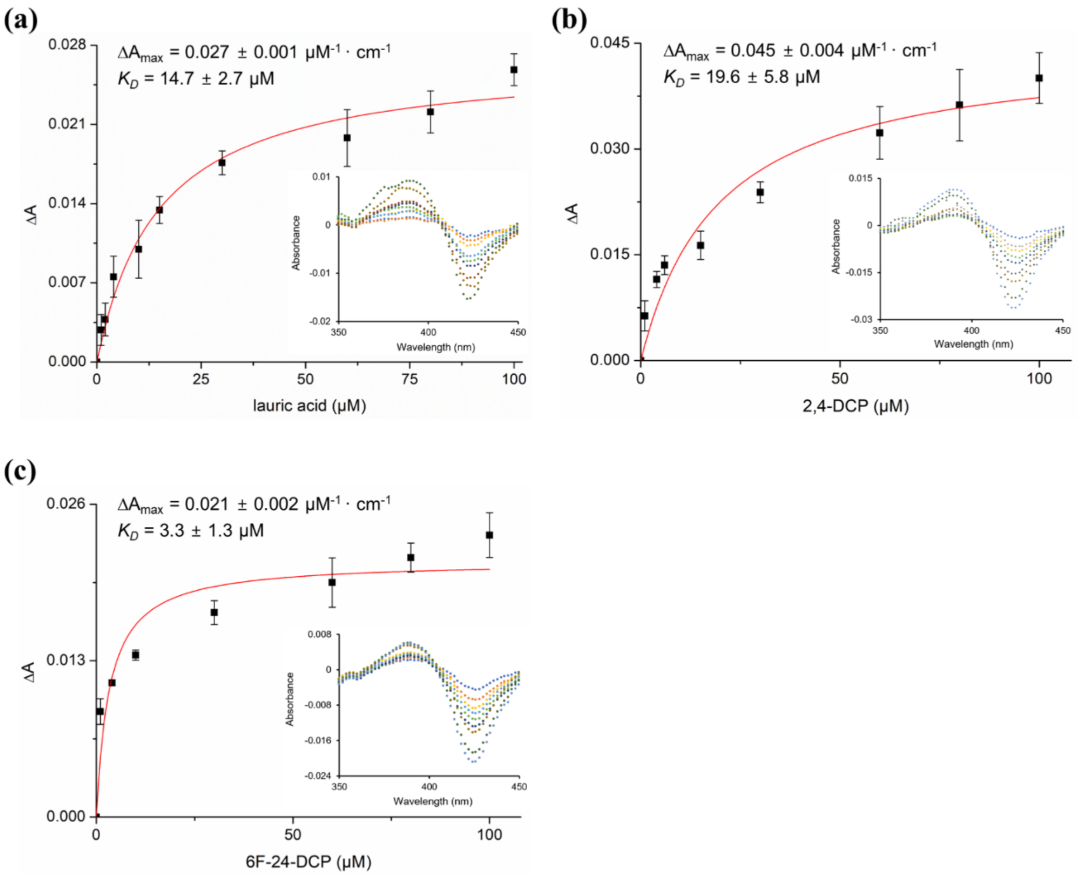


**Figure S96.** Substrate binding curves of P450$_{BSβ}$-F46A-R242A. The substrate dissociation constant ($K_D$) measurements of lauric acid (**a**), 2,4-DCP (**1**, **b**), and 6F-2,4-DCP (**19**, **c**) towards P450$_{BSβ}$-F46A-R242A. The insets show the Type I difference spectra. All experiments were performed in triplicate.
